# The recombination initiation functions DprA and RecFOR suppress microindel mutations in *Acinetobacter baylyi* ADP1

**DOI:** 10.1101/2023.11.22.568220

**Authors:** Mikkel M. Liljegren, João A. Gama, Pål J. Johnsen, Klaus Harms

**Affiliations:** Microbial Pharmacology and Population Biology Research Group, Department of Pharmacy, UiT The Arctic, University of Norway, Tromsø, Norway

## Abstract

Short-Patch Double Illegitimate Recombination (SPDIR) has been recently identified as a rare mutation mechanism. During SPDIR, ectopic DNA single-strands anneal with genomic DNA at microhomologies and get integrated in the course of DNA replication, presumably acting as Okazaki fragments. The resulting microindel mutations are highly variable in size and sequence. In the soil bacterium *Acinetobacter baylyi*, SPDIR mutations are tightly controlled by genome maintenance functions including RecA. In this study, we investigate the roles of DprA, RecFOR and RecBCD, which are cytoplasmic functions that load DNA single-strands with RecA. All three functions suppress SPDIR mutations in wildtype to levels below the detection limit. While SPDIR mutations are slightly elevated in the absence of DprA alone, they are strongly increased in the absence of both DprA and RecA. This SPDIR-avoiding function of DprA is not related to its role in natural transformation. These results suggest an antimutational function for DprA and offer an explanation for the ubiquity of *dprA* in the genomes of non-transformable bacteria.

## Introduction

In clonally reproducing organisms, faithful replication of the genetic material is necessary for stable inheritance of evolutionary adapted traits. However, replication is imperfect and may result in spontaneous mutations (Echols and Goodman, 1991; Kunkel and Bebenek, 2000). Mutations alter the genetic content and occur at low but quantifiable specific, evolutionary optimized frequencies (Denamur and Matic, 2006). Mutations, together with recombination, generate genetic diversity that is the substrate for evolutionary forces such as selection or drift (Didelot and Maiden, 2010; Hershberg, 2015; King, 1972).

Recently, Short-Patch Double Illegitimate Recombination (SPDIR) has been identified as a rare DNA mutation mechanism (Harms et al., 2016). It is thought that during SPDIR, single-stranded (ss) DNA molecules from intra- or extragenomic sources anneal at exposed ssDNA stretches in genomic DNA (such as replication forks) and get integrated into a nascent strand during DNA replication (Harms et al., 2016), acting effectively as primers for Okazaki fragments (Figure 1). Spontaneous annealing of fully heterologous DNA strands can occur at microhomologies, or at extended microhomologies that can contain mismatches and gaps, and can consequently result in small (three to approximately 100 bp) insertion-deletion (microindel) patterns of highly variable sequence (Harms et al., 2016).

**Figure 1:**
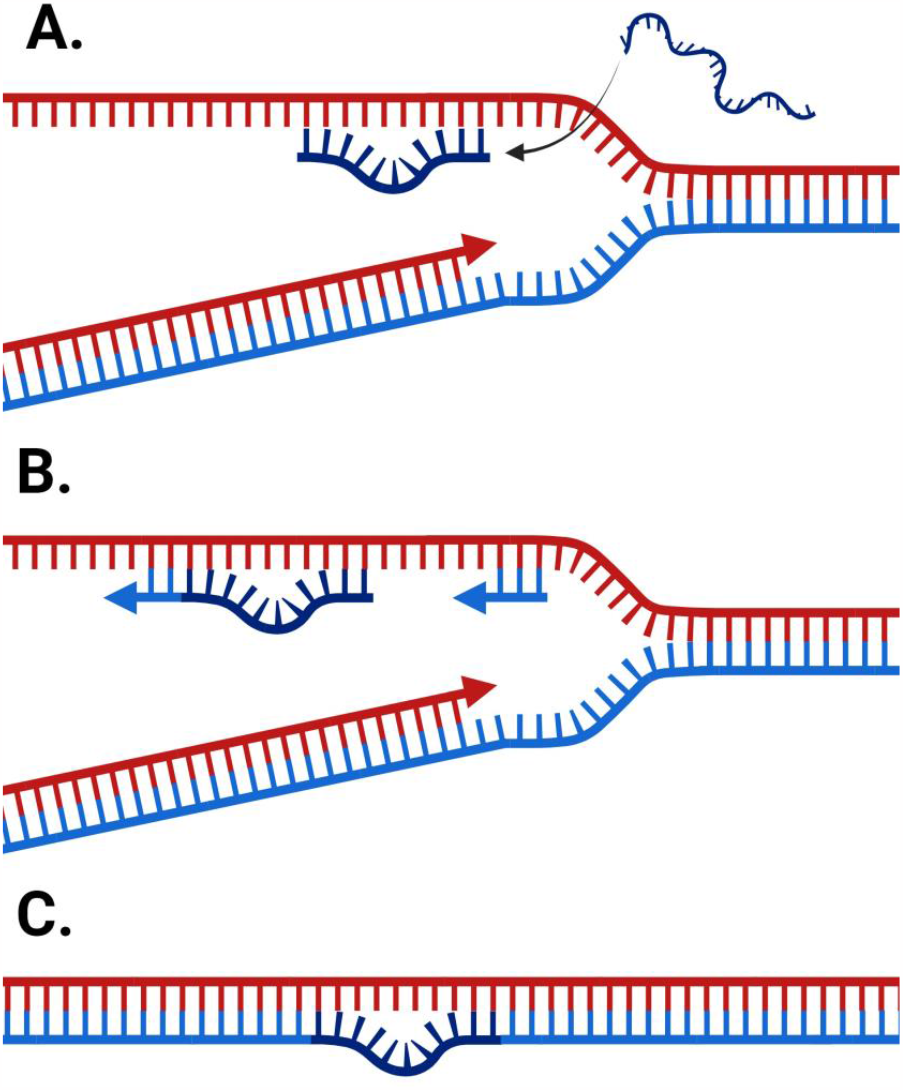
Formation of SPDIR mutations. A: A DNA single-strand can anneal with an exposed lagging strand at a microhomology during DNA replication. B: The strand is processed, extended, and subsequently integrated into the newly replicated strand. C: The result is a heteroduplex that is fixed in the genome after a subsequent round of replication. Created with BioRender.com.

In living organisms, ssDNA molecules are a DNA damage signal, and their level is generally tightly controlled by genome maintenance functions (Maslowska et al., 2019). In the gammaproteobacterial model organism *Acinetobacter baylyi*, it has been found that SPDIR mutations are very rare but detectable at above 10^-13^ per cell and locus (Harms et al., 2016). Remarkably, experimental studies demonstrated that this frequency can increase by orders of magnitude under genotoxic stress, in genome-maintenance mutants, or in the course of natural transformation. Under such conditions, ssDNA levels are elevated due to DNA damages and/or repair activity, lack of genomic maintenance, or by active DNA uptake (Johnston et al., 2014; Walker, 1984). Consequently, SPDIR mutations can play a role in adaptation under stress and evolution in general(Harms et al., 2016).

The quantification of mutation events is essential for their experimental characterization. In *A. baylyi*, SPDIR mutations can be conveniently detected and quantified using the *hisC*::’ND5i’ fusion allele (Harms et al., 2016; Overballe-Petersen et al., 2013). In this allele, the gene encoding histidinol phosphate aminotransferase has been rendered nonfunctional through a 5’-fusion in frame with a 128-bp DNA segment of coding DNA interrupted by two adjacent stop codons (Figure 2B). A cell harbouring *hisC*::’ND5i’ can grow on rich medium but cannot form a colony on minimal medium. When the two stop codons are mutationally removed or bypassed, the resulting *hisC*^+^ mutant allele can be expressed, resulting in colony formation on minimal medium (Figure 2A). Such mutations are typically deletions (i.e., single illegitimate recombination events), occasionally SPDIR events (Figure 2C), and rarely other mutation types. Notably, single nucleotide changes are effectively undetectable using the *hisC*::’ND5i’ construct.

**Figure 2:**
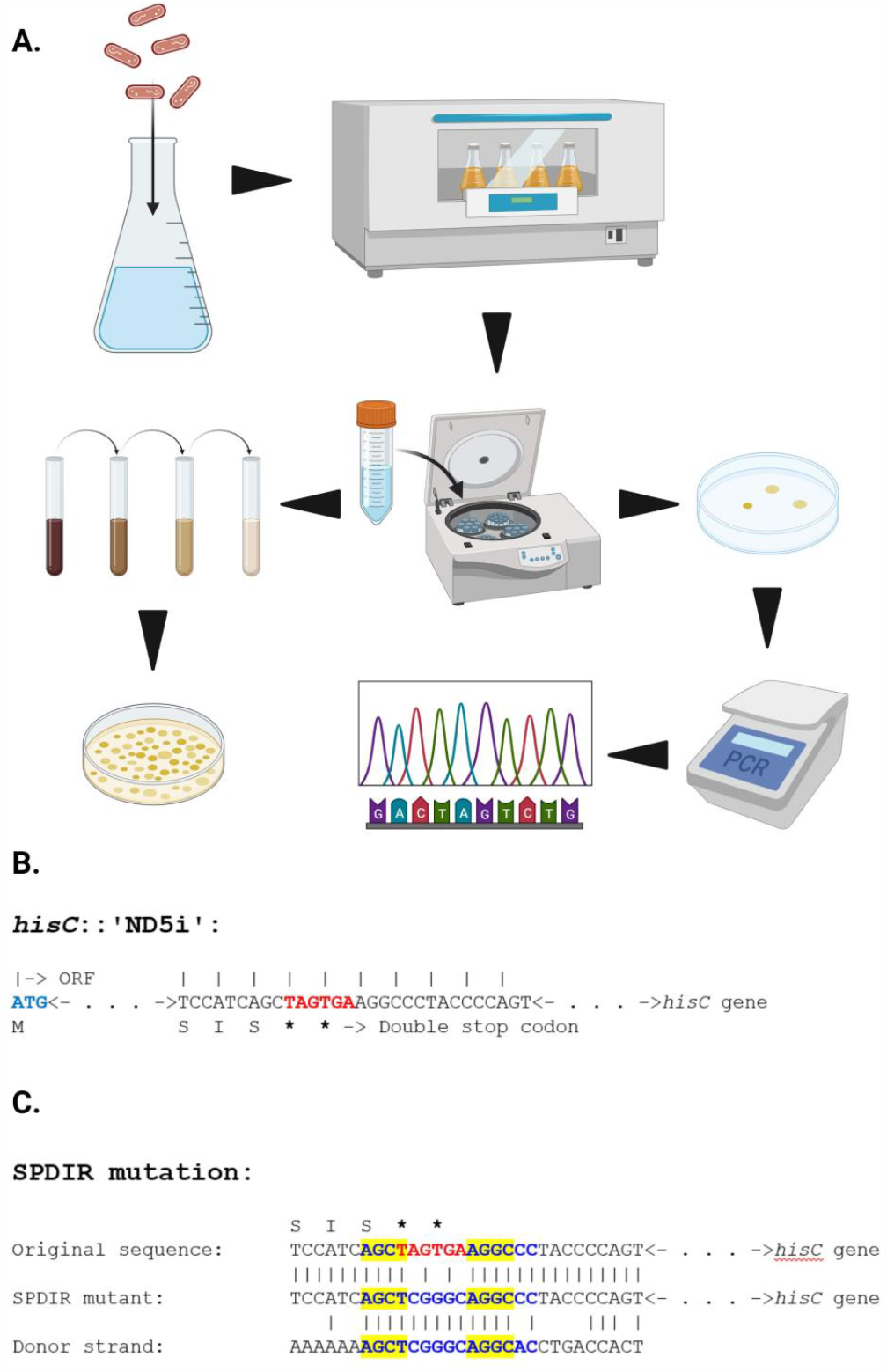
A: Experimental flowchart. His^-^ strains are grown non-selectively, washed, and plated on rich medium (in appropriate dilution) for CFU, and on minimal medium to screen for mutant titers. His^+^ colonies are isolated and identified by PCR and sequencing of the *hisC* allele. B: Sequence detail of the ‘ND5i’ insert with the sequential stop codons marked in red. C: Example SPDIR mutation, displayed as triple alignment of the original DNA sequence, the mutant sequence and the mutagenic, heterologous DNA donor strand. The double stop codon is indicated in red, the extended microhomology is indicated in blue, and the crossover sites are highlighted in yellow. Created with BioRender.com.

RecA was previously identified as a central function (in cooperation with ssDNA exonucleases) in avoiding SPDIR mutations in wildtype cells (Harms et al., 2016). RecA binds to ssDNA through replacement of ssDNA Binding Protein (SSB) (Bell et al., 2012), and it was hypothesized that RecA-bound ssDNA could not hybridize, and therefore SPDIR mutations were kept close to the limit of detection in wildtype. In addition, RecA is the key function in homologous recombination and therefore a central genome maintenance function (Bell and Kowalczykowski, 2016; Clark, 1973; Michel and Leach, 2012; Roca et al., 1990). Without RecA, recombinational repair of DNA damages is impaired, and the cell becomes hypersensitive to DNA damages (Clark, 1973). ssDNA can bind to RecA spontaneously, however, this process is inefficient and impeded by SSB (Bell et al., 2012; Bell and Kowalczykowski, 2016). In the cell, binding of RecA to ssDNA is facilitated by recombination initiation functions that can actively load RecA molecules on ssDNA (Cox, 2007), displacing SSB and in turn initiating homologous recombination. These recombination initiation functions include: RecBCD (Anderson and Kowalczykowski, 1997), RecFOR/RecOR (Sakai and Cox, 2009), and, during natural transformation, DprA (Mirouze et al., 2013; Mortier-Barrière et al., 2007; Quevillon-Cheruel et al., 2012; Yadav et al., 2013).

In this study, we experimentally investigated the role of the major RecA-loading functions on the containment and frequency of SPDIR mutations by using targeted knockout mutants of *A. baylyi*. DprA was studied with respect to absence and presence of RecA and to natural transformation, while the roles of RecFOR and RecBCD were explored regarding their DNA repair activities.

## Results

### *DprA suppresses SPDIR mutations in concert with RecA*. We created a *dprA* deletion mutant of the wildtype

*A. baylyi* strain that carried the *hisC*::’ND5i’ fusion allele and quantified its spontaneous His^+^ mutation frequency. From altogether 24 independently grown overnight cultures, we recovered 28 non-sibling His^+^ revertants. Sanger sequencing of the *hisC* allele revealed that one His^+^ clone resulted from a SPDIR mutation, corresponding to a calculated SPDIR frequency of 2.3×10^-12^ (Table 1; all SPDIR mutations found in this study are listed in supplemental file S1). In control experiments with the wildtype, no SPDIR mutations were detected among a total of 39 non-sibling His^+^ clones from 34 independent cultures (Table 1; detection limit: 1.8×10^-12^), confirming that SPDIR mutations in wildtype cells occur below the detection limit regardless of DprA.

**Table 1:** Impact of DprA and DprA/RecA-interactions on His^+^ and SPDIR frequency.

| 1. *baylyi* | Median | SPDIR | Calculated | Number of | CFU |
| --- | --- | --- | --- | --- | --- |
| genotype | His<sup>+</sup> | events | SPDIR | experiments | plated |
|  | frequency | per His<sup>+</sup> | frequency |  |  |
|  |  | event |  |  |  |
| *ΔdprA* | 6.4×10<sup>-11</sup> | 3.6% (1/28) | 2.3×10<sup>-12</sup> | 24 | 6.8×10<sup>12</sup> |
| wildtype (*dprA*<sup>+</sup>) | 5.7×10<sup>-11</sup> | <2.5% (0/39) | <1.8×10<sup>-12</sup> | 34 | 7.3×10<sup>12</sup> |
| *ΔdprA ΔrecA* | 2.2×10<sup>-10</sup> | 34% (12/35) | 7.6×10<sup>-11</sup> | 12 | 3.5×10<sup>12</sup> |
| *dprA*<sup>+</sup> *ΔrecA* | 7.1×10<sup>-11</sup> | 12% (3/25) | 8.6×10<sup>-12</sup> | 20 | 3.4×10<sup>12</sup> |
| *ΔdprA ΔrecJ ΔexoX* | 1.1×10<sup>-10</sup> | 43% (18/42) | 4.7×10<sup>-11</sup> | 12 | 4.2×10<sup>12</sup> |
| *dprA*<sup>+</sup> *ΔrecJ ΔexoX* | 2.5×10<sup>-10</sup> | 32% (43/135) | 8.0×10<sup>-11</sup> | 39 | 9.8×10<sup>12</sup> |
| *ΔdprA ΔrecA ΔrecJ ΔexoX* | 2.3×10<sup>-9</sup> | 86% (73/85) | 2.0×10<sup>-9</sup> | 11 | 2.9×10<sup>12</sup> |
| *dprA*<sup>+</sup> *ΔrecA ΔrecJ ΔexoX* | 1.8×10<sup>-9</sup> | 86% (60/70) | 1.5×10<sup>-9</sup> | 8 | 1.3×10<sup>12</sup> |

Previous work revealed a quantifiable increase in SPDIR mutation frequencies in a Δ*recA* mutant of *A. baylyi* (Harms et al., 2016). We deleted *dprA* in an *A. baylyi* Δ*recA* strain and observed that 34% of all non-sibling His^+^ mutations were caused by SPDIR, giving a SPDIR frequency of 7.6×10^-11^ (Table 1). In contrast, in a *dprA*^*+*^ Δ*recA* strain the calculated SPDIR frequency was approximately ninefold lower (8.6×10^-12^), and only 12% of the His^+^ events were caused by SPDIR mutations (Table 1), confirming our previous findings (Harms et al., 2016). We conclude that in concert DprA and RecA suppress the occurrence of SPDIR mutations in *A. baylyi* wildtype cells. Absence of DprA alone has a small but detectable effect on occurrence of SPDIR mutations relatively to wildtype (Table 1), comparable with the single RecA, RecJ, or ExoX deficiencies reported previously (Harms et al., 2016).

We further repeated the fluctuation experiments in a strain deficient for the ssDNA-specific exonucleases RecJ and ExoX that is prone to increased SPDIR frequencies (Harms et al., 2016) and facilitates the quantification of SPDIR events. In a Δ*dprA* Δ*recJ* Δ*exoX* strain, the calculated SPDIR frequency was 4.7×10^-11^ and was similar to that of the *dprA*^+^ Δ*recJ* Δ*exoX* strain (8.0×10^-11^; Table 1). Thus, DprA does not have a strong effect on SPDIR mutations in Δ*recJ* Δ*exoX* mutants. Overall, the combined deletions of the RecJ and ExoX ssDNA exonucleases have a similar effect on His^+^ and SPDIR frequencies as the Δ*dprA* Δ*recA* double mutation (Table 1).

We also investigated the effect of combined deletions of DprA, RecA and the ssDNA exonucleases. In the Δ*dprA* Δ*recA* Δ*recJ* Δ*exoX* quadruple mutant, the SPDIR frequency was 2.0×10^-9^, which was ≥850-fold higher compared to the Δ*dprA* single mutant. 86% of all His^+^ mutations in the quadruple mutant were caused by SPDIR (Table 1). However, the results were similar to those obtained with a *drpA*^+^ Δ*recA* Δ*recJ* Δ*exoX* strain (1.5×10^-9^; 86% SPDIR mutations; Table 1). These results confirm that the genome maintenance functions RecA, RecJ and ExoX together suppress SPDIR mutations in wildtype by up to three orders of magnitude, regardless of DprA.

*DprA does not modulate SPDIR formation with foreign DNA. A. baylyi* is naturally transformable (Metzgar et al., 2004), and during transformation DprA is thought to bind to taken-up ssDNA and facilitate subsequent recombination through RecA-loading (Sharma et al., 2023; Yadav et al., 2013). Our previous study has demonstrated that supplementation of the overnight culture with DNA from any source increased SPDIR frequencies (Harms et al., 2016), and it is conceivable that DprA modulates SPDIR mutation frequencies by binding to taken-up DNA. Low amounts of isogenic DNA also accumulate in growing cultures due to cell lysis, however, taken-up DNA cannot be distinguished from intracellular DNA as source for SPDIR mutations. We exposed the Δ*dprA* Δ*recJ* Δ*exoX* mutant to foreign DNA (purified from *Bacillus subtilis* 168) in the overnight cultures and obtained a median His^+^ frequency of 5.8×10^-10^ (Table 2) which was ≥six-fold higher than the frequency without DNA (9.8×10^-11^; Table 1). Supplementation with foreign DNA increased the proportion of SPDIR mutations among all non-sibling His^+^ mutations from 41% (Table 1) to 70% (Table 2). The *B. subtilis* DNA served as template for 12 of the 39 independent SPDIR mutants recovered. When repeating the transformation experiments in a *dprA*^+^ Δ*recJ* Δ*exoX* strain (Table 2), the SPDIR frequency (5.5×10^-10^) was similar to that of the Δ*dprA* Δ*recJ* Δ*exoX* strain, but the proportion of SPDIR events among His^+^ events was lower (51%) with eight out of 24 SPDIR mutations resulting from *B. subtilis* DNA (Table 2).

**Table 2:** Impact of transformation and competence on DprA-mediated SPDIR suppression.

| Experimental setup /<br>genetic background | Results | *A. baylyi* strain |  |
| --- | --- | --- | --- |
| | | $\Delta dprA \Delta recJ \Delta exoX$ | $dprA^+ \Delta recJ \Delta exoX^a$ |
| + *B. subtilis* DNA | median His<sup>+</sup> frequency | $5.8 \times 10^{-10}$ | $5.5 \times 10^{-10}$ |
|  | SPDIR fraction | 70% (39/56) | 51% (24/47) |
|  | SPDIR with *B. subtilis* DNA | 21% (12/56) | 17% (8/47) |
| | calculated SPDIR frequency | $4.0 \times 10^{-10}$ | $2.8 \times 10^{-10}$ |
|  | number of experiments | 10 | 9 |
| $\Delta comA$ | median His<sup>+</sup> frequency | $1.4 \times 10^{-10}$ | $6.5 \times 10^{-11}$ |
|  | SPDIR fraction | 27% (4/15) | 35% (8/23) |
| | calculated SPDIR frequency | $3.7 \times 10^{-11}$ | $2.3 \times 10^{-11}$ |
|  | number of experiments | 11 | 10 |

a: data for this strain taken from Harms et al., 2016.

To rule out any effect of horizontal gene transfer in our fluctuation assays, we deleted the *comA* gene encoding the DNA uptake pore essential for natural transformation (Overballe-Petersen et al., 2013). Both Δ*dprA* Δ*comA* Δ*recJ* Δ*exoX* and *dprA*^*+*^ Δ*comA* Δ*recJ* Δ*exoX* mutants displayed identical SPDIR frequencies (3.7×10^-11^ and 2.3×10^-11^, respectively; Table 2). Altogether, the results confirm that extracellular DNA can boost SPDIR mutations through natural transformation, however, the increase is largely independent of DprA. Notably, deletion of DprA does not block transformation, as demonstrated here through illegitimate recombination.

*RecFOR and RecBCD suppress SPDIR mutations*. In bacteria, RecA is actively loaded with ssDNA by DprA to initiate homologous recombination during natural transformation (Mirouze et al., 2013; Sharma et al., 2023; Yadav et al., 2013). In addition, the the RecFOR proteins as well as the RecBCD enzyme can load ssDNA with RecA to initiate homologous recombination and recombinational DNA repair. While RecFOR mainly initiates gap repair (Morimatsu and Kowalczykowski, 2003) and restores stalled and collapsed replication forks during DNA replication (Michel et al., 2001; Morimatsu et al., 2012), RecBCD is crucial for repairing double-strand DNA breaks (Dillingham and Kowalczykowski, 2008; Kuzminov, 1996; White et al., 2018). In many organisms, DNA strand breaks are processed towards the RecBCD DNA repair pathway, and in turn RecBCD deficiency is highly detrimental to those organisms when the resulting intermediates cannot be processed by RecFOR or by alternative recombinases (Bidnenko et al., 1999; Harms and Wackernagel, 2008; Courcelle et al., 2015). Here we extend the previous hypothesis that the frequency of SPDIR mutations increase due to induced DNA damages (Harms et al., 2016) by evaluating the effect of deleting the *recF, recO* and *recR* genes, as well as the *recBCD* operon, of *A. baylyi*. In a Δ*recFOR* mutant, the calculated SPDIR frequency was 6.6×10^-12^, and two SPDIR mutations were recovered among 30 non-sibling His^+^ mutations (Table 3). Using a Δ*recFOR* Δ*recJ* Δ*exoX* mutant, the calculated SPDIR frequency was 1.8×10^-10^ (Table 3) which was twofold higher than that of the Δ*recJ* Δ*exoX* strain (Table 1), and ≥27-fold higher than the Δ*recFOR* strain. The proportion of SPDIR mutations among His^+^ events in Δ*recFOR* Δ*recJ* Δ*exoX* was 46% (35% in Δ*recJ* Δ*exoX* and 6.7% in Δ*recFOR*). We conclude that the RecFOR DNA repair functions contribute to avoiding SPDIR mutations in wildtype cells.

**Table 3:**
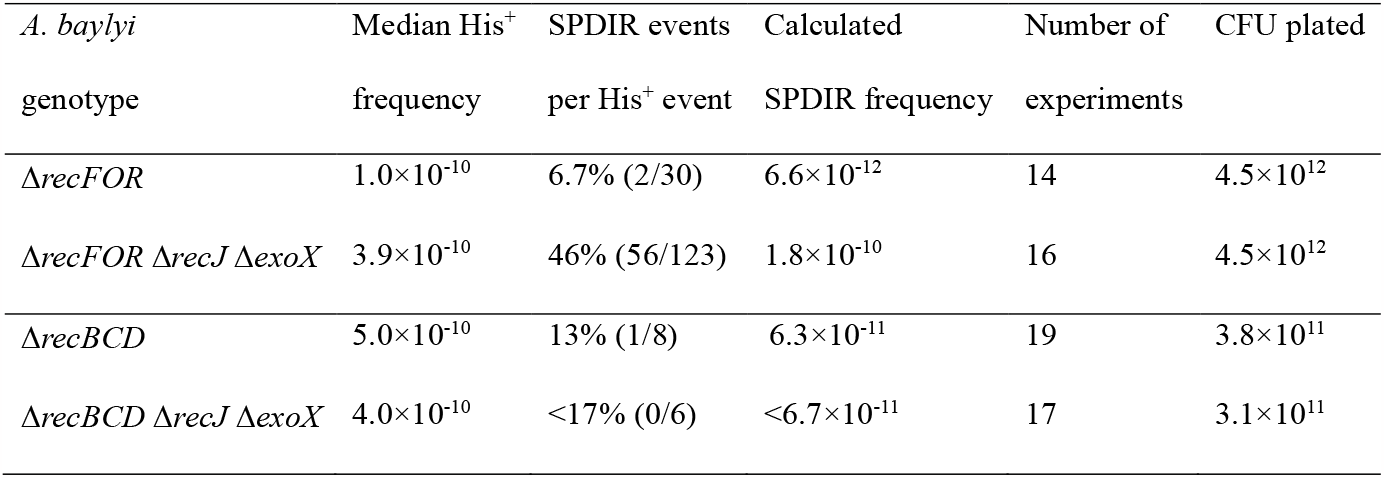
Impact of RecFOR and RecBCD on His^+^ and SPDIR frequency.

| *A. baylyi*<br>genotype | Median His<sup>+</sup><br>frequency | SPDIR events<br>per His<sup>+</sup> event | Calculated<br>SPDIR frequency | Number of<br>experiments | CFU plated |
| --- | --- | --- | --- | --- | --- |
| *ΔrecFOR* | 1.0×10<sup>-10</sup> | 6.7% (2/30) | 6.6×10<sup>-12</sup> | 14 | 4.5×10<sup>12</sup> |
| *ΔrecFOR ΔrecJ ΔexoX* | 3.9×10<sup>-10</sup> | 46% (56/123) | 1.8×10<sup>-10</sup> | 16 | 4.5×10<sup>12</sup> |
| *ΔrecBCD* | 5.0×10<sup>-10</sup> | 13% (1/8) | 6.3×10<sup>-11</sup> | 19 | 3.8×10<sup>11</sup> |
| *ΔrecBCD ΔrecJ ΔexoX* | 4.0×10<sup>-10</sup> | <17% (0/6) | <6.7×10<sup>-11</sup> | 17 | 3.1×10<sup>11</sup> |

The Δ*recBCD* mutant displayed a single SPDIR event among eight His^+^ clones, with a calculated SPDIR frequency of 6.3×10^-11^ (Table 3). Against our expectations, no SPDIR events were detected among the six recovered His^+^ clones of the Δ*recBCD* Δ*recJ* Δ*exoX* triple mutant (Table 3). All RecBCD-deficient strains displayed poor viability (<10% CFU per cell in the overnight culture, determined with a hemocytometer), growing slowly and mostly as filaments, and forming small colonies, often with irregular morphology.

Altogether, these results indicate a role for RecFOR and RecBCD in SPDIR avoidance in wildtype. Furthermore, RecBCD deficiency appears to suppress SPDIR mutations in the absence of RecJ and ExoX.

## Discussion

The *dprA* gene is essential to maintain a fully transformation-proficient phenotype in naturally transformable organisms (Mortier-Barrière et al., 2007). Despite this association with transformability, *dprA* is nearly ubiquitous in bacteria, including those not thought to be competent for transformation, and in these organisms *dprA* displays no discernible phenotype (Johnston et al., 2014; Smeets et al., 2006). Why, then, is DprA so ubiquitous and maintained even in naturally non-transformable bacteria? This enigma may be resolved by our observation that DprA acts as a component to contain SPDIR frequencies to a minimum through its ssDNA-binding and RecA-loading functions, regardless of natural transformation. This SPDIR-avoiding activity of DprA on intragenomic DNA may be universal. SPDIR mutations are thought to be mostly detrimental (Harms et al., 2016), and their avoidance may select for the continued maintenance of *dprA* in the bacterial genome even in the absence of natural competence.

In the process of natural transformation, DprA binds to taken-up ssDNA molecules and loads them with RecA protein for initiation of homologous recombination (Sharma et al., 2023). Lack of DprA alone prevents formation of the nucleoprotein filament and therefore leads to decrease of homologous recombination across transformable species (Huang et al., 2019; Hülter et al., 2017; Ithurbide et al., 2020; Mirouze et al., 2013). The failure of ssDNA molecules to get charged with RecA can explain the increased SPDIR frequencies observed. Indeed, in our experiments the Δ*dprA* and Δ*recA* mutants result in elevated SPDIR mutation frequencies, and their simultaneous deletion led to an increase higher than their individual effects (Table 1). In the absence of both DprA and RecA, ssDNA molecules (presumably bound to SSB) could interact more freely with single-stranded genomic DNA regions at replication forks or gaps, which can explain the higher SPDIR frequency as well as the increased SPDIR proportion among all His^+^ mutations (Table 1). In the Δ*recA* strain, the SPDIR frequency was approximately four times higher than in the Δ*dprA* strain (Table 1), implying that the majority of ssDNA molecules are not loaded with RecA by DprA. We speculate that either RecFOR, or spontaneous association of RecA, are responsible for this observation.

In this context, the effects of the *dprA* and *recA* deletions in a Δ*recJ* Δ*exoX* mutant are interesting. In *A. baylyi*, the latter two genes encode the two exonucleases specific for 5’- and 3’-ssDNA ends (RecJ and ExoX, respectively). In their combined absence, ssDNA molecules are prevented from quick degradation (Overballe-Petersen et al., 2013) which strongly increases the occurrence of SPDIR (Harms et al., 2016). Simultaneous deficiency of RecA, RecJ and ExoX increases the SPDIR frequency by at least three orders of magnitude (Table 1), exceeding the frequency of single-nucleotide changes (‘point mutations’) (Harms et al., 2016). This effect likely results from the combination of increased survival of ssDNA (lack of exonucleases) and increased hybridization (lack of RecA). The result is an increase in the pool of ssDNA as a substrate for SPDIR events. Notably, further lack of DprA appears to have no additional effect, in agreement with the earlier findings in other bacterial species that DprA protects ssDNA from exonuclease activity (Mortier-Barrière et al., 2007), a function that would be rendered irrelevant in exonuclease-deficient strains. In the absence of DprA and RecA, cytoplasmic ssDNA remains bound solely to SSB. However, SSB is an essential function (De Berardinis et al., 2008) making it difficult to study *in vivo* to which degree the SSB-ssDNA-complex resists exonuclease activity or generally inhibits SPDIR mutations compared to DprA and RecA.

In addition to DprA, we also identified a role in SPDIR suppression for the RecFOR proteins. We hypothesize that in the absence of the RecFOR proteins both the repair of gapped genomic DNA and the re-establishment of collapsed replication forks are impaired (Michel et al., 2007; Morimatsu and Kowalczykowski, 2003), and thus ssDNA intermediates can engage in SPDIR. This is supported by the Δ*recFOR* Δ*recJ* Δ*exoX* mutant where the ratio of SPDIR events among His^+^ clones, as well as an increased overall SPDIR frequency (Table 3), presumably due to the lack of degradation of free ssDNA ends.

The results with RecBCD should be critically evaluated considering the low viability of Δ*recBCD* mutants. RecBCD-deficient cells typically grow as filaments, and approximately only one in ten cells form a colony (Kickstein et al., 2007). The additional deletion of *recJ* and *exoX* further reduced viability to ≥8% (Table 3). The fact that we encountered a single SPDIR mutant among the rare His^+^ revertants indicated that RecBCD does have a role in SPDIR avoidance in *A. baylyi* similar to RecFOR. In a Δ*recBCD* Δ*recJ* Δ*exoX* mutant, however, no SPDIR mutants were found. This contrasts with all other experiments using a Δ*recJ* Δ*exoX* strain to date where SPDIR mutations make up for ≥27% of all His^+^ mutants. We hypothesize that DNA repair in general is impaired in the Δ*recBCD* Δ*recJ* Δ*exoX* strain, and the rare colony formers are possibly individual cells that have suffered very few or even no DNA damages. Such damages or their repair intermediates could be the source for the ssDNA causing SPDIR mutations in wildtype cells. The viability of a Δ*recBCD* Δ*recJ*

Δ*exoX* mutant was surprising in the first place, since a Δ*recBCD* Δ*recJ* mutant is thought to be non-viable (Kickstein et al., 2007). Conceivably, the 3’-ssDNA exonuclease ExoX in Δ*recBCD* Δ*recJ* destroys potential recombinogenic DNA 3’-termini. In a Δ*recBCD* Δ*recJ* Δ*exoX* strain these intermediates are possibly retained and could be processed by RecFOR, however, in the absence of RecJ initiation of RecFOR-mediated repair would be hampered, explaining the low viability phenotype. Thus, ExoX in *A. baylyi* would be the functional counterpart of the 3’-ssDNA exonuclease SbcB (<u>s</u>uppression of Rec<u>BC</u> mutations) in *Escherichia coli recBCD* mutants (Bidnenko et al., 1999).

In the experiments conducted for this study, we found no SPDIR mutant in wildtype cells (detection limit: <1.8×10^-12^; Table 1), but in our previous publication we recovered two SPDIR mutants, and based on those we calculated a SPDIR frequency of 5.5×10^-13^ (Harms et al., 2016). Since spontaneous mutations occur at random, the true biological SPDIR frequency for *A. baylyi* is likely below 10^-12^, probably somewhat above 10^-13^. How are SPDIR frequencies in other organisms under benign conditions? Based on our data we cannot predict. While annealing of DNA single-strands at (extended) microhomologies under physiological conditions is a physical phenomenon applicable to every biological entity using DNA as genetic material (Villarreal et al., 2012; Wetmur, 2006), we find that a set of gene products associated with genome maintenance (*recJ*; *exoX*; *recA*; *dprA*; *recFOR*; *recBCD*), individually and in concert, that suppress SPDIR mutations in wildtype *A. baylyi* by orders of magnitude. We only took a glimpse at horizontal gene transfer (*comA*), and further cellular pathways may potentially modulate SPDIR. Reports on SPDIR in other organisms are currently scarce (Harms et al., 2016) but imply that SPDIR mutations probably occur in all domains of life; although the cellular control for SPDIR mutations may vary heavily.

## Experimental Procedures

### Bacterial strains

The strains used in this study were derived from the *A. baylyi* ADP1 strain JV28 (*trpE27 rpoB1 alkM*::*nptII’*::tg4; (Kickstein et al., 2007) and are listed in Table 4. The construction of the *hisC*::’ND5i’ allele has been described elsewhere (Overballe-Petersen et al., 2013). The knockout mutations were crossed into the strains through natural transformation as described previously (Harms et al., 2007). For the Δ*recFOR* strains, the Δ*recF*, Δ*recO* and Δ*recR* deletions were introduced sequentially in that order. All constructs were verified phenotypically when applicable, and by PCR (references given in Table 4).

**Table 4:**
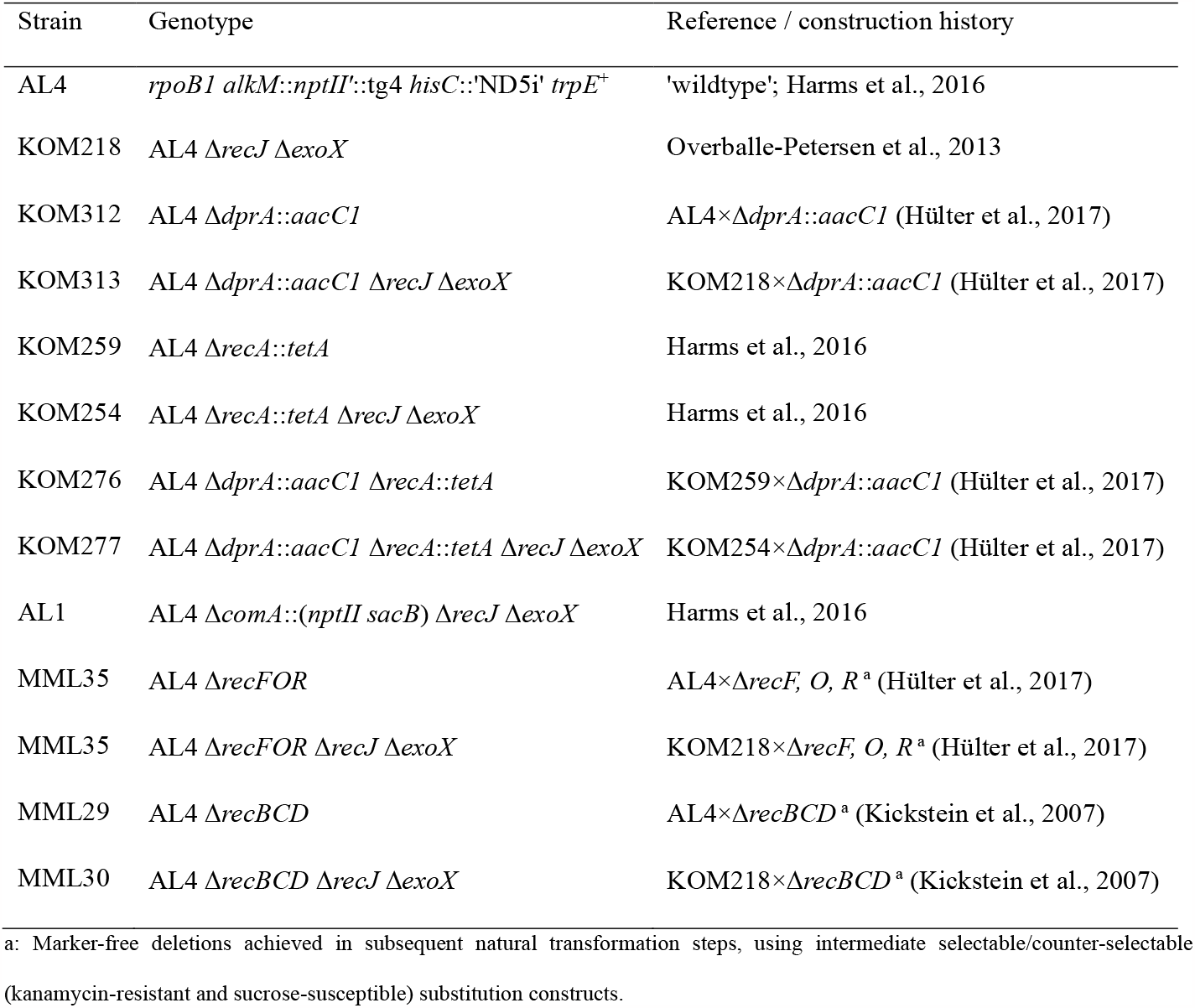
List of strains.

### Mutation experiments

The fluctuation assays were conducted as previously published (Harms et al., 2016) but with modifications (Figure 2A). Briefly, overnight cultures (ONC) were grown in LB (pH adjusted to 7.4) at 30°C with aeration/shaking. Each ONC was used to inoculate a single 20 mL LB assay 1:100. These assays were aerated for 15 hours at 30°C. The cultures were then chilled on ice, washed twice with PBS, and resuspended in 2 ml PBS. Appropriate cell dilutions were plated on LB (for CFU) and on M9 minimal medium supplied with 10 mM succinate (M9S; ten plates, 200 µl suspension per plate). The plates were incubated at 30°C for 48 hours (LB), 72 hours (M9S) or 84 hours (M9S for strains MML29 and MML30). Colonies were counted, His^+^ and CFU titers were determined, and the His^+^ frequencies were calculated as His^+^ revertant per CFU. For each group of mutation experiments, we determined the median His^+^ frequency.

### SPDIR frequency determination

*A. baylyi* His^+^ colonies were re-streaked on M9S and grown for 48 to 72 hours at 30°C, until colonies became visible. We amplified the recombinant region of the *hisC*::’ND5i’ allele from each isolate by PCR, using the DreamTaq (NEB) protocol and the primers hisC-ins-f (GACAAGCCATTTTTATTACACC) and hisC-ins-r (CAATTACGACTACACGATCG). The PCR products were processed according to the Exo-SAP protocol (NEB) and Sanger-sequenced by Azenta Life Services GeneWiz with hisC-ins-f as sequencing primer. SPDIR mutations were determined using BLAST (Figure 2C) and distinguished from other mutations (usually small deletions from three to 195 bp in size, see supplemental file S2 for full range). Identical mutations in individual fluctuation assays were considered siblings in the 15-hour-cultures and were treated as a single mutation event. The SPDIR frequency for each group of experiments was calculated as number of SPDIR mutations divided by number of all His^+^ mutations, multiplied by the median His^+^ frequency.

## Acknowledgements

Authors M.M.L. and K.H. were funded by a grant by the Norwegian Research Council (grant number 275672).

All figures used in the paper were produced using Biorender.com and are published with permission.

