## Supplemental File S1 for "The recombination initiation functions DprA and RecFOR suppress microindel mutations in *Acinetobacter baylyi* ADP1"

**Supplemental File 1: SPDIR mutations recovered in this study**

This file lists the DNA sequences of all SPDIR mutations found experimentally in this study with the *hisC*::'ND5i' detection allele (Overballe-Petersen et al., 2013) in *Acinetobacter baylyi* ADP1.

The mutations were assigned labels (‘MK38’ etc.) upon their first discovery, and those labels were kept when identical mutations were discovered in subsequent independent experiments. The labeling was continued from Harms et al. (2016).

Abbreviations:

**A** - ancestral His$^-$ DNA sequence (with two consecutive stop codons in **red**).

**R** - recombinant/mutant His$^+$ sequence.

**P** - sequence of ectopic DNA patch used for the double illegitimate recombination event (usually ADP1 chromosome); when applicable, **ND** (chromosomally inserted ‘ND5i’ DNA) or **BS** (*Bacillus subtilis* 168, donor DNA for natural transformation) are used instead.

Vertical bars indicate identical nucleotides. Extended microhomologies are indicated in **blue and bold** typeface. The illegitimate crossover joints are highlighted in yellow. When notable, stop and start codons are underlined.

**Pos** - position of the extended microhomology in the respective source DNA (*A. baylyi*: NC_005966; ND: EU153450; BS: NC_000964)

**R translated** or **R tl** - deduced amino acid sequence of the recombinant joint (**lowercase bold and in blue**: changed codons).

**ΔG$^0_{min}$** - minimal Free Energy of Hybridization, calculated for each extended microhomology (Wetmur, 2006).

**MK38.**

```
                                                                                    |-----| no net gain/loss
A   ACTACTAGGTCTTCTCTTAGCCTCAGCAGGAAAATCAGCCCAATTTGGACTTCATCCGTGACTTCCATCAGCTAGTGAAGGCCCTACCCCAGTATCAGCACTCCTTCATTCCAGTACAATAGTTATAGCTGGAGTATTTACCCTCATCCGCTTTTATCCATTAATAGAAAACAACCTCAC
    |||||||||||||||||||||||||||||||||||||||||||||||||||||||||||||||||||||||| ||||  |||||||||||||||||||||||||||||||||||||||||||||||||||||||||||||||||||||||||||||||||||||||||||||||||||||||
R   ACTACTAGGTCTTCTCTTAGCCTCAGCAGGAAAATCAGCCCAATTTGGACTTCATCCGTGACTTCCATCAGCAAGTGGTGGCCCTACCCCAGTATCAGCACTCCTTCATTCCAGTACAATAGTTATAGCTGGAGTATTTACCCTCATCCGCTTTTATCCATTAATAGAAAACAACCTCAC
    |    | | |              |    |      |      |   | |              |    |||||||||||||||            | | | |        |      |          |           | |  |    |||         | |  ||            | ||
P   AGATTTTGCTTAAAAAAGCAGAAAATAGCGGTGTGGATTTACCTCGTAATTGTGCATTGTTTATTGCCCAGCAAGTGGTGGCCAATGTACGTGAACTGGAAGGTGCACTCAATAAAGTCGTAGCAATTGCACGTTTTAAAGGTTCACAGATCGATTTGGATGTGGTGCGTGAATCCCTTA
Pos                                                                    1173<------------->1187
R translated                                                          ...I  S  k  w  w  P...
ΔG0min = -13.31 kcal/mol
```

**MK9.**

```
                                   |-- net 39 bp loss ---------------------------------------------|
A   ACTACTAGGTCTTCTCTTAGCCTCAGCAGGAAAATCAGCCCAATTTGGACTTCATCCGTGACTTCCATCAGCTAGTGAAGGCCCTACCCCAGTATCAGCACTCCTTCATTCCAGTACAATAGTTATAGCTGGAGTATTTACCCTCATCCGCTTTTATCCATTAATAGAAAACAACCTCAC
    |||||||||||||||||||||||||||||||| ||||||                                      | |||  | || ||||||| |||||||||||||||||||||||||||||||||||||||||||||||||||||||||||||||||||||||||||||||||||||||
R   ACTACTAGGTCTTCTCTTAGCCTCAGCAGGAATATCAGC---------------------------------------ACGGCGAT-CCTCAGTATCCGCACTCCTTCATTCCAGTACAATAGTTATAGCTGGAGTATTTACCCTCATCCGCTTTTATCCATTAATAGAAAACAACCTCAC
       |    |           | | | |||||||||||||                                      |||||||| |||||||||||||
P   TTCATCGTGCTGGTCAGATCCATGAACAGGAATATCAGC---------------------------------------ACGGCGAT-CCTCAGTATCCGCTAAAAGTGGTCGGTGAGACATCTACCACGGGTACTACTGTACGTTTCTGGCCAAGTGGGGATACTTTTAGCCAGAC
Pos                                 <-4595---------------------------------------------------------4628->
R translated              ...S  R  n  I  S                                     t  a  i   l  S  I  r  T...
ΔG0min = -13.74 kcal/mol (left) / -13.38 kcal/mol (right)
```

**MK27.**

```
                                                              |--------------------| no net gain/loss
A   ACTACTAGGTCTTCTCTTAGCCTCAGCAGGAAAATCAGCCCAATTTGGACTTCATCCGTGACTTCCATCAGCTAGTGAAGGCCCTACCCCAGTATCAGCACTCCTTCATTCCAGTACAATAGTTATAGCTGGAGTATTTACCCTCATCCGCTTTTATCCATTAATAGAAAACAACCTCAC
    ||||||||||||||||||||||||||||||||||||||||||||||||||||| |||    |   ||| | || || |||||||||||||||||||||||||||||||||||||||||||||||||||||||||||||||||||||||||||||||||||||||||||||||||||||||||||
R   ACTACTAGGTCTTCTCTTAGCCTCAGCAGGAAAATCAGCCCAATTTGGACTTCAACCGCATCCGCCAACCGCCAGAGAAGGCCCTACCCCAGTATCAGCACTCCTTCATTCCAGTACAATAGTTATAGCTGGAGTATTTACCCTCATCCGCTTTTATCCATTAATAGAAAACAACCTCAC
           |        |    || ||||    |    |              ||||||||||||||||||||||||||||||     |      | |      | |       |  ||   |        |    |       | |    ||||  |    |       |   || |   ||
P   GTTCGTCGAGTAAGAGTAAATCTGAGCGACTCATCAGGGTACGCGCCAGATTCAACCGCATCCGCCAACCGCCAGAGAAGCTGGCAACATCATATTGCGATTGATATTCAAAAAAACCTAAACCTGCCATAAGCTGGGATGCTTTAGCCGCTGCTGCATAACCGTTAATTTCATCAAAAC
Pos                                                 <-8798------------------8827->
R translated                                  ...T  S  t  a  s  a  n  r  q  r  R
ΔG0min = -17.29 kcal/mol
```

**MK144.**

```
                                                                                  |---| no net gain/loss
A   CTACTAGGTCTTCTCTTAGCCTCAGCAGGAAAATCAGCCCAATTTGGACTTCATCCGTGACTTCCAT-CAGCTAGTGAAGGCCCTACCCCAGTATCAGCACTCCTTCATTCCAGTACAATAGTTATAGCTGGAGTATTTACCCTCATCCGCTTTTATCCATTAATAGAAAACAACCTCAC
    |||||||||||||||||||||||||||||||||||||||||||||||||||||||||||||||||| |||| |   ||||||||||||||||||||||||||||||||||||||||||||||||||||||||||||||||||||||||||||||||||||||||||||||||||||||||||
R   CTACTAGGTCTTCTCTTAGCCTCAGCAGGAAAATCAGCCCAATTTGGACTTCATCCGTGACTTCCAT-CAGCGAACAAAGGCCCTACCCCAGTATCAGCACTCCTTCATTCCAGTACAATAGTTATAGCTGGAGTATTTACCCTCATCCGCTTTTATCCATTAATAGAAAACAACCTCAC
     |     |  |     | |  ||| | |  ||            ||          |      ||| ||||||||||||||   | || |  | || |         || |  |   |   |     |                     ||        | |              |      | |
P   ATTGCTCGGTTTGCTCTGGAATCAACGGCCAAGCTTCTTGTGCAGTGAAGCTCAAGATAGGGCTCATCCAGCGAACAAAGGCTTGAACCAAATGATACAACGCAGTTTGTGCTGAATGGCGAGCTGCTGAATCTGCCTTGGTGGTGTACTGACGATCTTTAATAATATCGAGATAGAAAT
Pos                                                                27277<---------------->27294
R translated                                                      ...S  I   S  e  q  R  P...
ΔG0min = -10.86 kcal/mol
```

**MK67.**

```
                                                                          |----| no net gain/loss
A  ACTACTAGGTCTTCTCTTAGCCTCAGCAGGAAAATCAGCCCAATTTGGACTTCATCCGTGACTTCCATCAGCTAGTGAAGGCCCTACCCCAGTATCAGCACTCCTTCATTCCAGTACAATAGTTATAGCTGGAGTATTTACCCTCATCCGCTTTTATCCATTAATAGAAAACAACCTCAC
R  ACTACTAGGTCTTCTCTTAGCCTCAGCAGGAAAATCAGCCCAATTTGGACTTCATCCGTGACTTCCATCAGGTACATAAGGCCCTACCCCAGTATCAGCACTCCTTCATTCCAGTACAATAGTTATAGCTGGAGTATTTACCCTCATCCGCTTTTATCCATTAATAGAAAACAACCTCAC
P  TTTTAGAATAATTCATGCAGTCTCCTTCGGATTTTGATCTTTGGAGAGTTTCTTTTTCTTATCTTCATCAGGTACATAGTTGTTGGCAATAATCTCACCGAACGGCCCAGTCAGATTTGGTTGTGCCAGTCTTTCACGCGTTTTAAGCGGAAGCAGTGGAAATACCAGTTCAGCAAAACG
Pos                                                            40845<-----------> 40859
R translated                                                   ...S  I  r  y  I  R...
```

ΔG$^0_{min}$ = -3.23 kcal/mol

**MK32.**

```
                                           |-- net 45 bp loss ------------------------------------------------|
A  ACTACTAGGTCTTCTCTTAGCCTCAGCAGGAAAATCAGCCCAATTTGGACTTCATCCGTGACTTCCATCAGCTAGTGAAGGCCCTACCCCAGTATCAGCACTCCTTCATTCCAGTACAATAGTTATAGCTGGAGTATTTACCCTCATCCGCTTTTATCCATTAATAGAAAACAACCTCAC
R  ACTACTAGGTCTTCTCTTAGCCTCAGCAGGAAAATCAGCCAAAATCCCGC---------------------------------------------CACCACAACCGACATCCAGTACAATAGTTATAGCTGGAGTATTTACCCTCATCCGCTTTTATCCATTAATAGAAAACAACCTCAC
P  TCCGAGTACATCTGCACCGCGACGTGCCATACTTTCAGCCAAAATCCCGC---------------------------------------------CACCACAACCGACATCCAGTACTTTCTTGCCCGCAAGGCCACCTGCATGTTCATCAATCCAGTTTAAACGTAAAGGATTAATTTG
Pos                                  <-49057------------------------------------------------------------------49094->
R translated                     ...I  S  q  n  p  a                                                 t  t  t  d  i  Q  Y...
```

ΔG$^0_{min}$ = -9.26 kcal/mol (left) / -9.28 kcal/mol (right)

**MK151.**

```
                                                                            |-| no net gain/loss
A  ACTACTAGGTCTTCTCTTAGCCTCAGCAGGAAAATCAGCCCAATTTGGACTTCATCCGTGACTTCCATCAGCTAGTGAA-GGCCCTACCCCAGTATCAGCACTCCTTCATTCCAGTACAATAGTTATAGCTGGAGTATTTACCCTCATCCGCTTTTATCCATTAATAGAAAACAACCTCA
A  ACTACTAGGTCTTCTCTTAGCCTCAGCAGGAAAATCAGCCCAATTTGGACTTCATCCGTGACTTCCATCAGCTTGGGAA-GGCCCTACCCCAGTATCAGCACTCCTTCATTCCAGTACAATAGTTATAGCTGGAGTATTTACCCTCATCCGCTTTTATCCATTAATAGAAAACAACCTCA
P  CAGGGCTTAATTGAGCAACGTCACTTAAAACAAGGTGATCGCTTGCCTGCCGAGCGTCAACTTGCCACCAGCTTGGGAATTTCCCGGCCCTCTCTGCGTGAAGCAATTCAGCAACTCAATAGTCTGGGGATTCTGGCAAGCCGTCGTGGAGACGGAACCTACATTCAGCAGCTCCCTGTG
Pos                                                          107537<------------------------>107562
R translated                                                   ...S  I  S  l  g  R   P  Y  P...
```

ΔG$^0_{min}$ = -18.37 kcal/mol

**R1520.**

```
                                                             |-- net 3 bp inserted ----------|
A  ACTACTAGGTCTTCTCTTAGCCTCAGCAGGAAAATCAGCCCAATTTGGACTTCATCCGTGACT-----TCCATCAGCTAGTGAAGGCCCTACCCCAGTATCAGCACTCCTTCATTCCAGTACAATAGTTATAGCTGGAGTATTTACCCTCATCCGCTTTTATCCATTAATAGAAAACAA
R  ACTACTAGGTCTTCTCTTAGCCTCAGCAGGAAAATCAGCCCAATTTGGACTTCATCCGAGCCAGTAAGTCCAGCC-CAAAT-AATACCCTGCCCCAGTATCAGCACTCCTTCATTCCAGTACAATAGTTATAGCTGGAGTATTTACCCTCATCCGCTTTTATCCATTAATAGAAAACAA
P  AAAGTTCAGATAGTTAATAATAGCGGCCAGATCTGCAATCCCGACCAATACCCATCCGAGCCAGTAAGTCCAGCC-CAAAT-AATACCCTGCCCCAGGCCCAATCAGATCATGTGCCATATCCACAAAAGACTTATATTGCAGATTGGCAAGCAGAAGCTCACCCATGGCGCGCATTAA
Pos                                                    <-118950-----------------------------118908->
R translated                                       ...S  S  e  p  v  s  p  a   q  i   i  P  c  P  S...
```

ΔG$^0_{min}$ = -7.08 kcal/mol (left) / -14.65 kcal/mol (right)

**K90.**

```
                                          |-- net 42 bp loss -------------------------------------|
A  ACTACTAGGTCTTCTCTTAGCCTCAGCAGGAAAATCAGCCCAATTTGGACTTCATCCGTGACTTCCATCAGCTAGTGAAGGCCCTACCCCAGTATCAGCACTCCTTCATTCCAGTACAATAGTTATAGCTGGAGTATTTACCCTCATCCGCTTTTATCCATTAATAGAAAACAACCTCAC
R  ACTACTAGGTCTTCTCTTAGCCTCAGCAGGAAAATCAGCTTA--------TTCA-------------------------------ACCCCA--A-CAGCACTCCTTCATTCCAGTACAATAGTTATAGCTGGAGTATTTACCCTCATCCGCTTTTATCCATTAATAGAAAACAACCTCAC
P  TGCGTCTGGTCAATGATTATGGGGCTTATGTAAATCAGCTTA--------TTCA-------------------------------ACCCCA--A-CAGCACTCCAATTAAGGTAGGCTGATGATGGTCAAGCATAGCCGCCCTTTGTGGGCACATTCATTTTTATCTGTGGTGATTCACT
Pos                               <-121895-------------------------------------------------------121925->
R translated                  ...K  I  S  l          f  n                                P  n     S  T  P...
```

ΔG$^0_{min}$ = -8.78 kcal/mol (left) / -12.65 kcal/mol (right)

**B11.**

```
                                                                            |--------------------| net 9 bp loss
A  ACTACTAGGTCTTCTCTTAGCCTCAGCAGGAAAATCAGCCCAATTTGGACTTCATCCGTGACTTCCATCAGCTAGTGAAGGCCCTACCCCAGTATCAGCACTCCTTCATTCCAGTACAATAGTTATAGCTGGAGTATTTACCCTCATCCGCTTTTATCCATTAATAGAAAACAACCTCAC
R  ACTACTAGGTCTTCTCTTAGCCTCAGCAGGAAAATCAGCCCAATTTGGACTTCATCCGTGACTTCCATCAGCT--T-ATT-CA--ACCCCA--A-CAGCACTCCTTCATTCCAGTACAAAAGGTATAGCTGGAGTATTTACCCTCATCCGCTTTTATCCATTAATAGAAAACAACCTCAC
P  CACTGGTTCAAGAGGTAGGTACGGAAGTGACCTTGCGTCTGGTCAATGATTATGGGGCTTATGTAAATCAGCT--T-ATT-CA--ACCCCA--A-CAGCACTCCAATTAAGGTAGGCTGATGATGGTCAAGCATAGCCGCCCTTTGTGGGCACATTCATTTTTATCTGTGGTGATTCACT
Pos                                                                     <-121897---------------------121925->
R translated                                                      ...I  S  l     f   n    P  n     S  T  P...
```

ΔG$^0_{min}$ = -7.55 kcal/mol (left) / -12.65 kcal/mol (right)

**R7.**

```
                                                                                 |---------------------| net 3 bp loss
A  ACTACTAGGTCTTCTCTTAGCCTCAGCAGGAAAATCAGCCCAATTTGGACTTCATCCGTGACTTCCATCAGCTAGTGAAGGCCCTACCCCAGTATCAGCACTCCTTCATTCCAGTACAATAGTTATAGCTGGAGTATTTACCCTCATCCGCTTTTATCCATTAATAGAAAACAACCTCAC
R  ACTACTAGGTCTTCTCTTAGCCTCAGCAGGAAAATCAGCCCAATTTGGACTTCATCCGTGACTTCCATCACCCACTTCT---CATGTTGCAGCATCAGCACTCCTTCATTCCAGTACAATAGTTATAGCTGGAGTATTTACCCTCATCCGCTTTTATCCATTAATAGAAAACAACCTCAC
P  CATTACTTAATCATCAATTATCTGGATTTGGTGCAGACGTTTTGGATCAGGAACCACCAGCAAAAGATCACCCACTTCT---CATGTTGCAGCATCAGCACCCAAATGTTCTGATTACTGGGCATATTGCTTGGGGAACAGAAGAGGCTCAACAGCGTTTATTTGATATTTTGCAAGATA
Pos                                                                   <-137928-------------------137897->
R translated                                                       ...I  t  h  f  s     c  c  S  I  S  T...
```

$\Delta G^0_{min}$ = -2.89 kcal/mol (left) / -13.67 kcal/mol (right)

**MK146.**

```
                                                                                    |-| no net gain/loss
A  ACTACTAGGTCTTCTCTTAGCCTCAGCAGGAAAATCAGCCCAATTTGGACTTCATCCGTGACTTCCATCAGCTAGTGAAGGCCCTACCCCAGTATCAGCACTCCTTCATTCCAGTACAATAGTTATAGCTGGAGTATTTACCCTCATCCGCTTTTATCCATTAATAGAAAACAACCTCAC
R  ACTACTAGGTCTTCTCTTAGCCTCAGCAGGAAAATCAGCCCAATTTGGACTTCATCCGTGACTTCCATCAGCTGGGGAAGGCCCTACCCCAGTATCAGCACTCCTTCATTCCAGTACAATAGTTATAGCTGGAGTATTTACCCTCATCCGCTTTTATCCATTAATAGAAAACAACCTCAC
P  AGAAAAAAATGATTATAATAAATGGGTGAAGAAAATAAAAAATAATTGTGAAGGAAAAATAAGTCATATAGCTGGGGAAGGTGCAGGTCTAGTTAGAAATCAATGTTATAGAGATGAGTATTTATCTAGATTATCATTTTTAACAACTGGAAAAACTTCGAAAGAAGAAAATATTCATGG
Pos                                                                172627<---------->172644
R translated                                                           ...S  w  g  R...
```

$\Delta G^0_{min}$ = -11.45 kcal/mol

**MK22.**

```
                                                                                 |----| no net gain/loss
A  ACTACTAGGTCTTCTCTTAGCCTCAGCAGGAAAATCAGCCCAATTTGGACTTCATCCGTGACTTCCATCAGCTAGTGAAGGCCCTACCCCAGTATCAGCACTCCTTCATTCCAGTACAATAGTTATAGCTGGAGTATTTACCCTCATCCGCTTTTATCCATTAATAGAAAACAACCTCAC
R  ACTACTAGGTCTTCTCTTAGCCTCAGCAGGAAAATCAGCCCAATTTGGACTTCATCCGTGACTTCCATCAACCTTCGAAGGCCCTACCCCAGTATCAGCACTCCTTCATTCCAGTACAATAGTTATAGCTGGAGTATTTACCCTCATCCGCTTTTATCCATTAATAGAAAACAACCTCAC
P  AGGTATTGAGAGCTTCAAAGCGACTCAAACTTACTAAGTTTATCTTTAGTTAAGTTTGGTTCACGGATCAACCTTCGAAGGCTCGAAAGAGCTTTCGACTACTAGGTTAAGCGTATGGCAAATTTAAAAGAAATTCGCGCCAAAGTAGCCAGTATCAAGAGCACGCAAAAGATTACTCGC
Pos                                                                   <-182927--182944->
R translated                                                       ...I  n  l  r  R  P...
```

$\Delta G^0_{min}$ = -8.43 kcal/mol

**MK4.**

```
                              |-- net 123 bp loss ----------------------------------------------------------------------------------|
A  AAAATCAGCCCAATTTGGACTTCATCCGTGACTTCCATCAGCTAGTGAAGGCCCTACCCCAGTATCAGCACTCCTTCATTCCAGTACAATAGTTATAGCTGGAGTATTTACCCTCATCCGCTTTTATCCATTAATAGAAAACAACCTCACTATTCAAACTTCAACACTTCTGGTTCTGGC
R  AAAATCAGCCCAAATTGGACCCAACCGAACCAGTATTTTTATGAA------------------------------------------------------------------------------------TCTGGTTCTGGC
P  TGCAGTCCAGCAAATTGGACCCAACCGAACCAGTATTTTTATGAA------------------------------------------------------------------------------------TCTGGTTCCCGT
Pos        <-192781------------------------------------------------------------------------------------------------------------------192823->
R tl.    ...P  I  W  T  q  p  n  q  y  f  y  e                                                                                              S  G  S...
```

$\Delta G^0_{min}$ = -8.08 kcal/mol (left) / -9.63 kcal/mol (right)

**MK26.**

```
                                                |-- net 21 bp loss ---------------------------------|
A  ACTACTAGGTCTTCTCTTAGCCTCAGCAGGAAAATCAGCCCA-ATTTGGACTTCATCCGTGACTTCCATCAGCTAGTGAAGGCCCTACCCCAGTATCAGCACTCCTTCATTCCAGTACAATAGTTATAGCTGGAGTATTTACCCTCATCCGCTTTTATCCATTAATAGAAAACAACCTCAC
R  ACTACTAGGTCTTCTCTTAGCCTCAGCAGGAAAATCAGCCCA-ATTTG---TACA--CG-G-C--CC-T--G-T--T-CA-G---AACGCCAG-ATCACCACTCCTTCATTCCAGTACAATAGTTATAGCTGGAGTATTTACCCTCATCCGCTTTTATCCATTAATAGAAAACAACCTCAC
P  CAGACTCAAACGAACACTACGATGCGGACAGCGATTTTCCCAGATTTG---TACA--CG-G-C--CC-T--G-T--T-CA-G---AACGCCAG-ATCACCACTTCGATATCCTGCACGAAGGCCTGAATCACCTCTTTTGGTTCAAGCAGAACCGAAGGTAAAATGGGAATCCACTGCTGT
Pos                                      <-194725---------------------------------------------------194680->
R translated                         ...S  P  I  c    t    r    p    c     s    e    r  s  S  t  T  P...
```

$\Delta G^0_{min}$ = -6.65 kcal/mol (left) / -17.27 kcal/mol (right)

**MK39.**

```
                                          |-- net 69 bp loss ------------------------------------------------------------------------|
A  ACTACTAGGTCTTCTCTTAGCCTCAGCAGGAAAATCAGCCCAATTTGGACTTCATCCGTGACTTCCATCAGCTAGTGAAGGCCCTACCCCAGTATCAGCACTCCTTCATTCCAGTACAATAGTTATAGCTGGAGTATTTACCCTCATCCGCTTTTATCCATTAATAGAAAACAACCTCAC
R  ACTACTAGGTCTTCTCTTAGCCTCAGCAGGAATCGCGCCC----------------------------------------------------------------------------CACAGTGCACCGCCTATAGCTGGAGTATTTACCCTCATCCGCTTTTATCCATTAATAGAAAACAACCTCAC
P  TGCATTAAAATGGGTTTTTAATAATGCAGGAATCGCGCCC----------------------------------------------------------------------------CACAGTGCACCGCCTATAGCACCTGCAATGAGCATGGAAGGCAAAATCCAAACCGATGAGGTTCCTTGAAA
Pos                         <-202033-----------------------------------------------------------------------------------------202067->
R translated             ...S  R  n  r  a  p                                                                       Q  c  t  a  Y  S...
```

$\Delta G^0_{min}$ = -8.67 kcal/mol (left) / -10.35 kcal/mol (right)

**MK94.**

```
                                                                   |-- net 9 bp loss --|
A   ACTACTAGGTCTTCTCTTAGCCTCAGCAGGAAAATCAGCCCAATTTGGACTTCATCCGTGACTTCCATCAGCTAGTGAAGGCCCTACCCCAGTATCAGCACTCCTTCATTCCAGTACAATAGTTATAGCTGGAGTATTTACCCTCATCCGCTTTTATCCATTAATAGAAAACAACCTCAC
    ||||||||||||||||||||||||||||||||||||||||||||||||||||||||||||||||    | || | |||  |  ||| ||||||||||||||||||||||||||||||||||||||||||||||||||||||||||||||||||||||||||||||||||||||||||||||||||
R   ACTACTAGGTCTTCTCTTAGCCTCAGCAGGAAAATCAGCCCAATTTGGACTTCATCCGTGACTT--A-CATC-AGT--A--CCC-ACCCCAGTATCAGCACTCCTTCATTCCAGTACAATAGTTATAGCTGGAGTATTTACCCTCATCCGCTTTTATCCATTAATAGAAAACAACCTCAC
         || |             ||||   ||     |          |||    | |     ||||  | |||| |||  |  ||| |||||||      ||    ||      | |||  |    |    |   ||       ||     | |     ||                ||
P   CAACGTAAATACAGCACCTAACTCATATGGTTTTTGTAATGCTCCTGGCACACCTGAAGTACTT--A-CATC-AGT--A--CCC-ACCCCAGCTAATTCAGCAGTTACACCTAATACTGTGTATTGCACACCAAATTTATAGTCGATATAACTGTCATCCTTTGCTGGGCCGCCTGCAAC
Pos                                                             <-242058----------------242036->
R translated                                                 ...T  y     i   s    t    h   P  S...
```

$\Delta G^0_{min}$ = -9.53 kcal/mol

**A78.**

```
                                                                          |-----------------------| net 12 bp loss
A   ACTACTAGGTCTTCTCTTAGCCTCAGCAGGAAAATCAGCCCAATTTGGACTTCATCCGTGACTTCCATCAGCTAGTGAAGGCCCTACCCCAGTATCAGCACTCCTTCATTCCAGTACAATAGTTATAGCTGGAGTATTTACCCTCATCCGCTTTTATCCATTAATAGAAAACAACCTCAC
    |||||||||||||||||||||||||||||||||||||||||||||||||||||||||||||||||||||||| |||             |||||||| |||||||||||||||||||||||||||||||||||||||||||||||||||||||||||||||||||||||||||||||||
R   ACTACTAGGTCTTCTCTTAGCCTCAGCAGGAAAATCAGCCCAATTTGGACTTCATCCGTGACTTCCATCAGCCAGTA------------CAGTATCAACACTCCTTCATTCCAGTACAATAGTTATAGCTGGAGTATTTACCCTCATCCGCTTTTATCCATTAATAGAAAACAACCTCAC
      |     ||    ||| |  |   | ||  || ||| | | ||     |      |    ||     ||||||||||||            ||||||||||||    |            ||  |  |            |  ||    | ||        | ||  | ||    ||       |
P   CTTCGGTTGTACAATCTCACTCAAATCATCAACATCTGGCAAAGCCCGTACATTTAACTGGCAATCATCAGCCAGTA------------CAGTATCAACACCTGTAATGATTGCGCCAGAACGTGCTCGCCAGTATTGCACATCGAGCCTTGCATCTACACCAGTAATCCACTTGGATTC
Pos                                                                   <-247271--------------------247248->
R translated                                                       ...I  S  q  y            S  I  n  T...
```

$\Delta G^0_{min}$ = -11.59 kcal/mol (left) / -12.31 kcal/mol (right)

**MK5.**

```
                                                                           |-----| no net gain/loss
A   ACTACTAGGTCTTCTCTTAGCCTCAGCAGGAAAATCAGCCCAATTTGGACTTCATCCGTGACTTCCATCAGCTAGTGAAGGCCCTACCCCAGTATCAGCACTCCTTCATTCCAGTACAATAGTTATAGCTGGAGTATTTACCCTCATCCGCTTTTATCCATTAATAGAAAACAACCTCAC
    ||||||||||||||||||||||||||||||||||||||||||||||||||||||||||||||||||||||||     || |||||||||||||||||||||||||||||||||||||||||||||||||||||||||||||||||||||||||||||||||||||||||||||||||||
R   ACTACTAGGTCTTCTCTTAGCCTCAGCAGGAAAATCAGCCCAATTTGGACTTCATCCGTGACTTCCATCAGCTTTCCAATGCCCTACCCCAGTATCAGCACTCCTTCATTCCAGTACAATAGTTATAGCTGGAGTATTTACCCTCATCCGCTTTTATCCATTAATAGAAAACAACCTCAC
    |    ||    ||      | |    |      |    | ||       ||        |       ||||||||||||||||||       | |  |
P   AGCCTTATCGTTTGGCACGGGCAATGTCTTTCACCAATCCATGGGCAGATCCGTTAGGCACAGGTTATCAGCTTTCCAATGCCCTGATGGCCTTTGGGCGAGGTGAATGGTTTGGCACAGGTTTAGGTCACAGTGTTCAAAAACTCTCGTATTTACCCGAAGCGCATACCGACTTTATGT
Pos                                                                         <-272208---272226->
R translated                                                             ...I  S  f  p  m  P  Y...
```

$\Delta G^0_{min}$ = -13.32 kcal/mol

**K22.**

```
                                 |-- net 57 bp loss ------------------------------------------|
A   ACTACTAGGTCTTCTCTTAGCCTCAGCAGGAAAATCAGCCCAATTTGGACTTCATCCGTGACTTCCATCAGCTAGTGAAGGCCCTACCCCAGTATCAGCACTCCTTCATTCCAGTACAATAGTTATAGCTGGAGTATTTACCCTCATCCGCTTTTATCCATTAATAGAAAACAACCTCAC
    ||||||||||||||||||||||||||||  |||                                                           ||||||||||||||||||||||||||||||||||||||||||||||||||||||||||||||||||||||||||||||||||||||||
R   ACTACTAGGTCTTCTCTTAGCCTCAGCAGCCAAA-----------------------------------------------------------CTATCAGCACTCCTTCATTCCAGTACAATAGTTATAGCTGGAGTATTTACCCTCATCCGCTTTTATCCATTAATAGAAAACAACCTCAC
       |    ||      ||      |||||||||||                                                           ||||||||||||    | |  |    |  |         ||     ||  | | |           ||     |            |
P   TTCAAGCTGTGGCGAATTTTTGGCAGCAGCCAAA-----------------------------------------------------------CTATCAGCACTCACATAATAGATGGCCTTTTGGCCTTCTTTCATACGTGCTTTATAATCGGCAAATGAGGTCGAGATTTCGTCCTGATT
Pos                         <-317466--------------------------------------------------------------317444->
R translated        ...S  L  S  s  q                                                      t  I  S  T  P...
```

$\Delta G^0_{min}$ = -9.32 kcal/mol (left) / -14.19 kcal/mol (right)

**MK33.**

```
                                                                         |---| no net gain/loss
A   ACTACTAGGTCTTCTCTTAGCCTCAGCAGGAAAATCAGCCCAATTTGGACTTCATCCGTGACTTCCATCAGCTAGTGAAGGCCCTACCCCAGTATCAGCACTCCTTCATTCCAGTACAATAGTTATAGCTGGAGTATTTACCCTCATCCGCTTTTATCCATTAATAGAAAACAACCTCAC
    ||||||||||||||||||||||||||||||||||||||||||||||||||||||||||||||||||||||| | |||||||||||||||||||||||||||||||||||||||||||||||||||||||||||||||||||||||||||||||||||||||||||||||||||||||||
R   ACTACTAGGTCTTCTCTTAGCCTCAGCAGGAAAATCAGCCCAATTTGGACTTCATCCGTGACTTCCATCAGATTGGGAAGGCCCTACCCCAGTATCAGCACTCCTTCATTCCAGTACAATAGTTATAGCTGGAGTATTTACCCTCATCCGCTTTTATCCATTAATAGAAAACAACCTCAC
     |||     |  | ||         |          |           |           |      |||||||||||||||              |||||       | |    | |   | |    ||   | |     |   ||  |     |     |   | ||    |         | |
P   TCTATATTTTTGTGTCCAGTGATAAACCTTATCCACGTATTGCAAGTGGAAGAGGTAAAGGCAAGCATCAGATTGGGAAGTTTTTTGGGCCGTATCCAAGTGCATATAGTGCCCGAGATACTTTGCTGGTATTACAAAAACTATTTAATGTGCGTCAATGTGAAAACAGTTATTTTGCCC
Pos                                                          336238<------------->336252
R translated                                                   ...S  I  r  l  g  R...
```

$\Delta G^0_{min}$ = -10.54 kcal/mol

**R1523.**

```
                                         |-- net 39 bp loss ------------------------------|
A   ACTACTAGGTCTTCTCTTAGCCTCAGCAGGAAAATCAGCCCAATTTGGACTTCATCCGTGACTTCCATCAGCTAGTGAAGGCCCTACCCCAGTATCAGCACTCCTTCATTCCAGTACAATAGTTATAGCTGGAGTATTTACCCTCATCCGCTTTTATCCATTAATAGAAAACAACCTCAC
    |||||||||||||||||||||||||||||||||||||||||| ||||    | |  ||||                     |||              ||||||||||||||||||||||||||||||||||||||||||||||||||||||||||||||||||||||||||||||||||||
R   ACTACTAGGTCTTCTCTTAGCCTCAGCAGGAAAATCAGCGCAATC---AAT--ATCC---------------------AGG-------------TCAGCACTCCTTCATTCCAGTACAATAGTTATAGCTGGAGTATTTACCCTCATCCGCTTTTATCCATTAATAGAAAACAACCTCAC
     |   |||    |   |          |  ||  |||||||||||   |||  ||||                     |||              ||||||||     |            ||||           | ||      |            |      |     |   |   |   |
P   TCAGGTAGAAGCTGGGTATCAAAGTCAATAAAGGTCAGCGCAATC---AAT--ATCC---------------------AGG-------------TCAGCACTAAACCGAAAAGCATGTGTAGTGTCGCACCAAATACCACAACCACCAGCAGCGAACAAAGCATGGTCAGTAACTCAACC
Pos                                   <-352932--------------------------------------------------352960->
R translated                       ...I  S  a  I     n    i  q                  v               S  T...
```

$\Delta G^0_{min}$ = -11.27 kcal/mol (left) / -10.22 kcal/mol (right)

**R18.**

```
                                                                             |-----| no net gain/loss
A  ACTACTAGGTCTTCTCTTAGCCTCAGCAGGAAAATCAGCCCAATTTGGACTTCATCCGTGACTTCCATCAGCTAGTGAAGGCCCTACCCCAGTATCAGCACTCCTTCATTCCAGTACAATAGTTATAGCTGGAGTATTTACCCTCATCCGCTTTTATCCATTAATAGAAAACAACCTCAC
   ||||||||||||||||||||||||||||||||||||||||||||||||||||||||||||||||||||||||   ||  ||||||||||||||||||||||||||||||||||||||||||||||||||||||||||||||||||||||||||||||||||||||||||||||||||||||
R  ACTACTAGGTCTTCTCTTAGCCTCAGCAGGAAAATCAGCCCAATTTGGACTTCATCCGTGACTTCCATCATGGAGGTAAGGCCCTACCCCAGTATCAGCACTCCTTCATTCCAGTACAATAGTTATAGCTGGAGTATTTACCCTCATCCGCTTTTATCCATTAATAGAAAACAACCTCAC
      |       | |  | | |         ||  |   |  || |      |   |  |     |||||||||||||||||||| |   ||         ||| |    |   | ||   |      |    |  || ||||   |    | | |      ||        |         |
P  GGAAGATACATTGCAATAACCAGTCCACCGACTAAAATGCCCAAAATCGCCATAATCAGCGGTTCCATCATGGAGGTAAGGCCATCGACCGCATGATCGACTTCACTTTCATAATAATTTGCGACTTTATCCAGCATTTCATCGAGTGCTCCAGATTCTTCACCAATCGCTACCATTTGG
Pos                                                                <-353463-----353483->
R translated                                                   ...T  S  I  m  e  v  R  P...
```

$\Delta G^0_{min}$ = -17.64 kcal/mol

**MK19.**

```
                                                                               |-- net 12 bp loss ---|
A  ACTACTAGGTCTTCTCTTAGCCTCAGCAGGAAAATCAGCCCAATTTGGACTTCATCCGTGACTTCCATCAGCTAGTGAAGGCCCTACCCCAGTATCAGCACTCCTTCATTCCAGTACAATAGTTATAGCTGGAGTATTTACCCTCATCCGCTTTTATCCATTAATAGAAAACAACCTCAC
   ||||||||||||||||||||||||||||||||||||||||||||||||||||||||||||||||||||||||| | | ||   || || ||     ||||||||||||||||||||||||||||||||||||||||||||||||||||||||||||||||||||||||||||||||||||||
R  ACTACTAGGTCTTCTCTTAGCCTCAGCAGGAAAATCAGCCCAATTTGGACTTCATCCGTGACTTCCATCAGC-A-T-CA---CC-ACACC-----CAGCACTCCTTCATTCCAGTACAATAGTTATAGCTGGAGTATTTACCCTCATCCGCTTTTATCCATTAATAGAAAACAACCTCAC
    ||      |       |           ||         | ||  |       |    |  ||||||||| | | ||   || |||||     ||||||         ||    |          ||
P  TCTGAATCTGCGAGAAGAACGCCTGTTAAAAATGCACCAAGAGTTGTGCTGATTCCTAATAAGTCCATCAGC-A-T-CA---CC-ACACC-----CAGCACAATAAATAGCCCAACCGCAGTGATCAGTTCTGTTGCACCACTTTTGGCAACAAAACGAAAAAATGGCCGCATGATATAG
Pos                                                                 <-410224----------------------410249->
R translated                                                    ...S  I  S  i    t     t  p      S  T...
```

$\Delta G^0_{min}$ = -12.00 kcal/mol (left) / -6.72 kcal/mol (right)

**MK10.**

```
                    |-- net 99 bp loss ----------------------------------------------------------------------------------------------------|
A  CATCCGTGACTTCCATCAGCTAGTGAAGGCCCTACCCCAGTATCAGCACTCCTTCATTCCAGTACAATAGTTATAGCTGGAGTATTTACCCTCATCCGCTTTTATCCATTAATAGAAAACAACCTCACTATTCAAACTTCAACACTTCTGGTTCTGGCTCTAATGTCACTAATGAAAACAA
   |||||||||||||||||| |      |                                                                                                            ||| | ||||||||||||||||||||||||||||||||||||||||||
R  CATCCGTGACTTCCATCACCAGTGCCAT---------------------------------------------------------------------------------------------------AATCGCAATCATCAACACTTCTGGTTCTGGCTCTAATGTCACTAATGAAAACAA
   ||     |    ||||||||||||||||                                                                                                   |||||||||||||||||||     |    |  |              ||  |   |
P  CAGTAAATAAGCCCATCACCAGTGCCAT---------------------------------------------------------------------------------------------------AATCGCAATCATCAACACACTGAGAATGGCATTGGTATCGCCATATACATGAAT
Pos                <-417321--------------------------------------------------------------------------------------------------------417354->
R tl.     ...S  I  t  s  a  i                                                                                                   i  a  i  h  N  T..
```

$\Delta G^0_{min}$ = -6.68 kcal/mol (left) / -7.44 kcal/mol (right)

**MK75.**

```
                                                                                    |--| no net gain/loss
A  ACTACTAGGTCTTCTCTTAGCCTCAGCAGGAAAATCAGCCCAATTTGGACTTCATCCGTGACTTCCATCAGCTAGTGAAGGCCCTACCCCAGTATCAGCACTCCTTCATTCCAGTACAATAGTTATAGCTGGAGTATTTACCCTCATCCGCTTTTATCCATTAATAGAAAACAACCTCAC
   |||||||||||||||||||||||||||||||||||||||||||||||||||||||||||||||||||||||||| |  |||||||||||||||||||||||||||||||||||||||||||||||||||||||||||||||||||||||||||||||||||||||||||||||||||||||
R  ACTACTAGGTCTTCTCTTAGCCTCAGCAGGAAAATCAGCCCAATTTGGACTTCATCCGTGACTTCCATCAGCTACTCCAGGCCCTACCCCAGTATCAGCACTCCTTCATTCCAGTACAATAGTTATAGCTGGAGTATTTACCCTCATCCGCTTTTATCCATTAATAGAAAACAACCTCAC
      |           |     ||                  |    |     |       |    | | |||||||||||| |       | | |    |         || | | |  ||   |    | ||   ||    |    |   |       ||  ||         |    ||
P  TACAGCGTCAGGCACCAGCCGTTCCTTTCAGCGCAATTCAACAGCAAGTGGAAAAAGAGCTCGGTAAACCGCTACTCCAGGCCTTTCAGTCTTTTGATCAACAACCTTTTGCCGCAGCATCAATTGGTCAGGTACATCGGGCTGTTTTGCCAGACGGACAAGAAGTTGTGGTCAAAGTAC
Pos                                                                    424549<----------->424537
R translated                                                             ...S  y  s  R  P...
```

$\Delta G^0_{min}$ = -10.64 kcal/mol

**O131.**

```
                                                                               |---| no net gain/loss
A  ACTACTAGGTCTTCTCTTAGCCTCAGCAGGAAAATCAGCCCAATTTGGACTTCATCCGTGACTTCCATCAGCTAGTGAAGGCCCTACCCCAGTATCAGCACTCCTTCATTCCAGTACAATAGTTATAGCTGGAGTATTTACCCTCATCCGCTTTTATCCATTAATAGAAAACAACCTCAC
   |||||||||||||||||||||||||||||||||||||||||||||||||||||||||||||||||||||||| | ||||||||||||||||||||||||||||||||||||||||||||||||||||||||||||||||||||||||||||||||||||||||||||||||||||||||||||
R  ACTACTAGGTCTTCTCTTAGCCTCAGCAGGAAAATCAGCCCAATTTGGACTTCATCCGTGACTTCCATCAGCAACTTAAGGCCCTACCCCAGTATCAGCACTCCTTCATTCCAGTACAATAGTTATAGCTGGAGTATTTACCCTCATCCGCTTTTATCCATTAATAGAAAACAACCTCAC
   |    |        | |    | || ||    | | |    | |     |   |      |   |  ||||||||||||||| ||   |  |    | | | |  ||   |     | | |         | | | |              |   |   | |   || || |
P  ATGTGTTACCTAATTTTCGTGCGCATCATCCAGAACTTAGAATTGAACTCCATAGTTTAAATAACGATCAGCAACTTAAGGCGCTTAAACGTGGTGAACTCGATTTATGTTTTATCCGAGATAATATTCAGCATCGTGATGGTATCCATAGTCAATTGGTATTAAGTGAACCACTCATTT
Pos                                                                   <-444536---444518->
R translated                                                    ...S  I  S  n  l  R  P  Y...
```

$\Delta G^0_{min}$ = -16.89 kcal/mol

**MK20.**

```
                                                                       |-----------| no net gain/loss
A  ACTACTAGGTCTTCTCTTAGCCTCAGCAGGAAAATCAGCCCAATTTGGACTTCATCCGTGACTTCCATCAGCTAGTGAAGGCCCTACCCCAGTATCAGCACTCCTTCATTCCAGTACAATAGTTATAGCTGGAGTATTTACCCTCATCCGCTTTTATCCATTAATAGAAAACAACCTCAC
   ||||||||||||||||||||||||||||||||||||||||||||||||||||||||||||||||| ||| |    |  ||||||||||||||||||||||||||||||||||||||||||||||||||||||||||||||||||||||||||||||||||||||||||||||||||||||||
R  ACTACTAGGTCTTCTCTTAGCCTCAGCAGGAAAATCAGCCCAATTTGGACTTCATCCGTGACTCCCAACGCTCAACGAAGGCCCTACCCCAGTATCAGCACTCCTTCATTCCAGTACAATAGTTATAGCTGGAGTATTTACCCTCATCCGCTTTTATCCATTAATAGAAAACAACCTCAC
     |||    |           |         ||| |        |     |  |     ||||||||||||||||||||    | ||       |         |                     |  | |   | ||                                    || |
P  TATACGGATTAGCCCCCACAAATGCAGGTAGGAATGAATGATGAATATTGATAATTTTCATCTCCCAACGCTCAACGAAGGCTTCATCCAGAATTTGCATGTAGCGTGCAAGGACTAACAAGTCATTGCCCTGCATCAATTCGTCAATTTGGGCGTAAGCTTCACGTTTGTTCTCTTTAT
Pos                                                               <-497533-----497513->
R translated                                                   ...T  p  n  a  q  r  R  P...
```

$\Delta G^0_{min}$ = -8.61 kcal/mol

**K52.**

```
                                                                         |----------| no net gain/loss
A  ACTACTAGGTCTTCTCTTAGCCTCAGCAGGAAAATCAGCCCAATTTGGACTTCATCCGTGACTTCCATCAGCTAGTGAAGGCCCTACCCCAGTATCAGCACTCCTTCATTCCAGTACAATAGTTATAGCTGGAGTATTTACCCTCATCCGCTTTTATCCATTAATAGAAAACAACCTCAC
R  ACTACTAGGTCTTCTCTTAGCCTCAGCAGGAAAATCAGCCCAATTTGGACTTCATCCGTGACTTCCATCGGCAAATCAGCACCCTACCCCAGTATCAGCACTCCTTCATTCCAGTACAATAGTTATAGCTGGAGTATTTACCCTCATCCGCTTTTATCCATTAATAGAAAACAACCTCAC
P  AAGCTTTCTTAAAATCATTGCTGTTCCAAACACAAATAATTTATACGGATTTCTCATGACAACGACATCGGCAAATCAGCACCCTACCCGTCGCGATCTTCTAAAATGGTTTACTGGTATTCCTTTTTTACCACTCGGGGCAATGGCAACAGCAGCAACACTGGCAGGATGCAATGACAG
Pos                                                                     <-508476--------508453->
R translated                                                        ...S  I  g  k  s  a  P  Y
```

$\Delta G^0_{min}$ = -17.32 kcal/mol

**MK13.**

```
                                                                            |-- net 12 bp loss ----|
A  ACTACTAGGTCTTCTCTTAGCCTCAGCAGGAAAATCAGCCCAATTTGGACTTCATCCGTGACTTCCATCAGCTAGTGAAGGCCCTACCCCAGTATCAGCACTCCTTCATTCCAGTACAATAGTTATAGCTGGAGTATTTACCCTCATCCGCTTTTATCCATTAATAGAAAACAACCTCAC
R  ACTACTAGGTCTTCTCTTAGCCTCAGCAGGAAAATCAGCCCAATTTGGACTTCATCCGTGACTTCCATCAGCAAGT------CC-A-TGC--TA--AGCACTCCTTCATTCCAGTACAATAGTTATAGCTGGAGTATTTACCCTCATCCGCTTTTATCCATTAATAGAAAACAACCTCAC
P  AAATATGCGGCACGGGTTTTTGACTACGTGCAGTCACACGTTCGGTCACTTTTACTAATATCGGATTCAGCAAGT------CC-A-TGC--TA--AGCACTCCTTTTACAACTGGATAAGCTGGCAAACAAAATAGACCAAATGATTTCATGATCATTTGAAAAAAATAAATAATATTAPos
                                           <-532435-----------------------532409->
R translated                                                           ...I  S  f  s        m   l      S  T  P  S...
```

$\Delta G^0_{min}$ = -7.40 kcal/mol (left) / -15.36 kcal/mol (right)

**MK8.**

```
                                            |-- net 150 bp loss ---------------------------------------------------------------------------------------|
A  GTCTTCTCTTAGCCTCAGCAGGAAAATCAG-CCCAATTTGGACTTCATCCGTGACTTCCATCAGCTAGTGAAGGCCCTACCCCAGTATCAGCACTCCTTCA(90xN)TCTGGTTCTGGCTCTAATGTCACTAATGAAAAAATGCGTTTCTGGAGTCCAGAGGTTCGTGAGTTAGAGCCTTAT
R  GTCTTCTCTTAGCCTCAGCAGGAAAATCAG-CCCATAGCAC-------------------------------------------------------------------GCATCATCCCCACCACACGATCACCTGCAAAAATACGTTTCTGGAGTCCAGAGGTTCGTGAGTTAGAGCCTTAT
P  ATTGTGCTCAATAGCTTCATCTTCTTTCAGCCCCATAGCAC-------------------------------------------------------------------GCATCATCCCCACCACACGATCACCTGCAAAAATACGCATTAGATCATCTTCAAGTGAGAGATAAAAACGTGAA
Pos                           <-638643---------------------------------------------------------------------------------------------638592->
R translated               ...I  S  P  i  a  r                                                                                          I  r  p  t  t  r  s  p  a  K  i  R...
```

$\Delta G^0_{min}$ = -8.77 kcal/mol (left) / -9.80 kcal/mol (right)

**MK98.**

```
                                                 |-- net 6 bp loss ---------------------------|          |-- direct repeat ----------------------------------|
A  ACTACTAGGTCTTCTCTTAGCCTCAGCAGGAAAATCAGCCCAATTTGGACTTCA-TCCGTGACTTCCATCAGCTAGTGAAGGCCCTACCC-CAGTATCAGCACTCCTTCATTCCAGTACAATAGTTATAGCTGGAGTATTTACCCTCATCCGCTTTTATCCATTAATAGAAAACAACCTC
R  ACTACTAGGTCTTCTCTTAGCCTCAGCAGGAAAATCAGCACTCCTT-CATTCCAGTACAATAGTT--AT-AGCT-G-G-AGTATTTACCC-CAGTATCAGCACTCCTTCATTCCAGTACAATAGTTATAGCTGGAGTATTTACCCTCATCCGCTTTTATCCATTAATAGAAAACAACCTC
P  ACTTCCATCAGCTAGTGAAGGCCCTACCCCAGTATCAGCACTCCTT-CATTCCAGTACAATAGTT--AT-AGCT-G-G-AGTATTTACCCTCATCCGCTTTTATCCATTAATAGAAAACAACCTCACTATTCAAACTTCAACACTTCTGGTTCTGGCTCTAATGTCACTAATGAAAAAAT
Pos                                        <-109 of 'ND5i' insert ---------------------------------161-> (12467..12519 of EU153450)
R translated                          ...K  I  S  t  p  s  f  q  y  n  s  y    S  w    s  I  Y  P  S  I  S  T  P  S  F  Q  Y  N  S  Y  S  W  S  I  Y  P  H...
```

**$\Delta G^0_{min}$ = -6.84 kcal/mol + -6.04 kcal/mol = -12.88 kcal/mol / Class 2**

Note: The ectopic DNA patch originates from sequence downstream of stop codon, inside the 'ND5i' DNA segment (**purple**) and results in a duplication *in frame*, separated by two nucleotides.

**K49.**

```
                                                                           |----| no net gain/loss
A  ACTACTAGGTCTTCTCTTAGCCTCAGCAGGAAAATCAGCCCAATTTGGACTTCATCCGTGACTTCCATCAGCTAGTGAAGGCCCTACCCCAGTATCAGCACTCCTTCATTCCAGTACAATAGTTATAGCTGGAGTATTTACCCTCATCCGCTTTTATCCATTAATAGAAAACAACCTCAC
R  ACTACTAGGTCTTCTCTTAGCCTCAGCAGGAAAATCAGCCCAATTTGGACTTCATCCGTGACTTCCATCAGGTACATAAGGCCCTACCCCAGTATCAGCACTCCTTCATTCCAGTACAATAGTTATAGCTGGAGTATTTACCCTCATCCGCTTTTATCCATTAATAGAAAACAACCTCAC
P  AGGCGAAGGTGGATACGGATTTTCATTGGTATTGAGTTTAAGTAAGTTCTGGATTTTAGGTTGCTCACCAGGTACATAAGGCTCTAACTCACGAACCTCTGGACTCCAGAAACGCATTTTTTCATTAGTGACATTCATCTTTCTATTCTCAAAAAAGGGCTGAAAATCAGCCCCGTAGAG
Pos                                                                     <-657707-----657687->
R translated                                                        ...S  I  r  y  i  R  P  Y...
```

$\Delta G^0_{min}$ = -12.55 kcal/mol

**A4.**

```
                             |-- net 42 bp loss --------------------------------|
A  ACTACTAGGTCTTCTCTTAGCCTCAGCAGGAAAATCAGCCCAATTTGGACTTCATCCGTGACTTCCATCAGCTAGTGAAGGCCCTACCCCAGTATCAGCACTCCTTCATTCCAGTACAATAGTTATAGCTGGAGTATTTACCCTCATCCGCTTTTATCCATTAATAGAAAACAACCTCAC
R  ACTACTAGGTCTTCTCTTAGCCTCACCAGG------------------------------------------TACATAAGGCCCTACCCCAGTATCAGCACTCCTTCATTCCAGTACAATAGTTATAGCTGGAGTATTTACCCTCATCCGCTTTTATCCATTAATAGAAAACAACCTCAC
P  TAAGTTCTGGATTTTAGGTTGCTCACCAGG------------------------------------------TACATAAGGCTCTAACTCACGAACCTCTGGACTCCAGAAACGCATTTTTTCATTAGTGACATTCATCTTTCTATTCTCAAAAAAGGGCTGAAAATCAGCCCCGTAGAG
Pos                       <-657709------------------------------------------------657687->
R translated           ...L  t  R                                              y  i  R  P  Y...
```

$\Delta G^0_{min}$ = -9.21 kcal/mol (left) / -8.64 kcal/mol (right)

**MK42.**

```
                               |-- net 39 bp loss ---------------------------------------------------|
A  ACTACTAGGTCTTCTCTTAGCCTCAGCAGGAAAATCA--GCCCAATTTGGACTTCATCCGTGACTTCCATCAGCTAGTGAAGGCCCTACCCCAGTATCAGCACTCCTTCATTCCAGTACAATAGTTATAGCTGGAGTATTTACCCTCATCCGCTTTTATCCATTAATAGAAAACAACCTCAC
R  ACTACTAGGTCTTCTCTTAGCCTCAGCAGCATAATCATAGCCA-----------------------------------------ATGAATCCAATATAACCACTCCTTCATTCCAGTACAATAGTTATAGCTGGAGTATTTACCCTCATCCGCTTTTATCCATTAATAGAAAACAACCTCAC
P  GAAATCGGTATTTACATATTGTGATGCAGCATAATCATAGCCA-----------------------------------------ATGAATCCAATATAACCACCCTGAAATTTACTATTTATACTTTTTGTAGAATATTCAGAATTCTGAAAAATGATAAATTCAGTCAGCTCGGTATTATG
Pos                          <-659756---------------------------------------------------------------659794->
R translated             ...S  t  i  i  i  a  n                                        e  s  n  i  t  T  P...
```

$\Delta G^0_{min}$ = -10.08 kcal/mol (left) / -8.44 kcal/mol (right)

**MK2.**

```
                                                                      |-- net 6 bp loss ------------|
A  ACTACTAGGTCTTCTCTTAGCCTCAGCAGGAAAATCAGCCCAATTTGGACTTCATCCGTGACTTCCATCAGCTAGTGAAGGCCCTACCCCAGTATCAGCACTCCTTCATTCCAGTACAATAGTTATAGCTGGAGTATTTACCCTCATCCGCTTTTATCCATTAATAGAAAACAACCTCACT
R  ACTACTAGGTCTTCTCTTAGCCTCAGCAGGAAAATCAGCCCAATTTGGACTTCATCCGTGACTTCCATCAGC-ATTCTTGCCGTTAACCCC----CA-CACCACTTCATTCCAGTACAATAGTTATAGCTGGAGTATTTACCCTCATCCGCTTTTATCCATTAATAGAAAACAACCTCACT
P  AATGAAGGCGGATTCAGTAAAAATGGCCACTCTTTCTGATAACCCAATCCATAATTAAATAAAGAAATCAGC-ATTCTTGCCGTTAACCCC----CA-CACCACTTCATCTTCGACCTGTAAACTTGGGAAATAAAGCGTTTGTTTGATGTATTGTACTTTGTAAGGGGTTGGTGGCGCGT
Pos                                                                <-669516---------------------------669480->
R translated                                                    ...I  S  t  l  a  v  n  p     h  t  t  S  F...
```

$\Delta G^0_{min}$ = -5.84 kcal/mol (left) / -8.81 kcal/mol (right)

**MK34.**

```
                                                      |-- net 3 bp loss -------------|
A  ACTACTAGGTCTTCTCTTAGCCTCAGCAGGAAAATCAGCCCAATTTGGACTTCATCCGTGACTTCCATCAGCTAGTGAAGGCCCTACCCCAGTATCAGCACTCCTTCATTCCAGTACAATAGTTATAGCTGGAGTATTTACCCTCATCCGCTTTTATCCATTAATAGAAAACAACCTCAC
R  ACTACTAGGTCTTCTCTTAGCCTCAGCAGGAAAATCAGCCCAATTTGGACTCCATACC-ACCAGCAATCA-C-AATTTCTGCACTACCCCAGTATCAGCACTCCTTCATTCCAGTACAATAGTTATAGCTGGAGTATTTACCCTCATCCGCTTTTATCCATTAATAGAAAACAACCTCAC
P  GATAACCAGAACGTGCTTTAGGCAAAACATAAGGCGCGTTACTCATCGACTCCATACC-ACCAGCAATCA-C-AATTTCTGCACTACCCGCCTTAATCATATCTACAGCTTGCATCACTGCTTTCATGCCAGAACCACACAATTTATTGATCGTGACAGCACCTGTTGTATCAGGCAGTC
Pos                                                <-683085---------------------------683125->
R translated                                   ...W  T  p  y  h  q  q  s  q  f  l  h  Y  P...
```

$\Delta G^0_{min}$ = -7.07 kcal/mol (left) / -9.81 kcal/mol (right)

**MK44.**

```
                                                                |-----------| no net gain/loss
A  ACTACTAGGTCTTCTCTTAGCCTCAGCAGGAAAATCAGCCCAATTTGGACTTCATCCGTGACTTCCATCAGCTAGTGAAGGCCCTACCCCAGTATCAGCACTCCTTCATTCCAGTACAATAGTTATAGCTGGAGTATTTACCCTCATCCGCTTTTATCCATTAATAGAAAACAACCTCAC
R  ACTACTAGGTCTTCTCTTAGCCTCAGCAGGAAAATCAGCCCAATTTGGACTTCATCCGTGACTACCATTAATCACCGAAGGCCCTACCCCAGTATCAGCACTCCTTCATTCCAGTACAATAGTTATAGCTGGAGTATTTACCCTCATCCGCTTTTATCCATTAATAGAAAACAACCTCAC
P  ATCCACGTCAGCCATAACCACGTCAATGGTCGCCAAATAAATGATCAAGTCCGAAACCAGACTACCATTAATCACCGAAGGCATTTGCTGCAAGTGGGTGAAGGTTGGCGTACGAATACGTGTACGATAGCTCATGGTTGACTTGTCAGATGTGAGGTAATACGTGCTCGCACCTTTTAC
Pos                                                             <-717313-------717291->
R translated                                                ...V  T  t  I  n  h  r  R  P...
```

$\Delta G^0_{min}$ = -13.04 kcal/mol

**S75.**

```
                                     |-- net 90 bp loss ---------------------------------------------------------------------|
A  ACTACTAGGTCTTCTCTTAGCCTCAGCAGGAAAATCAGCCCAATTTGGACTTCATCCGTGACTTCCATCAGCTAGTGAAGGCCCTACCCCAGTATCAGCACTCCTTCATTCCAGTACAATAGTTATAGCTGGAGTATTTACCCTCATCCGCTTTTATCCATTAATAGAAAACAACCTCAC
R  ACTACTAGGTCTTCTCTTAGCCTCAGCAGGAAAA------------------------------------------------------------------------------------------ATAGCTGGAGTATTTACCCTCATCCGCTTTTATCCATTAATAGAAAACAACCTCAC
P  AGCATGCAAACTAAGCCATGCAGAAGCAGGAAAA------------------------------------------------------------------------------------------ATAGCCGAAGCAAACAATAGCCCCTTGGGATTAAGCAGCGTTGCACAAAACAGTTC
Pos                        <-760735-------------------------------------------------------------------------------------------760716->
R translated            ...S  R  K  n                                                                                            S  W  S...
```

$\Delta G^0_{min}$ = -13.32 kcal/mol (left) / -6.69 kcal/mol (right)

**MK49.**

```
                                                                                                                 net 1 bp inserted |---------|
A  ACTACTAGGTCTTCTCTTAGCCTCAGCAGGAAAATCAGCCCAATTTGGACTTCATCCGTGACTTCCATCAGCTAGTGAAGGCCCTACCCCAGTATCAGCACTCCTTCATTCCAGTACAATAGTTATAGCTGGAGTATTTACCCTCATCCGCTTT-TATCCATTAATAGAAAACAACCTCAC
R  ACTACTAGGTCTTCTCTTAGCCTCAGCAGGAAAATCAGCCCAATTTGGACTTCATCCGTGACTTCCATCAGCTAGTGAAGGCCCTACCCCAGTATCAGCACTCCTTCATTCCAGTACAATAGTTATAGCTGGAGTATTTACCCTCATCAGGAGAATATTCATTAATAGAAAACAACCTCAC
P  ATGGCTTTACGTCTTGCTGGTGAGTTGCTTGATGCAGCTGAAGGTAAAGGTGCAGCAATCAAAAAACGTGAAGATGTGCATCGTATGGCTGAAGCCAACAAAGCCTTCTCTCACTACCGTTTCTAAGCGGATAAAACAGTCCCTCATCAGGAGAATATTCATGCGTACAGCAGCTCGATTC
Pos                                                                                                                      <-865908------865929->
R translated                                                                                               m  v  i  a  g  v  f  t  p  i  r  r  i  f  I  N  R  K  Q  P...
```

$\Delta G^0_{min}$ = -8.18 kcal/mol

Note: Clearly a frameshift SPDIR mutant, but no new obvious start codon found (ATG, TTG, or GTG). Possibly multiple ATA codons (underlined) for rare translation initiation as demonstrated in *E. coli* [Hecht et al. (2017), Nucleic Acids Res 45:3625-26; https://doi.org/10.1093/nar/gkx070].

```
MK87.                                                                                                                   |-----| no net gain/loss
A  AGCTAGTGAAGGCCCTACCCCAGTATCAGCACTCCTTCATTCCAGTACAATAGTTATAGCTGGAGTATTTACCCTCATCCGCTTTTATCCATTAATAGAAAACAACCTCACTATTCAAACTTCAACACTTCTGGTTCTGGCTCTAATGTCACTAATGAAAAAATGCGTTTCTGGAGTCCA
   ||||||||||||||||||||||||||||||||||||||||||||||||||||||||||||||||||||||||||||||||||||||||||||||||||||||||||||||     |||  |||||||||||||||||||||||||||||||||||||||||||||||||||||||||||||||||
R  AGCTAGTGAAGGCCCTACCCCAGTATCAGCACTCCTTCATTCCAGTACAATAGTTATAGCTGGAGTATTTACCCTCATCCGCTTTTATCCATTAATAGAAAACAACCTCACATGTCACACTTCAACACTTCTGGTTCTGGCTCTAATGTCACTAATGAAAAAATGCGTTTCTGGAGTCCA
             | ||          | |     |       |      |||                  |    ||     |        |     | |   ||| ||   || |    |||||||||||||||     ||    | |      |        | | |  ||        | |     |||      |||
P  GTGCTTGTGCGACCTGTGATGGTTTTTTCTATAAAAACCAGAAAGTCATGGTTGTCGGTGGTGGTAATACTGCAGTTGAAGAAGCTTTGTATTTATCCAATATTGCCTCACATGTCACACTGGTGCATCGCCGTGATAGCTTGCGCTCTGAAAAAATCTTGCAAGATCATCTTTTTGCCA
Pos                                                                                                               873330<--------------->873315
R translated                                                                                                            m  s  h  S...
ΔG0min = -8.41 kcal/mol


B18.                                                                          |--| no net gain/loss
A  ACTACTAGGTCTTCTCTTAGCCTCAGCAGGAAAATCAGCCCAATTTGGACTTCATCCGTGACTTCCATCAGCTAGTGAAGGCCCTACCCCAGTATCAGCACTCCTTCATTCCAGTACAATAGTTATAGCTGGAGTATTTACCCTCATCCGCTTTTATCCATTAATAGAAAACAACCTCAC
   |||||||||||||||||||||||||||||||||||||||||||||||||||||||||||||||||||||||||| || |||||||||||||||||||||||||||||||||||||||||||||||||||||||||||||||||||||||||||||||||||||||||||||||||||||||||||||||
R  ACTACTAGGTCTTCTCTTAGCCTCAGCAGGAAAATCAGCCCAATTTGGACTTCATCCGTGACTTCCATCAGATATTGCAGGCCCTACCCCAGTATCAGCACTCCTTCATTCCAGTACAATAGTTATAGCTGGAGTATTTACCCTCATCCGCTTTTATCCATTAATAGAAAACAACCTCAC
            | || |  | ||  |     | | |    |          |  | |     | | ||   | ||||||||||||     |     |   |         |         ||    |    |   | || | | ||        |       |  ||                     |   |
P  TGCGGGTTCTATTTTTCTGGCACCTCTGGCAGACAAAATTGGTCGCCGTTTACTGATTTTAATTTGTTTAGCTATTGCAGGCATCAGCATGCTCGCCTGTGGTTTGGTGCATAGCCACAGTATGTTGGCAGCATTACGCTTTGTGACAGGTATTGGGGTGGGGGGTATTTTAGCCAGTAG
Pos                                                                 968658 <-----------> 968670
R translated                                                            ...S  y  c  R  P...
ΔG0min = -12.91 kcal/mol


MK150.                                                                      |--| no net gain/loss
A  ACTACTAGGTCTTCTCTTAGCCTCAGCAGGAAAATCAGCCCAATTTGGACTTCATCCGTGACTTCCATCAGCTAGTGAAGGCCCTACCCCAGTATCAGCACTCCTTCATTCCAGTACAATAGTTATAGCTGGAGTATTTACCCTCATCCGCTTTTATCCATTAATAGAAAACAACCTCAC
   |||||||||||||||||||||||||||||||||||||||||||||||||||||||||||||||||||||||| |  ||||||||||||||||||||||||||||||||||||||||||||||||||||||||||||||||||||||||||||||||||||||||||||||||||||||||||||||||
R  ACTACTAGGTCTTCTCTTAGCCTCAGCAGGAAAATCAGCCCAATTTGGACTTCATCCGTGACTTCCATCAGCGAAAGAAGGCCCTACCCCAGTATCAGCACTCCTTCATTCCAGTACAATAGTTATAGCTGGAGTATTTACCCTCATCCGCTTTTATCCATTAATAGAAAACAACCTCAC
   |  |  |     |         | |    |    |||    | |||       |       || || |||||||||||||||        |  |    |    |    | ||  |    |  ||     | |    |    |   |  ||   |   ||          |
Ab AGCAGAACAAAATGGTGCGCATTTACATCTACTTGCAGGTGCAATTGCTGGGATTGATGGAATTGCAGCAGCGAAAGAAGGCGGCTTGGAAAAAGTGACCTACAAAGGCTGCAAAAGTCAAATAGTTGGCGTGGCAGTTATGCCGAGCAGTTAATTGATTTGGATCAGGTACATACAGT
Pos                                                             982222 <------------------> 982241
R translated                                                   ...T  S  I  S  e  r  R  P...
ΔG0min = -16.15 kcal/mol


MK149.                                                                      |-- net 6 bp inserted -----|
A  ACTACTAGGTCTTCTCTTAGCCTCAGCAGGAAAATCAGCCCAATTTGGACTTCATCCGTGACTTCCATCAGC-T----AGTGAAGG--CCCTACCCCAGTATCAGCACTCCTTCATTCCAGTACAATAGTTATAGCTGGAGTATTTACCCTCATCCGCTTTTATCCATTAATAGAAAACA
   |||||||||||||||||||||||||||||||||||||||||||||||||||||||||||||||||||||||| |     ||| | ||      | |  ||| ||||||||||||||||||||||||||||||||||||||||||||||||||||||||||||||||||||||||||||||||
R  ACTACTAGGTCTTCTCTTAGCCTCAGCAGGAAAATCAGCCCAATTTGGACTTCATCCGTGACTTCCATCAGCCTGGAAAGT-ACGGATTTTTTCATCAGCATCAGCACTCCTTCATTCCAGTACAATAGTTATAGCTGGAGTATTTACCCTCATCCGCTTTTATCCATTAATAGAAAACA
   |  |  ||   |     | || |  |  ||  |           |      ||     |      |||||||||||||||| ||||||||||||||||||||||||||       |           |         |         ||  | |        |  | |  |||     |
P  AAGAGCAGCGGTGTGGTAAGTCGGAAGAGTGATCAAGTGATGAAACACACCTGCTTCACGTGCAGCATCAGCCTGGAAAGT-ACGGATTTTTTCATCAGCATCAGCAGCTAATTCAGTGTTATCGTACTCAGCACTCATCAATTTAGCGCGGTCATACGCTGAAACATCTTTACCTTCAG
Pos                                                                <-1081210-----------------------1081170->
R translated                                                     ...S  I  S  l  e  s   t  d  f  f  i  S  I  S  T...
ΔG0min = -7.78 kcal/mol (left) / -11.87 kcal/mol (right)


MK7.                                                                          |-----------| net 3 bp inserted
A  ACTACTAGGTCTTCTCTTAGCCTCAGCAGGAAAATCAGCCCAATTTGGACTTCATCCGTGACTTCCATCAGCTAGT-GAAG-GCC-CTACCCCAGTATCAGCACTCCTTCATTCCAGTACAATAGTTATAGCTGGAGTATTTACCCTCATCCGCTTTTATCCATTAATAGAAAACAACCT
   |||||||||||||||||||||||||||||||||||||||||||||||||||||||||||||||||||||||||| |  | | |||   |||||||||||||||||||||||||||||||||||||||||||||||||||||||||||||||||||||||||||||||||||||||||||||||
R  ACTACTAGGTCTTCTCTTAGCCTCAGCAGGAAAATCAGCCCAATTTGGACTTCATCCGTGACTTCCATCAGCTACTTCATGCGCCATTACCCCAGTATCAGCACTCCTTCATTCCAGTACAATAGTTATAGCTGGAGTATTTACCCTCATCCGCTTTTATCCATTAATAGAAAACAACCT
         |                                                 |  |  ||| |||||||||||||||||||||||||   |
P  GGCTAAAGCGAGTAAACCTTGTCCCTTAAAGGCTTCTTGTGAACGACTGTAATGACGTTGTGTAACATGAGCTACTTCATGCGCCATTACCCCAGCCACCTCATCAATATTTCGTGCATTTGCGATTAAACCTGTATTTAAGGCAAATAACCCACCTGGAACAGCAAATGCATTAATTTG
Pos                                                                <-1326406------------1326377->
R translated                                                     ...S  I  S  y  f  m  r  h  Y  P  S...
ΔG0min = -20.34 kcal/mol


MK16.                                                                         |-------| no net gain/loss
A  ACTACTAGGTCTTCTCTTAGCCTCAGCAGGAAAATCAGCCCAATTTGGACTTCATCCGTGACTTCCATCAGCTAGTGAAGGCCCTACCCCAGTATCAGCACTCCTTCATTCCAGTACAATAGTTATAGCTGGAGTATTTACCCTCATCCGCTTTTATCCATTAATAGAAAACAACCTCAC
   ||||||||||||||||||||||||||||||||||||||||||||||||||||||||||||||||||||||||||      |||   |||||||||||||||||||||||||||||||||||||||||||||||||||||||||||||||||||||||||||||||||||||||||||||||||||||
R  ACTACTAGGTCTTCTCTTAGCCTCAGCAGGAAAATCAGCCCAATTTGGACTTCATCCGTGACTTCCATCAGCTACATGAGGTTCTACCCCAGTATCAGCACTCCTTCATTCCAGTACAATAGTTATAGCTGGAGTATTTACCCTCATCCGCTTTTATCCATTAATAGAAAACAACCTCAC
     |       | |  ||        |         |||     |        |||     |      ||  |||||||||||||||||||||  ||  ||   |         |     |          |          |          |         ||          ||||
P  TGTGAAGACACATGGCTAGCTTAGATTTACCTGATCATTTAATCCACACCTTATCACAAGTTTTAAATCAGCTACATGAGGTTCTACCAAAGATTCGTCAAGAACCTGATTTCTCAGCACCTGCATTTAAATGGCAAGATAAACAACTTAAGCCCATTTTTGACCCCAAAAAATTTGATT
Pos                                                                  <-1331952----1331931->
R translated                                                       ...I  S  y  m  R  m  Y  P...
ΔG0min = -14.25 kcal/mol
```

**MK1.**

```
                                                |-- net 108 bp loss ----------------------------------------------------------------------------------------------------|
A   CAGGAAAATCAGCCCAATTTGGACTTCATCCGTGACTTCCATCAGCTAGTGAAGGCCCTACCCCAGTATCAGCACTCCTTCATTCCAGTACAATAGTTATAGCTGGAGTATTTACCCTCATCCGCTTTTATCCATTAATAGAAAACAACCTCACTATTCAAACTTCAACACTTCTGGTTCT
    |||||||||||||||||||||||||||||||||||| || ||| | |  |                                                                                                         ||||||||||||||||||||||
R   CAGGAAAATCAGCCCAATTTGGACTTCATCCGTGACTACCTTCATCCACAG---------------------------------------------------------------------------------------------------------AAACTTCAACACTTCTGGTTCT
                                    ||||||||||||||||||                                                                                                         |||||||||||    |    | |
P   CTCCCAAGAACTTGGTTTGACCAAATTTAAACTGACTACCTTCATCCACAG---------------------------------------------------------------------------------------------------------AAACTTCAACAAAAATATGTTT
Pos     <-1374284----------------------------------------------------------------------------------------------------------1374255->
R translated                    ...V  T  t  f  i  h  r                                                                                                      N  F  N  T...
```

$\Delta G^0_{min}$ = -9.00 kcal/mol (left) / -13.56 kcal/mol (right)

**MK128.**

```
              |-- net 153 bp loss ----------------------------------------------------------------------------------------------|
A   CACTACACTCCCACTACTAGGTCTTCTCTTAGCCTCAGCAGGAAAATCAGCCCAATTTGGACTTCATCCGTGACTTCCATCAGCTAGTGAAGGCCCTACCCCAGTATCAGCACTCCTTCA(60bp)AAACAACCTCACTATTCAAACTTCAACACTTCTGGTTCTGGCTCTAATGTCACTAA
    |||||||||||||||       | | |                                                                                              ||   |     ||     ||||||||||||||||||||||||||||||||||
R   CACTACACTCACACTCGATCCACATATTG--------------------------------------------------------------------------------------------CCAGCATTCAGTTTACGGGTTCAACACTTCTGGTTCTGGCTCTAATGTCACTAA
         ||||||||||||||||||||||||                                                                                            ||||||||||||||||||||||||||  |||  |      |     |
P   TGAAGCACTCACACTCGATCCACATATTG--------------------------------------------------------------------------------------------CCAGCATTCAGTTTACGGGTTCAACACAGGTGGGCCGTATCATTGGTGCCAATG
Pos     <-1427297--------------------------------------------------------------------------------------------------------------------------1427241->
tl ...Y  T  P  T  r  s  t  y  c                                                                                                q  h  s  v  y  g  F  N  T  S  G...
```

$\Delta G^0_{min}$ = -14.25 kcal/mol (left) / -10.14 kcal/mol (right)

**MK23.**

```
                                                                                |------------------| net 3 bp inserted
A   ACTACTAGGTCTTCTCTTAGCCTCAGCAGGAAAATCAGCCCAATTTGGACTTCATCCGTGACTTCCATCAGCTAGT---GAAGGCCC-TACCCCAGTATCAGCACTCCTTCATTCCAGTACAATAGTTATAGCTGGAGTATTTACCCTCATCCGCTTTTATCCATTAATAGAAAACAACCTCAC
    |||||||||||||||||||||||||||||||||||||||||||||||||||||||||||||||||||||||||| | |||    |    ||  | ||||||||||||||||||||||||||||||||||||||||||||||||||||||||||||||||||||||||||||||||||||||||||||||
R   ACTACTAGGTCTTCTCTTAGCCTCAGCAGGAAAATCAGCCCAATTTGGACTTCATCCGTGACTTCCATCATC-AGTTTTGGCAACCAGTTCCCCAGTATCAGCACTCCTTCATTCCAGTACAATAGTTATAGCTGGAGTATTTACCCTCATCCGCTTTTATCCATTAATAGAAAACAACCTCAC
    |         |     |||     |  | |||||| || |   |    ||  | ||      |||| |||||| ||||||||||||||||||||||||       ||                ||  |   |   |  |    |   |                          ||
P   ATCATTCCCCCCACGTTTAAAGAGAACAAGAAAATGAGACTCTTCGAGAGGTTATAACCTGCTTCAATCATC-AGTTTTGGCAACCAGTTCCCCAGTGCATAAACCATAATCAGTGTCATAAAAAATTGAGCACCAGAACATCATCGTACTAAACGCACGCTGTTCCTGAAAAATCATTTTGACT
Pos                                                                        <-1432395-----------------1432429->
R translated                                                           ...T  S  I  I   s  f  g  n  q  f  P  S...
```

$\Delta G^0_{min}$ = -13.31 kcal/mol left / -9.44 kcal/mol (right)

**MK130.**

```
                                                                                  |-------| no net gain/loss
A   ACTACTAGGTCTTCTCTTAGCCTCAGCAGGAAAATCAGCCCAATTTGGACTTCATCCGTGACTTCCATCAGCTAGTGAAGGCCCTACCCCAGTATCAGCACTCCTTCATTCCAGTACAATAGTTATAGCTGGAGTATTTACCCTCATCCGCTTTTATCCATTAATAGAAAACAACCTCAC
    ||||||||||||||||||||||||||||||||||||||||||||||||||||||||||||||||||||||||||||| ||| |||||||||||||||||||||||||||||||||||||||||||||||||||||||||||||||||||||||||||||||||||||||||||||||||||||
R   ACTACTAGGTCTTCTCTTAGCCTCAGCAGGAAAATCAGCCCAATTTGGACTTCATCCGTGACTTCCATCAGCCAGTCAAGACCCTACCCCAGTATCAGCACTCCTTCATTCCAGTACAATAGTTATAGCTGGAGTATTTACCCTCATCCGCTTTTATCCATTAATAGAAAACAACCTCAC
     | | |         | |    | |          |        |||||   |       |   |   |  |||||||||||||  | |            | | |  |  |  |                 |  | | ||   |||  |||       |    | |   | | | | | |||
P   CCGATTGTTCGACTTTTACTGCAAACACATCGGCACCCGATTTTTTGGCAATTCGATAGGCACTGTCTGGGCCAGTCAAGACCTCATCTGCCATAATGGCCACATCAAAACGTGCGGTCAGACGTGCTAAAGCATCTGTATTTTCAATCGCACAGGGCTGCTCAATCAGATCGATCCCAC
Pos                                                                       1445455<----------->1445467
R translated                                                                 ...S  q  s  R  P...
```

$\Delta G^0_{min}$ = -10.20 kcal/mol

**MK101.**

```
                                                                              |--------| no net gain/loss
A   ACTACTAGGTCTTCTCTTAGCCTCAGCAGGAAAATCAGCCCAATTTGGACTTCATCCGTGACTTCCATCAGCTAGTGAAGGCCCTACCCCAGTATCAGCACTCCTTCATTCCAGTACAATAGTTATAGCTGGAGTATTTACCCTCATCCGCTTTTATCCATTAATAGAAAACAACCTCAC
    ||||||||||||||||||||||||||||||||||||||||||||||||||||||||||||||||||||||||| |||   ||| |||||||||||||||||||||||||||||||||||||||||||||||||||||||||||||||||||||||||||||||||||||||||||||||||
R   ACTACTAGGTCTTCTCTTAGCCTCAGCAGGAAAATCAGCCCAATTTGGACTTCATCCGTGACTTCCATGAGCAGGTGTAGGCCCTACCCCAGTATCAGCACTCCTTCATTCCAGTACAATAGTTATAGCTGGAGTATTTACCCTCATCCGCTTTTATCCATTAATAGAAAACAACCTCAC
             | |   |     ||    |    |      |    |  | |  | |  |  ||||||||||||||||||| |    |     || |||      |  |          ||  ||  |        | |     |    |     |     |  |  |    |  | | |
P   TTGTATTTCTGTAAACAAGCACTTCCTGGCCGGACACTGGCCAGATCTAGATGAATCAGGCTATCCATGAGCAGGTGTAGGCCATCAATCTCATACACCACATTGAGTTGACCAATTTTGTGTTCGAGTAGTTTACGTAAACTGTGGCGCTGTCGGGGGAGGACCAGTGGAATATCCCCA
Pos                                                                     <-1544281--1544300->
R translated                                                        ...S  m  S  r  c  R  P...
```

$\Delta G^0_{min}$ = -18.73 kcal/mol

**MK89.**

```
                                                                          |-- net 51 bp loss ------------------------------------------------------|
A   ACTACTAGGTCTTCTCTTAGCCTCAGCAGGAAAATCAGCCCAATTTGGACTTCATCCGTGACTTCCATCAGCTAGTGAAGGCCCTACCCCAGTATCAGCACTCCTTCATTCCAGTACAATAGTTATAGCTGGAGTATTTACCCTCATCCGCTTTTATCCATTAATAGAAAACAACCTCAC
    |||||||||||||||||||||||||||||||||||||||||||||||||||||||||||||||||| ||||| |                                                  | |  ||||||| |||||||||||||||||||||||||||||||||||||||||
R   ACTACTAGGTCTTCTCTTAGCCTCAGCAGGAAAATCAGCCCAATTTGGACTTCATCCGTGACTTCAATCAGTTT--------------------------------------------------GCGTTTCAGTATTTTCCCTCATCCGCTTTTATCCATTAATAGAAAACAACCTCAC
         |      | || |  |   |               |          | |    ||||||||||||                                                  ||||||||||||||||||        | |     |     |  | || |  |  |
P   GGCTATCTCCTGTATCATGCCGAGTGAGCTTGGGTTGAAGAAGCCAAATGCGCTTTATAATCTTCAATCAGTTT--------------------------------------------------GCGTTTCAGTATTTTCCCTGTCGCTTCTCTTGGAAAGTTTGACAAAATTCAAAA
Pos                                                                     <-1564847----------------------------------------------------------------1564816->
R translated                                                        ...T  S  I  S  l                                                     r  f  S  I  Y  P...
```

$\Delta G^0_{min}$ = -8.77 kcal/mol (left) / -10.99 kcal/mol (right)

**MK143.**

```
                                                                            |---| no net gain/loss
A  ACTACTAGGTCTTCTCTTAGCCTCAGCAGGAAAATCAGCCCAATTTGGACTTCATCCGTGACTTCCATCAGCTAGTGAAGGCCCTACCCCAGTATCAGCACTCCTTCATTCCAGTACAATAGTTATAGCTGGAGTATTTACCCTCATCCGCTTTTATCCATTAATAGAAAACAACCTCAC
R  ACTACTAGGTCTTCTCTTAGCCTCAGCAGGAAAATCAGCCCAATTTGGACTTCATCCGTGACTTCCATCAGCGAGTTAAGGCCCTACCCCAGTATCAGCACTCCTTCATTCCAGTACAATAGTTATAGCTGGAGTATTTACCCTCATCCGCTTTTATCCATTAATAGAAAACAACCTCAC
P  CTCAACCAAAAATCAGGTTTCCAGCTTAAGACAGTCAAGGCAAAAACAACAAATACGAGCACTCCAGCCAGCGAGTTAAGCCATACCCACAATAAAGAAAAGGTATATGCATAATGGCAATTTCAACATCAGAACAAGATGATGTCGTCAGCGAGATCAATATTACCCCTTTAGTAGACG
Pos                                                                1582641<------------>1582654
R translated                                                         ...I  S  e  l  R  P...
```

$\Delta G^0_{min}$ = -11.87 kcal/mol

**MK15.**

```
                                                                    |-----------| net 3 bp loss
A  ACTACTAGGTCTTCTCTTAGCCTCAGCAGGAAAATCAGCCCAATTTGGACTTCATCCGTGACTTCCATCAGCTAGTGAAGGCCCTACCCCAGTATCAGCACTCCTTCATTCCAGTACAATAGTTATAGCTGGAGTATTTACCCTCATCCGCTTTTATCCATTAATAGAAAACAACCTCAC
R  ACTACTAGGTCTTCTCTTAGCCTCAGCAGGAAAATCAGCCCAATTTGGACTTCATCCGTGACTTTCAACATC-A--AAAGGCCCTACCCCAGTATCAGCACTCCTTCATTCCAGTACAATAGTTATAGCTGGAGTATTTACCCTCATCCGCTTTTATCCATTAATAGAAAACAACCTCAC
P  GATTGCATCAGGCTTACCAATCACCGCAGATTCAAGAACCTCTTGGCTCAAGTAAAGTACTTTCAACATC-A--AAAGGCCCTACACGATAACCCGATGTTGTAATTACATCATCTATACGCCCGACAAAGTCGATACTGCCAAACGCATTCATTTTCACAGTATCACCCGTAATATA
Pos                                                                <-1609947---------1609924->
R translated                                                    ...T  f  n  I   k    R  P  Y...
```

$\Delta G^0_{min}$ = -17.55 kcal/mol

**MK35.**

```
                                                                      |-- net 63 bp loss ---------------------------------------------------------------|
A  ACTACTAGGTCTTCTCTTAGCCTCAGCAGGAAAATCAGCCCAATTTGGACTTCATCCGTGACTTCCATCAGCTAGTGAAGGCCCTACCCCAGTATCAGCACTCCTTCATTCCAGTACAATAGTTATAGCTGGAGTATTTACCCTCATCCGCTTTTATCCATTAATAGAAAACAACCTCAC
R  ACTACTAGGTCTTCTCTTAGCCTCAGCAGGAAAATCAGCCCAATTTGGACTTCATCCGTGACTTCCCCCAGCGA---------------------------------------------------------------TCTGCCTTATCTGCTTTTATCCATTAATAGAAAACAACCTCAC
P  TTGTTTTTGCAGCTTTGGCCATAATCGGACCAAAATTTCTGGCGGCTTTACGATAAGTTAAATTCCCCCAGCGA---------------------------------------------------------------TCTGCCTTATCTGCATAAATGAATGCATAATCGGCTTCAAGTG
Pos                                                                  <-1701316------------------------------------------------------------------1701291->
R translated                                                    ...T  S  p  s  d                                                                 l  P  y  l  L...
```

$\Delta G^0_{min}$ = -8.12 kcal/mol (left) / -7.50 kcal/mol (right)

**A19.**

```
                                                                          |--------------------| net 6 bp loss
A  ACTACTAGGTCTTCTCTTAGCCTCAGCAGGAAAATCAGCCCAATTTGGACTTCATCCGTGACTTCCATCAGCTAGTGAAGGCCCTACCCCAGTATCAGCACTCCTTCATTCCAGTACAATAGTTATAGCTGGAGTATTTACCCTCATCCGCTTTTATCCATTAATAGAAAACAACCTCAC
R  ACTACTAGGTCTTCTCTTAGCCTCAGCAGGAAAATCAGCCCAATTTGGACTTCATCCGTGACTTCCATCAGTTATCGTCGTCACT------GGATCAGCACTCCTTCATTCCAGTACAATAGTTATAGCTGGAGTATTTACCCTCATCCGCTTTTATCCATTAATAGAAAACAACCTCAC
P  TGAAAGATGCGATGCATCAATGACTTCAAACTGATTGCATTGAATATGCTGTTGCATAAATTCACCATCAGTTATCGTCGTCACT------GGATCAGCACTACCCGCGATAATCAATGTCGGAATTGAGATTGATGCCAATTTTTCACGTACATCCGCTTTAGCCAAAGCACGGCAAGC
Pos                                                                    <-1713833--------------------1713802->
R translated                                                        ...S  I  S  y  r  r  h  w       I  S  T  P...
```

$\Delta G^0_{min}$ = -9.20 kcal/mol (left) / -11.56 kcal/mol (right)

**MK41.**

```
                                                                   |-- net 6 bp loss ------------------|
A  ACTACTAGGTCTTCTCTTAGCCTCAGCAGGAAAATCAGCCCAATTTGGACTTCATCCGTGACTTCCATCAGCTAGTGAAGGCCCTACCCCAGTATCAGCAC-TCCTTCATTCCAGTACAATAGTTATAGCTGGAGTATTTACCCTCATCCGCTTTTATCCATTAATAGAAAACAACCTCAC
R  ACTACTAGGTCTTCTCTTAGCCTCAGCAGGAAAATCAGCCCAATTTGGACTTCATCCGTGACTTC-A-AAGCGAG-G--TGCCTTA--GTATTTTCACCAAGTCCTTCATTCCAGTACAATAGTTATAGCTGGAGTATTTACCCTCATCCGCTTTTATCCATTAATAGAAAACAACCTCAC
P  TTATCTGCAATACGGCTTCCCACGACAATCGTTGTACCTGCAATGGTAGGTGCCGAAGTGACTTC-A-AAGCGAG-G--TGCCTTA--GTATTTTCACCAAGTCCTTCATGTAAATTCACTGTACCATTAACACCAAAGTCTTTACAACGAGCGCCTGTATCTGCATTTACTGCAATCAAA
Pos                                                           <-1722199-----------------------------------1722154->
R translated                                               ...V  T  S    k  r  g     a  l     v  f  s  p  s  P  S  F...
```

$\Delta G^0_{min}$ = -9.00 kcal/mol (left) / -8.90 kcal/mol (right)

**R17.**

```
                                                                      |-- net 3 bp loss ---------|
A  ACTACTAGGTCTTCTCTTAGCCTCAGCAGGAAAATCAGCCCAATTTGGACTTCATCCGTGACTTCCATCAGC-TAGTGAAGGCCCTACCCCAGTATCAGCACTCCTTCATTCCAGTACAATAGTTATAGCTGGAGTATTTACCCTCATCCGCTTTTATCCATTAATAGAAAACAACCTCAC
R  ACTACTAGGTCTTCTCTTAGCCTCAGCAGGAAAATCAGCCCAATTTGGACTTCATCCGTGACTTCCACCTGCGTAGTAAAGACCATTT----GTTTCAGCACTCCTTCATTCCAGTACAATAGTTATAGCTGGAGTATTTACCCTCATCCGCTTTTATCCATTAATAGAAAACAACCTCAC
P  GTATAAATTATTAACATCCAGATAAAATAATCCTAAACGTAAAGCCGGACTTACATTGTAATAGCCACCTGCGTAGTAAAGACCATTT----GTTTCAGCACTTTTGAAACGCCCATTCACCGCTGAAATCCTGATATTTTCATAATTGGTCGAGTCGCGCTGATTCACCTTGTCTATAAA
POS                                                                    <-1730059---------------------1730093->
R translated                                                        ...S  t  c  v  v  k  t  i  c     f  S  T...
```

$\Delta G^0_{min}$ = -10.32 kcal/mol (left) / -10.58 kcal/mol (right)

**A60.**

```
                                                                              |-| no net gain/loss
A   ACTACTAGGTCTTCTCTTAGCCTCAGCAGGAAAATCAGCCCAATTTGGACTTCATCCGTGACTTCCATCAGCTAGTGAAGGCCCTACCCCAGTATCAGCACTCCTTCATTCCAGTACAATAGTTATAGCTGGAGTATTTACCCTCATCCGCTTTTATCCATTAATAGAAAACAACCTCAC
R   ACTACTAGGTCTTCTCTTAGCCTCAGCAGGAAAATCAGCCCAATTTGGACTTCATCCGTGACTTCCATCAGCTATTTAAGGCCCTACCCCAGTATCAGCACTCCTTCATTCCAGTACAATAGTTATAGCTGGAGTATTTACCCTCATCCGCTTTTATCCATTAATAGAAAACAACCTCAC
P   TATGGTTTTTTGTTTGCTTTTAAAATTATATTTATTTTACTTAGGCTATTTTATTTTTTAATTTGCTTAGGCTATTTAAGGCTACAAAGTAAATGAATCGATTTGCCCAAAAGTCAAAGTAAACTTGCAAAAAAGACGCAGATAATCAAACAACGGATGTCTCCTAACCAAATTAATGAA
Pos                                                                           <----------> 1747667-1747656
R translated                                                              ...S  y  l  R  P...
```

$\Delta G^0_{min}$ = -9.87 kcal/mol

**MK85.**

```
                                                                            |-- net 78 bp loss ---------------------------------------------------------------------------|
A   ACTACTAGGTCTTCTCTTAGCCTCAGCAGGAAAATCAGCCCAATTTGGACTTCATCCGTGACTTCCATCAGCTAGTGAAGGCCCTACCCCAGTATCAGCACTCCTTCATTCCAGTACAATAGTTATAGCTGGAGTATTTACCCTCATCCGCTTTTATCCATTAATAGAAAACAACCTCAC
R   ACTACTAGGTCTTCTCTTAGCCTCAGCAGGAAAATCAGCCCAATTTGGACTTCATCCGTGACTTCCATCAG------------------------------------------------------------------------------AATTACGACAACAAACCGAAAACAACCTCAC
P   GAAAGAAATCTTAAAATCCATTATTGATGACCTGTGTGGTATTTTGTTTCCAATGCGGCTTGGCCCATCAG------------------------------------------------------------------------------AATTACGACAACAAACCGAAAACTATTATAT
Pos                                                                 <-1768627------------------------------------------------------------------------------------------1768598->
R translated                                                    ...S  I  r                                                                                        i  t  t  t  n  r  K  Q...
```

$\Delta G^0_{min}$ = -7.58 kcal/mol (left) / -5.21 kcal/mol (right)

**MK46.**

```
                                                                             |---| no net gain/loss
A   ACTACTAGGTCTTCTCTTAGCCTCAGCAGGAAAATCAGCCCAATTTGGACTTCATCCGTGACTTCCATCAGCTAGTGAAGGCCCTACCCCAGTATCAGCACTCCTTCATTCCAGTACAATAGTTATAGCTGGAGTATTTACCCTCATCCGCTTTTATCCATTAATAGAAAACAACCTCAC
R   ACTACTAGGTCTTCTCTTAGCCTCAGCAGGAAAATCAGCCCAATTTGGACTTCATCCGTGACTTCCATCAGCCACTTAAGGCCCTACCCCAGTATCAGCACTCCTTCATTCCAGTACAATAGTTATAGCTGGAGTATTTACCCTCATCCGCTTTTATCCATTAATAGAAAACAACCTCAC
P   TTGAAATGAGTAATGGTTCATCTGTCGAACATAAAGAACTTGCCTCTAAAGCATTTGAAATTTTGACACAGCCACTTAAGGCCTAATACTTAAAACTAGCCCAAAGAAGAGTCGAGAGCTTTGGGCTAAATTTTTGCAATAATATAGAGGTGAAATATGAAGTTTGCCACAATAAAAAAT
Pos                                                                1816041<------------->1816055
R translated                                                      ...I  S  h  l  S  P...
```

$\Delta G^0_{min}$ = -14.01 kcal/mol

**MK51.**

```
                                                                         |--------------------------| no net gain/loss
A   ACTACTAGGTCTTCTCTTAGCCTCAGCAGGAAAATCAGCCCAATTTGGACTTCATCCGTGACTTCCATCAGCTAGTGAAGGCCCTACCCCAGTATCAGCACTCCTTCATTCCAGTACAATAGTTATAGCTGGAGTATTTACCCTCATCCGCTTTTATCCATTAATAGAAAACAACCTCAC
R   ACTACTAGGTCTTCTCTTAGCCTCAGCAGGAAAATCAGCCCAATTTGGACTTCATCCGTGACTTCCACCAGAACCGCCAGCACCACCACCAGCGCCAGCACTCCTTCATTCCAGTACAATAGTTATAGCTGGAGTATTTACCCTCATCCGCTTTTATCCATTAATAGAAAACAACCTCAC
P   ACCACCTGAGCCTCCAGCACCACCAGATCCACCTGCACCGCCATTACCACCGCCTGACCCAGAACCACCAGAACCGCCTGCACCACCACCAGCGCCAGCACCGCCATTGGCACCCGAACCACCATCACCGCCAGAACCGCCATTAGCACCGACACAGTTAATAGTATTGCCTGAAACTGT
Pos                                                                 <-1854121-------------------1854157->
R translated                                                    ...S  t  r  t  a  c  t  t  t  S  a  S  T...
```

$\Delta G^0_{min}$ = -15.14 kcal/mol

**A6.**

```
                                                                             |----| no net gain/loss
A   ACTACTAGGTCTTCTCTTAGCCTCAGCAGGAAAATCAGCCCAATTTGGACTTCATCCGTGACTTCCATCAGCTAGTGAAGGCCCTACCCCAGTATCAGCACTCCTTCATTCCAGTACAATAGTTATAGCTGGAGTATTTACCCTCATCCGCTTTTATCCATTAATAGAAAACAACCTCAC
R   ACTACTAGGTCTTCTCTTAGCCTCAGCAGGAAAATCAGCCCAATTTGGACTTCATCCGTGACTTCCATCAGCCATAGCAGGCCCTACCCCAGTATCAGCACTCCTTCATTCCAGTACAATAGTTATAGCTGGAGTATTTACCCTCATCCGCTTTTATCCATTAATAGAAAACAACCTCAC
P   ATCTGTATTTGGACATGAAGAATCTTTGACAGGGTCACCAAAGACTAGATGCATCGATGGGTCATCACCAGCCATAGCAGGCCCTTTGTATCTACAACCTTGATAACATGCTGTTTGTGATGAGATTGCTTGAGAATAAGATACAGTTTGTTCATATGTCCAAGAATGCCAGCCAGGGAC
Pos                                                                  <-1854425--1854444->
R translated                                                    ...S  I  S  h  s  R  P  Y...
```

$\Delta G^0_{min}$ = -16.79 kcal/mol

**O140.**

```
                  |-- net 129 bp loss ------------------------------------------------------------------------------------------------------------------------------|
A   GGTCTTCTCT-TAGCCTCAGCAGGAAAATCAGCCCAATTTGGACTTCATCCGTGACTTCCATCAGCTAGTGAAGGCCCTACCCCAGTATCAGCACTCCTTCATTCCAGTACAATAGTTATAGCTGGAGTATTTACCCTCATCCGCTTTTATCCATTAATAGAAAACAACCTCACTATTCAA
R   GGTCTTCTCT-TAGCCCCAACAGGTTAGCC---------------------------------------------------------------------------------------------------------------------------------AGAAAACAACCTCACTATTCAA
P   GTAGGGCTCTATAGCCCCAACAGGTTAGCC---------------------------------------------------------------------------------------------------------------------------------AGAAAACAACCACCTCAGTTAT
Pos       <-1855272--------------------------------------------------------------------------------------------------------------------------------------------------1855306->
R tl ...S  L   S  p  n  R  l  a                                                                                                                                      R  K  Q  P...
```

$\Delta G^0_{min}$ = -17.87 kcal/mol (left) / -14.05 kcal/mol (right)

**MK97.**

```
                                                                       |-- net 6 bp loss -------|
A  ACTACTAGGTCTTCTCTTAGCCTCAGCAGGAAAATCAGCCCAATTTGGACTTCATCCGTGACTTCCATCAGCTAGTGAAGGCCCTACCCCAGTATCAGCACTCCTTCATTCCAGTACAATAGTTATAGCTGGAGTATTTACCCTCATCCGCTTTTATCCATTAATAGAAAACAACCTCAC
R  ACTACTAGGTCTTCTCTTAGCCTCAGCAGGAAAATCAGCCCAATTTGGACTTCATCCGTGACTTCCATC--C-A-TG-TCGCAATA-CGCACTAGCAGCACTCCTTCATTCCAGTACAATAGTTATAGCTGGAGTATTTACCCTCATCCGCTTTTATCCATTAATAGAAAACAACCTCAC
P  ATTAATAATGCAATCGATTGTAAACTCTTTGAAATTACGAAAATTGCCAACCAGATGCACGCTTCCATC--C-A-TG-TCGCAATA-CGCACTAGCAGCATCCTTTTGCCAGTCTTATCTGCCACCCAGCCCCATATAAATGGCGCAAAAAATCGTGTAATGATGGCAATCGAAGAAAGT
Pos                                                                <-1865765---------------------1865802->
R translated                                                  ...T  S  I   h   v   a  I   r  t  s  S  T...
```

$\Delta G^0_{min}$ = -8.90 kcal/mol (left) / -12.17 kcal/mol (right)

**MK105.**

```
              |-- net 93 bp loss -----------------------------------------------------------------------------------------------------------|
A  TTTGGACTTCATCCGTGACTTCCATCAGCTAGTGAAGGCCCTACCCCAGTATCAGCACTCCTTCATTCCAGTACAATAGTTATAGCTGGAGTATTTACCCTCATCCGCTTTTATCCATTAATAGAAAACAACCTCACTATTCAAACTTCAACACTTCTGGTTCTGGCTCTAATGTCAC
R  TTTGGACTTCAGCCGTTACCATGCGACTTTGGACAGAATGCTTT-------------------------------------------------------------------------------------------CCTGTCGCTGGAGGGCTGATTGGCTCTGACTCTAATGTCAC
P  GGCATACTTCAGCCGTTACCATGCGACTTTGGACAGAATGCTTT-------------------------------------------------------------------------------------------CCTGTCGCTGGAGGGCTGATTGGCTCTGACTCTAATTTACG
Pos     <-1868500----------------------------------------------------------------------------------------------------------------------------------1868576->
R tl ...T  S  a  V  T  m  r  l  w  t  e  c  f                                                                                       p  v  a  g  g  l  i  G  S  d  S  N  V...
```

$\Delta G^0_{min}$ = -14.75 kcal/mol (left) / -16.94 kcal/mol (right)

**MK90.**

```
                                                                          |-- net 9 bp loss ------------------|
A  ACTACTAGGTCTTCTCTTAGCCTCAGCAGGAAAATCAGCCCAATTTGGACTTCATCCGTGACTTCCATCAGCTAGTGAAGGCCCTACCCCAGTATCAGCACTCCT-TC-ATTCCAGTACAATAGTTATAGCTGGAGTATTTACCCTCATCCGCTTTTATCCATTAATAGAAAACAACCTCAC
R  ACTACTAGGTCTTCTCTTAGCCTCAGCAGGAAAATCAGCCCAATTTGGACTTCATCCGTGACTTCCATCAGC-AGTGAA-----TA-CTTTG-AT--GC-CAGCTATCAATTCCAGTACAATAGTTATAGCTGGAGTATTTACCCTCATCCGCTTTTATCCATTAATAGAAAACAACCTCAC
P  AATCACCGCATGCATAACAAATTGATTATTGCGGACAACACAATTGCGGTGACTGGAGGACGCAATATCAGC-AGTGAA-----TA-CTTTG-AT--GC-CAGCTATCAATTCCAGTTTACAGATCTCGATATTTTGTTTTATGGTACAGCCGCATCTCAGGCAGAGAAAGTCTTTCAAACT
Pos                                                                     <-1874200--------------------------------1874239->
R translated                                                      ...I  S  s  e      y  f  d     a  s  y  q  F  Q  Y...
```

$\Delta G^0_{min}$ = -13.69 kcal/mol (left) / -8.91 kcal/mol (right)

**A100.**

```
                                                                       |-- net 57 bp loss -------------------------------------|
A  ACTACTAGGTCTTCTCTTAGCCTCAGCAGGAAAATCAGCCCAATTTGGACTTCATCCGTGACTTCCATCAGCTAGTGAAGGCCCTACCCCAGTATCAGCACTCCTTCATTCCAGTACAATAGTTATAGCTGGAGTATTTACCCTCATCCGCTTTTATCCATTAATAGAAAACAACCTCAC
R  ACTACTAGGTCTTCTCTTAGCCTCAGCAGGAAAATCAGCCCAATTTGGACTTCATCCGTGACTTCCA---------------------------------------------------------ATAGCTGGAGTATTTACCCTCATCCGCTTTTATCCATTAATAGAAAACAACCTCAC
P  TCTTCGGTTGGATACATCGCATCATGAGTATTAATCTGAGTTTGCGTCACCTGACTGATGACTTCCA---------------------------------------------------------ATAGCTGTTGTATAGAGGTTACCTCATTGTGTGCAATATTAAATGATTCACAACAA
Del                                                             <-1883762-------------------------------------------------------------1883783->
R translated                                                 ...V  T  S  n                                                          S  W  S  I...
```

$\Delta G^0_{min}$ = -11.57 kcal/mol (left) / -9.12 kcal/mol (right)

**A26.**

```
                                                                       |--------| no net gain/loss
A  ACTACTAGGTCTTCTCTTAGCCTCAGCAGGAAAATCAGCCCAATTTGGACTTCATCCGTGACTTCCATCAGCTAGTGAAGGCCCTACCCCAGTATCAGCACTCCTTCATTCCAGTACAATAGTTATAGCTGGAGTATTTACCCTCATCCGCTTTTATCCATTAATAGAAAACAACCTC
R  ACTACTAGGTCTTCTCTTAGCCTCAGCAGGAAAATCAGCCCAATTTGGACTTCATCCGTGACTTCCACCTGCAGCATAAGGCCCTACCCCAGTATCAGCACTCCTTCATTCCAGTACAATAGTTATAGCTGGAGTATTTACCCTCATCCGCTTTTATCCATTAATAGAAAACAACCTC
P  ATTGACGCCTCCCTGAGCATTTACAGCTTTCATATAATCGATAAATCCACCAAAAAAGCCCGTTCCACCTGCAGCATAAGGCCCTACACGATAACTTTGTAATGGAAAATATTGAGCATTTTTATCAGACAAAACTTTTTCAGCGTTCGATTTATCTGTGTTTGTTTGATCTTGAGAC
Pos                                                                <-1928206---------1928232->
R translated                                                  ...T  S  t  c  s  i  R  P  Y...
```

$\Delta G^0_{min}$ = -16.91 kcal/mol

**MK30.**

```
                                                                  |-- net 9 bp loss ---------------------|
A  ACTACTAGGTCTTCTCTTAGCCTCAGCAGGAAAATCAGCCCAATTTGGACTTCATCCGTGACTTCCATCAGCTAGTGAAGGCCCTACCCCAGTATCAGCACTCCTTCATTCCAGTACAATAGTTATAGCTGGAGTATTTACCCTCATCCGCTTTTATCCATTAATAGAAAACAACCTCAC
R  ACTACTAGGTCTTCTCTTAGCCTCAGCAGGAAAATCAGCCCAATTTGGACTTCATCCGTGAC---------CACAGCATCTGTTTCGGCCCAGTTCGGCACACCTTCATTCCAGTACAATAGTTATAGCTGGAGTATTTACCCTCATCCGCTTTTATCCATTAATAGAAAACAACCTCAC
P  AAAGGCTGTCCTTCGGTTAAACGGATTTCAATTGGTGGTTGATCTGGCACATAAATCGTGAC---------CACAGCATCTGTTTCGGCCCAGTTCGGCACACCTTCATAAATAAATGCAAACAACACCACACGGCGAATCTGCGACCAGTGCTGTCCATTAATCCTAAGCCACTCACCC
Pos                                                          <-1944867-----------------------------------1944910->
R translated                                            ...S  V  T          t  a  s  v  s  a  q  f  g  T  P  S  F...
```

$\Delta G^0_{min}$ = -7.18 kcal/mol (left) / -16.21 kcal/mol (right)

```
MK28.                                |-- net 42 bp loss ----------------------------------------|
A  ACTACTAGGTCTTCTCTTAGCCTCAGCAGGAAAATCAGCCCAATTTGGACTTCATCCGTGACTTCCATCAGCTAGTGAAGGCCCTACCCCAGTATCAGCACTCCTTCATTCCAGTACAATAGTTATAGCTGGAGTATTTACCCTCATCCGCTTTTATCCATTAATAGAAAACAACCTCAC
   |||||||||||||||||||||||||||||||||| |                                                 |  | ||| ||||||||||||||||||||||||||||||||||||||||||||||||||||||||||||||||||||||||||||||||||||||
R  ACTACTAGGTCTTCTCTTAGCCTCAGCAGGAAAAACTTTT------------------------------------------ATCAATCTAGTTTCAGCACTCCTTCATTCCAGTACAATAGTTATAGCTGGAGTATTTACCCTCATCCGCTTTTATCCATTAATAGAAAACAACCTCAC
     || | |    |     ||| ||  ||||||||||||||                                          |||||||||||||||||||| |      |  ||          |   |   ||  | ||    |         ||   | |  ||  |   |  |  |
P  TGTATTGGTCGATGATGAAGCTTCTACAGGAAAAACTTTT------------------------------------------ATCAATCTAGTTTCAGCACTACAAGCGGCTGGTCTTGATCAAATGTCTCATGTCGTAACTGCAACTTTGGCGGATTGGTCATCAGGTATTCATATGGC
Pos                  <-1949398--------------------------------------------------------------------1949355->
R translated      ...S  L  S  R  K  n  f  y                                             q  s  S  f  S  T  P...
```
$\Delta G^0_{min}$ = -13.62 kcal/mol (left) / -12.85 kcal/mol (right)

```
MK37.                                                                  |---------| no net gain/loss
A  ACTACTAGGTCTTCTCTTAGCCTCAGCAGGAAAATCAGCCCAATTTGGACTTCATCCGTGACTTCCATCAGCTAGTGAAGGCCCTACCCCAGTATCAGCACTCCTTCATTCCAGTACAATAGTTATAGCTGGAGTATTTACCCTCATCCGCTTTTATCCATTAATAGAAAACAACCTCAC
   ||||||||||||||||||||||||||||||||||||||||||||||||||||||||||||||||||| |  ||| || ||||||||||||||||||||||||||||||||||||||||||||||||||||||||||||||||||||||||||||||||||||||||||||||||||||||
R  ACTACTAGGTCTTCTCTTAGCCTCAGCAGGAAAATCAGCCCAATTTGGACTTCATCCGTGACTTCCAACTTCTATTGCAGGCCCTACCCCAGTATCAGCACTCCTTCATTCCAGTACAATAGTTATAGCTGGAGTATTTACCCTCATCCGCTTTTATCCATTAATAGAAAACAACCTCAC
      | |   |         || |   || |       ||    |           |    | ||||||||||||||||||             | ||      |          |         |  ||      |       | |            ||   |             |
P  GTCAATCCATGGCCGTGCTGCATGTACACGTTTTAAGGCTTCTTCAAATGCAGCACGAGCAATTCCAACTTCTATTGCAGCATGCATAAGTTGAGCAAATGGTCCGACCAAAGTTGGGTGTTGAAATGCACTGTCAAAAGGAATGACATCTTCGGCAGCAACGAATACCTGAGTGAACTT
Pos                                                        2050271<----------------->2050288
R translated                                                ...T  S  n  f  y  c  R...
```
$\Delta G^0_{min}$ = -8.53 kcal/mol

```
AD41.                                                                     |--------| no net gain/loss
A  ACTACTAGGTCTTCTCTTAGCCTCAGCAGGAAAATCAGCCCAATTTGGACTTCATCCGTGACTTCCATCAGCTAGTGAAGGCCCTACCCCAGTATCAGCACTCCTTCATTCCAGTACAATAGTTATAGCTGGAGTATTTACCCTCATCCGCTTTTATCCATTAATAGAAAACAACCTCAC
   |||||||||||||||||||||||||||||||||||||||||||||||||||||||||||||||||||||| | | || | ||||||||||||||||||||||||||||||||||||||||||||||||||||||||||||||||||||||||||||||||||||||||||||||||||||
R  ACTACTAGGTCTTCTCTTAGCCTCAGCAGGAAAATCAGCCCAATTTGGACTTCATCCGTGACTTCCATCATCGACTGTATGCCCTACCCCAGTATCAGCACTCCTTCATTCCAGTACAATAGTTATAGCTGGAGTATTTACCCTCATCCGCTTTTATCCATTAATAGAAAACAACCTCAC
   |  |   |  |  |  |     |||  |    | |     || ||||          ||| | | |||||||||||||||||| |   |   ||              |               |    |     |  |          |  || |  ||| |    |||  |
P  ATGAAATAGCCCGCCATATCAGTCATGACTGCATTGGTTGCATTTTGCCAAGTGAGGGTGCCATTCATCATCGACTGTATGCCTTTATCCCATAGGCTTGAAATACGGTCAGTAAGTCCACCAAACTTTTTCTCGAAGTCTTTCATCTGGGCTTCAGACATAAGATTAAACGATTGCGTA
Pos                                                          <-2139062--------2139087->
R translated                                          ...S  V  T  S  I  i  d  c  m  P  Y...
```
$\Delta G^0_{min}$ = -15.19 kcal/mol

```
A17.                                                                        |--------------| net 9 bp loss
A  ACTACTAGGTCTTCTCTTAGCCTCAGCAGGAAAATCAGCCCAATTTGGACTTCATCCGTGACTTCCATCAGCTAGTGAAGGCCCTACCCCAGTATCAGCACTCCTTCATTCCAGTACAATAGTTATAGCTGGAGTATTTACCCTCATCCGCTTTTATCCATTAATAGAAAACAACCTCAC
   |||||||||||||||||||||||||||||||||||||||||||||||||||||||||||||||||||||||||  |||||       ||  |||||||||||||||||||||||||||||||||||||||||||||||||||||||||||||||||||||||||||||||||||||||||
R  ACTACTAGGTCTTCTCTTAGCCTCAGCAGGAAAATCAGCCCAATTTGGACTTCATCCGTGACTTCCATCAGCT--TGAAG-------CCGTGTATCAGCACTCCTTCATTCCAGTACAATAGTTATAGCTGGAGTATTTACCCTCATCCGCTTTTATCCATTAATAGAAAACAACCTCAC
      |    | |   ||      | |   ||   ||  || |  | |   |           |    |||||||  |||||       |||||||||||||||   |      |        |      |  |                 ||         |||  |    |
P  CAAATATTGGCACATCGAGAGATAAAGCGGCCCATGGGCACTTTGTTCGCCGTGATATACCCGATGATCAGCT--TGAAG-------CCGTGTATCAGCACTGGCTGGCAAAACGTATTGTGAACCGCCCAGCAAGCGACATGCTGCGTGCCGGCTGGTTTTATGAAGGCATTCAGGATG
Pos                                                                   <-2152623------------------2152597->
R translated                                                      ...I  S  l    k         P  c  I  S  T  P...
```
$\Delta G^0_{min}$ = -13.48 kcal/mol (left) / -15.82 kcal/mol (right)

```
MK6.            |-- net 123 bp loss ----------------------------------------------------------------------------------------------------------------------------|
A  CAGGAAAATCAGCCCAATTTGGACTTCATCCGTGACTTCCATCAGCTAGTGAAGGCCCTACCCCAGTATCAGCACTCCTTCATTCCAGTACAATAGTTATAGCTGGAGTATTTACCCTCATCCGCTTTTATCCATTAATAGAAAACAACCTCACTATTCAAACTTCAACACTTCTGGTTCTG
   ||||||||||||| |||                                                                                                                            |||||     |    ||||||||||||||||||||||||||
R  CAGGAAAATCAGCGCAA---------------------------------------------------------------------------------------------------------------------------AAAAACCTATGCCGCGTTCAAACTTCAACACTTCTGGTTCTG
       |||||||||||||                                                                                                                           |||||||||||||||||||||||||||  | |      |
P  AGCAAAAATCAGCGCAA---------------------------------------------------------------------------------------------------------------------------AAAAACCTATGCCGCGTTCAAACTTCATAAATCTGTCATAAA
Pos    <-2204361--------------------------------------------------------------------------------------------------------------------------------------------------2204322->
R tl...K  I  S  a  k                                                                                                                              K  p  m  p  r  S  N  F  N...
```
$\Delta G^0_{min}$ = -15.68 kcal/mol (left) / -13.55 kcal/mol (right)

```
MK61.                                                                       |---| no net gain/loss
A  ACTACTAGGTCTTCTCTTAGCCTCAGCAGGAAAATCAGCCCAATTTGGACTTCATCCGTGACTTCCATCAGCTAGTGAAGGCCCTACCCCAGTATCAGCACTCCTTCATTCCAGTACAATAGTTATAGCTGGAGTATTTACCCTCATCCGCTTTTATCCATTAATAGAAAACAACCTCAC
   |||||||||||||||||||||||||||||||||||||||||||||||||||||||||||||||||||||||| ||| |||||||||||||||||||||||||||||||||||||||||||||||||||||||||||||||||||||||||||||||||||||||||||||||||||||||
R  ACTACTAGGTCTTCTCTTAGCCTCAGCAGGAAAATCAGCCCAATTTGGACTTCATCCGTGACTTCCATCAGCCAGTCAAGGCCCTACCCCAGTATCAGCACTCCTTCATTCCAGTACAATAGTTATAGCTGGAGTATTTACCCTCATCCGCTTTTATCCATTAATAGAAAACAACCTCAC
          |             ||         |                       |    ||    ||||||||||||        |      ||      |     |    ||  || |    |         |        |  | |      | ||     ||      |  ||
Ab CAGCAATGAGACGTCTAGCAGCTTTAGTTTTGACATCAGGTTTGACTATTACGGGTCTCAGCTATGCTCAGCCAGTCAATCAATCTGCACTTATTCCAACAACAAGTGTGAGTGTGAAAGATGCATTAACATTCAAAGATAATACGCCTGTAAAACTTCAAGGTCAGGTGGTTAAGTCCT
Pos                                                             2211010<---------->2210998
R translated                                                      ...I  S  q  s  R...
```
$\Delta G^0_{min}$ = -10.48 kcal/mol

**MK18.**

```
                                                                                                                                                     |---------| net 1 bp inserted
A  AATTTGGACTTCATCCGTGACTTCCATCAGCTAGTGAAGGCCCTACCCCAGTATCAGCACTCCTTCATTCCAGTACAATAGTTATAGCTGGAGTATTTACCCTCATCCGCTTTTATCCATTAATAGAAAACAACCTCACTATT-CAAACTTCAACACTTCTGGTTCTGGCTCTAATGTCA
R  AATTTGGACTTCATCCGTGACTTCCATCAGCTAGTGAAGGCCCTACCCCAGTATCAGCACTCCTTCATTCCAGTACAATAGTTATAGCTGGAGTATTTACCCTCATCCGCTTTTATCCATTAATAGAAAACAACCTCACAATGACAACCATCAACACTTCTGGTTCTGGCTCTAATGTCA
P  ATACTCTTCCTTGATCAATAGACTCCTTATTGATTTTTTGTAGATCAAGCAAAAAAAAGTCTCTACTTTGCATGGCTTTTCGCTTATCTACACCACTGATATTTAAAAAAATGAGTGATGCCAAAATACCTACAATCACAATGACAACCATCAACTCAACAAGCGTAAAGCCAGACTGAC
Pos                                                                                                                                                 <-2317653--2317672->
R translated                                                                                                                                            m  t  t  i  N...
```

$\Delta G^0_{min}$ = -7.37 kcal/mol

**MK17.**

```
                                                   |-- net 162 bp loss ------------------------------------------------|
A  CTACACTCCCACTACTAGGTCTTCTCTTAGCCTCAGCAGGAAAATCAGCCCACCAATTTGGACTTCATCCGTGACTTCCATCAGCTAGTGAAGG(120xN)TCTGGCTCTAATGTCACTAATGAAAAAATGCGTTTCTGGAGTCCAGAGGTTCGTGAGTTAGAGCCTTATGTACCTGGTG
R  CTACACTCCCACTACTAGGTCTTCTCTTAGCCTCAGCAGGAAAATCAGCCCAC-------------------------------------------------TGTGTACCCTCAAGCAACGTGAAAAAATGCGTTTCTGGAGTCCAGAGGTTCGTGAGTTAGAGCCTTATGTACCTGGTG
P  TCAAAGTATCAAGATGGGCTTACCCGATGATATGCAAACCGAATACGGCCCAC-------------------------------------------------TGTGTACCCTCAAGCAACGTGAAAAAATCCAGCAAGTGGTGCAGCGTTCAGTTGAACAAGGTGCAAAACTGATCACTG
Pos                                             <-2502313---------------------------------------------------------------------2502276->
R translated                                 ...I  S  P  f                                                  c  t  l  k  q  r  E  K  M  R...
```

$\Delta G^0_{min}$ = -9.25 kcal/mol (left) / -9.95 kcal/mol (right)

**MK40.**

```
                                                                        |-- no net gain/loss -------|
A  ACTACTAGGTCTTCTCTTAGCCTCAGCAGGAAAATCAGCCCAATTTGGACTTCATCCGTGACTTCCATCAGCTAGTGAAGGCCCTACCCCAGTATCAGCACTC-CTTCATTCCAGTACAATAGTTATAGCTGGAGTATTTACCCTCATCCGCTTTTATCCATTAATAGAAAACAACCTCAC
R  ACTACTAGGTCTTCTCTTAGCCTCAGCAGGAAAATCAGCCCAATTTGGACTTCATCCGTGACTTCCTTCAGCACCTGCCTGCTCACATACCAAAGCAGCACTC-CTTCATTCCAGTACAATAGTTATAGCTGGAGTATTTACCCTCATCCGCTTTTATCCATTAATAGAAAACAACCTCAC
P  ACGCATACGACGCACATCATCATCCTGAATGTGACGACGGTCTTCACGTAGATGCAAAGTAATTCCTTCAGCACCTGCCTGCTCACATACCAAAGCAGCACTCACTGGATCTGGATAACGTGTTCCTCTCGCCTGTCTCAAGGTCGCTACATGATCAATATTGACACCAAGCAATGCAGC
Pos                                                               <-2536867-------------------------2536910->
R translated                                                  ...T  S  f  S  t  c  l  l  t  y  q  s  S  T  P...
```

$\Delta G^0_{min}$ = -9.56 kcal/mol (left) / -12.28 kcal/mol (right)

**MK21.**

```
                |-- net 123 bp loss ------------------------------------------------------------------------------------------------|
A  TCAGCCCAATTTGGACTTCATCCGTGACTTCCATCAGCTAGTGAAGGCCCTACCCCAGTATCAGCACTCCTTCATTCCAGTACAATAGTTATAGCTGGAGTATTTACCCTCATCCGCTTTTATCCATTAATAGAAAACAACCTCACTATTCAAACTTCAACACTTCTGGTTCTGGCTCTAA
R  TCAGCCCAATTTGGACGTC-------------------------------------------------------------------------------------------------------CGACAATAGCGACAAAACCACTTCTGGTTCTGGCTCTAA
P  TTTCCAACATTTGGACGTC-------------------------------------------------------------------------------------------------------CGACAATAGCGACAAAACCACTTCTGAAGTCGGAAGGAA
Pos      <-2538818------------------------------------------------------------------------------------------------------------------2538854->
R tl    ...I  W  T  S                                                                                                                  d  n  s  d  k  t  T  S  G...
```

$\Delta G^0_{min}$ = -11.23 kcal/mol (left) /  -9.51 kcal/mol (right)

**R4.**

```
                                                                          |-- no net gain/loss -------|
A  ACTACTAGGTCTTCTCTTAGCCTCAGCAGGAAAATCAGCCCAATTTGGACTTCATCCGTGACTTCCATCAGCTAGTGAAGGCCCTACC-CCAGTATCAGCACTCCTTCATTCCAGTACAATAGTTATAGCTGGAGTATTTACCCTCATCCGCTTTTATCCATTAATAGAAAACAACCTCA
R  ACTACTAGGTCTTCTCTTAGCCTCAGCAGGAAAATCAGCCCAATTTGGACTTCATCCGTGACTTCCATCCGC-AGTTTCGACCAAAATACCACGATCTGCACTCCTTCATTCCAGTACAATAGTTATAGCTGGAGTATTTACCCTCATCCGCTTTTATCCATTAATAGAAAACAACCTCA
P  CAATTCTGGTTTTTCTTGTTCTACAATCCCTTTTAGAATTTTAGCAACTTCTAAAGCACCAATTTCATCCGC-AGTTTCGACCAAAATACCACGATCTGCACCTAGAGCCATTGAGGAACGGATTTGTTCCTGTGCTTCTTTAGGACCGATAGAAACCACAACGATTTCAGATACTGTTC
Pos                                                                  <-2611050----------------------2611088->
R translated                                                  ...T  S  I  r  s  f  d  q  i  t  t  I  c  T...
```

$\Delta G^0_{min}$ = -10.01 kcal/mol (left) / -9.03 kcal/mol (right)

**MK31.**

```
                                                                        |-- net 48 bp loss ---------------------------------------------------------------|
A  ACTACTAGGTCTTCTCTTAGCCTCAGCAGGAAAATCAGCCCAATTTGGACTTCATCCGTGACTTCCATCAGCTAGTGAAGGCCCTACCCCAGTATCAGCACTCCTTCATTCCAGTACAATAGTTATAGCTGGAGTATTTACCCTCATCCGCTTTTATCCATTAATAGAAAACAACCTCAC
R  ACTACTAGGTCTTCTCTTAGCCTCAGCAGGAAAATCAGCCCAATTTGGACTTCATCCGTGACTTCCAGCAGCTGCACCTGCGGCCAAT-------------------------------------------------CAATCCAATTTCCAGTTTTATCCATTAATAGAAAACAACCTCAC
P  AATGAACTTTCAGCGTTATGTCGCACCAACCAATGGTATGGCAACGGGAGAGTTAAATCAGGCACCAGCAGCTGCACCTGCGGCCAAT-------------------------------------------------CAATCCAATTTCCAGTTTTATCCATTACAAAATCCGCAGTTACT
Pos                                                                 <-2638447--------------------------------------------------------------------------------2638497->
R translated                                                    ...S  s  s  c  t  c  g  q  s                                                   I  q  f  P  v  L  S  I  N
```

$\Delta G^0_{min}$ = -9.36 kcal/mol (left) / -17.19 kcal/mol (right)

**MK11.**

```
                         |-- net 123 bp loss ----------------------------------------------------------------------------------------------------------------------------|
A  ACTACTAGGTCTTCTCTTAGCCTCAGCAGGAAAATCAGCCCAATTTGGACTTCATCCGTGACTTCCATCAGCTAGTGAAGGCCCTACCCCAGTATCAGCACTCCTTCATTCCAGTACAATAGTTATAGCTGGAGTATTTACCCTCATCCGCTTTTATCCATTAATAGAAAACAACCTCAC
   |||||||||||||||||||||| ||||                                                                                                                           ||             | |||||||||||||
R  ACTACTAGGTCTTCTCTTAGCCACAGC---------------------------------------------------------------------------------------------------------------------------CTAAAGCAGCGATTAATAAAACAACCTCAC
         |        |  |||||||||                                                                                                                           ||||||||||||||||||||||||||||
P  CTCTTGAGCATAATCCGAAGCCACAGC---------------------------------------------------------------------------------------------------------------------------CTAAAGCAGCGATTAATAAAACAACCTCGT
Pos                  <-2656541----------------------------------------------------------------------------------------------------------------------------------------------2656574->
R translated      ...S  h  S                                                                                                                             l  k  q  r  l  i  K  Q  P  H...
```

$\Delta G^0_{min}$ = -9.37 kcal/mol (left) / -15.52 kcal/mol (right)

**R1522.**

```
                                                                           |-- net 3 bp inserted --------|
A  ACTACTAGGTCTTCTCTTAGCCTCAGCAGGAAAATCAGCCCAATTTGGACTTCATCCGTGACTTCCATCAGCTAGTGA---AG---GCCCTACCCCAGTATCAGCACTCCTTCATTCCAGTACAATAGTTATAGCTGGAGTATTTACCCTCATCCGCTTTTATCCATTAATAGAAAACA
   |||||||||||||||||||||||||||||||||||||||||||||||||||||||||||||||||||||||| ||||    ||   |||  ||   |||||| ||||||||||||||||||||||||||||||||||||||||||||||||||||||||||||||||||||||||||||
R  ACTACTAGGTCTTCTCTTAGCCTCAGCAGGAAAATCAGCCCAATTTGGACTTCATCCGTGACTTCCATCAGCAAGTGGCACAGATTGCCAAAC---AGTATCTGCACTCCTTCATTCCAGTACAATAGTTATAGCTGGAGTATTTACCCTCATCCGCTTTTATCCATTAATAGAAAACA
        | | | ||    |            ||       |   ||||         |       | |||||||||||||||||||||||||||   ||||||||||||   |   |   |    |          |  |       ||  |        ||| | |       |    |
P  TACTTTCGATTTTTGGATTTATATTAATTCAATTATTAGCATTTTTGTGTCCTTGGCTCATTGGCGATCAGCAAGTGGCACAGATTGCCAAAC---AGTATCTGCACTTTGTTGCTTTTGGTGTACCAGGTTTAGTTTATATGACACTATTTCGCTTCGTTAGCGAAGGTCATAGTTTA
Pos                                                                   <-2700471------------------------2700433->
R translated                                                      ...I  S  k  w  h  r  l  p  n     S  I  c  T...
```

$\Delta G^0_{min}$ = -10.41 kcal/mol (left) / -11.91 kcal/mol (right)

**MK3.**

```
                                                                          |-------| no net gain/loss
A  ACTACTAGGTCTTCTCTTAGCCTCAGCAGGAAAATCAGCCCAATTTGGACTTCATCCGTGACTTCCATCAGCTAGTGAAGGCCCTACCCCAGTATCAGCACTCCTTCATTCCAGTACAATAGTTATAGCTGGAGTATTTACCCTCATCCGCTTTTATCCATTAATAGAAAACAACCTCAC
   ||||||||||||||||||||||||||||||||||||||||||||||||||||||||||||||||||||||| ||   || ||||||||||||||||||||||||||||||||||||||||||||||||||||||||||||||||||||||||||||||||||||||||||||||||||||
R  ACTACTAGGTCTTCTCTTAGCCTCAGCAGGAAAATCAGCCCAATTTGGACTTCATCCGTGACTTCCATCAGTTATCAAATGCCCTACCCCAGTATCAGCACTCCTTCATTCCAGTACAATAGTTATAGCTGGAGTATTTACCCTCATCCGCTTTTATCCATTAATAGAAAACAACCTCAC
   | |   |         | || |         |   | |   ||       |           || ||||||||||||||||||||          |     |  |  |    |  ||     |                   |         |  | | ||| ||    |         |
P  AGTCTCAATCTAAACTTGAGTCATTCTTTAGATGACTGGGAAAAGGATCGTAATCAAAGTTATTACATCAGTTATCAAATGCCCTTTGGATACTACGATTTTGGTCTAAGTTATAACTGGTATCGCTATGATCAGTATGTTGCGGGCTTTAATGCACCGATTCATTATCAGGGTGAAAGC
Pos                                <-2736164-----2736142->
R translated                                                ...T  S  I  S  f  q  m  P  Y...
```

$\Delta G^0_{min}$ = -16.30 kcal/mol

**MK127.**

```
                                 |-- net 42 bp loss --------------------------|
A  ACTACTAGGTCTTCTCTTAGCCTCAGCAGGAAAATCAGCCCAATTTGGACTTCATCCGTGACTTCCATCAGCTAGTGAAGGCCCTACCCCAGTATCAGCACTCCTTCATTCCAGTACAATAGTTATAGCTGGAGTATTTACCCTCATCCGCTTTTATCCATTAATAGAAAACAACCTCAC
   |||||||||||||||||||||||||||||| ||                                           ||||||||||||||||||||||||||||||||||||||||||||||||||||||||||||||||||||||||||||||||||||||||||||||||||||||||
R  ACTACTAGGTCTTCTCTTAGCCTCAGCAGGCAA------------------------------------------CGAAGGCCCTACCCCAGTATCAGCACTCCTTCATTCCAGTACAATAGTTATAGCTGGAGTATTTACCCTCATCCGCTTTTATCCATTAATAGAAAACAACCTCAC
     |    |         |     ||||||||||                                          |||||||||     |     |        |  |            ||    | | |    ||  | ||  | |    |  |                |
P  CTTTACTGCGAGCGGTGTCGATGCAGCAGGCAA------------------------------------------CGAAGGCCCACAAACCACGGCCTGGGCGTTCAAAGTGGTCGGTGCAGCGCCATCAGCGCCATCGATCCAAAGCGTACTAGACGATCAGGGCAGCGTTACTGGCGA
Pos                       <-2804639-------------------------------------------2804621->
R translated         ...L  S  R  q                                            r  R  P...
```

$\Delta G^0_{min}$ = -11.15 kcal/mol (left) / -11.59 kcal/mol (right)

**AC46.**

```
                                                                       |-------------------| net 3 bp loss
A  ACTACTAGGTCTTCTCTTAGCCTCAGCAGGAAAATCAGCCCAATTTGGACTTCATCCGTGACTTCCATCAGCTAGTGAAGGCCCTACCCCAGTATCAGCACTCCTTCATTCCAGTACAATAGTTATAGCTGGAGTATTTACCCTCATCCGCTTTTATCCATTAATAGAAAACAACCTCA
   |||||||||||||||||||||||||||||||||||||||||||||||||||||||||||||||||||| |||| ||| | |     |  ||||||||||||||||||||||||||||||||||||||||||||||||||||||||||||||||||||||||||||||||||||||||||
R  ACTACTAGGTCTTCTCTTAGCCTCAGCAGGAAAATCAGCCCAATTTGGACTTCATCCGTGACTTCCATGAGCTCGTGCATG---TTTGGCAGTATCAGCACTCCTTCATTCCAGTACAATAGTTATAGCTGGAGTATTTACCCTCATCCGCTTTTATCCATTAATAGAAAACAACCTCA
      |    |     |  |    | |       |   |  | | |  | |        |     | ||||||||||||||||   ||||||||||||||||   | | |    | | | | |   |    | ||      |      | |  |||
P  TTAATATTGCGGATTGGTGTGATGATTTTTCGACACAATCGATTGAGAAACAGTCTTGGCTGATTCATGAGCTCGTGCATG---TTTGGCAGTATCAGCAAGGCATAAAGTTAATCCGAAAAGGAATTTTTGAAAGAAAATATCAGTACGTTTTGC
Pos                                                               <-2826900-------------------2826867->
R translated                                                ...T  S  m  S  s  c  m     f  g  S  I  S  T  P...
```

$\Delta G^0_{min}$ = -11.84 kcal/mol (left) / -14.34 kcal/mol (right)

**MK24.**

```
                                                                            |--------------| net 3 bp loss
A  ACTACTAGGTCTTCTCTTAGCCTCAGCAGGAAAATCAGCCCAATTTGGACTTCATCCGTGACTTCCATCAGCTAGTGAAGGCCCTACCCCAGTATCAGCACTCCTTCATTCCAGTACAATAGTTATAGCTGGAGTATTTACCCTCATCCGCTTTTATCCATTAATAGAAAACAACCTCAC
   ||||||||||||||||||||||||||||||||||||||||||||||||||||||||||||||||||||||||| ||| | |     |  |||||||||||||||||||||||||||||||||||||||||||||||||||||||||||||||||||||||||||||||||||||||||||
R  ACTACTAGGTCTTCTCTTAGCCTCAGCAGGAAAATCAGCCCAATTTGGACTTCATCCGTGACTTCCATCAGCTCGTGCATG---TTTGGCAGTATCAGCACTCCTTCATTCCAGTACAATAGTTATAGCTGGAGTATTTACCCTCATCCGCTTTTATCCATTAATAGAAAACAACCTCAC
      |    |     |  |    | |       |   |  |   |  | |        |     | ||| ||||||||||||   ||||||||||||||||     |      |     | |   |    | ||      |           |||    |    | || |
P  TTAATATTGCGGATTGGTGTGATGATTTTTCGACACAATCGATTGAGAAACAGTCTTGGCTGATTCATGAGCTCGTGCATG---TTTGGCAGTATCAGCAAGGCATAAAGTTAATCCGAAAAGGAATTTTTGAAAGAAAATATCAGTACGTTTTGCAGCAGGGAAAGCAGTTTTTGCATT
Pos                                                               <-2826900-------------------2826867->
R translated                                                      ...I  S  s  c  m     f  g  S  I  S  T...
```

$\Delta G^0_{min}$ = -11.84 kcal/mol (left) / -14.34 kcal/mol (right)

```
MK103.                                                      |-- net 18 bp loss ---------------------------|
A  ACTACTAGGTCTTCTCTTAGCCTCAGCAGGAAAATCAGCCCAATTTGGACTTCATCCGTGACTTCCATCAGCTAGTGAAGGCCCTACCCCAGTATCAGCACTCCTTCATTCCAGTACAATAGTTATAGCTGGAGTATTTACCCTCATCCGCTTTTATCCATTAATAGAAAACAACCTCAC
   ||||||||||||||||||||||||||||||||||||||||||||||||||||||||| | |   |||| | | | |     |  ||||    ||  ||||||  ||||||||||||||||||||||||||||||||||||||||||||||||||||||||||||||||||||||||||||
R  ACTACTAGGTCTTCTCTTAGCCTCAGCAGGAAAATCAGCCCAATTTGGACTTCATCC-TCAA--CCATTACC-A-T-----C--TACC----TA--AGCACTAATTCATTCCAGTACAATAGTTATAGCTGGAGTATTTACCCTCATCCGCTTTTATCCATTAATAGAAAACAACCTCAC
   ||       |                  ||| |                  |||||| ||||  |||||||| | |     |  ||||    ||  ||||||||||||||| |      |  |||  || |   ||       |  ||    ||  ||  ||   | ||||  |
P  ACCTCGTCATTAGTAAAGCTAACACGTTGGATACGGTTTATTTTAAATTGATCATCC-TCAA--CCATTACC-A-T-----C--TACC----TA--AGCACTAATTCATTCAATATTTGTCTTTAATGCGGCCTATTTCGTTATTTTCAAGGTTGCTCACTTTTAATAAAATTAGACGTA
Pos                                                   <-2884015------------------------------------------2884056->
R translated                                       ...S  S   s  t    i  t   i          y  l        S  I  n  S  F...
```

$\Delta G^0_{min}$ = -5.72 kcal/mol (left) / -14.21 kcal/mol (right)

```
A71.                                                                    |-------| no net gain/loss
A  ACTACTAGGTCTTCTCTTAGCCTCAGCAGGAAAATCAGCCCAATTTGGACTTCATCCGTGACTTCCATCAGCTAGTGAAGGCCCTACCCCAGTATCAGCACTCCTTCATTCCAGTACAATAGTTATAGCTGGAGTATTTACCCTCATCCGCTTTTATCCATTAATAGAAAACAACCTCAC
   |||||||||||||||||||||||||||||||||||||||||||||||||||||||||||||||||||||  | |||| ||||||||||||||||||||||||||||||||||||||||||||||||||||||||||||||||||||||||||||||||||||||||||||||||||||||||
R  ACTACTAGGTCTTCTCTTAGCCTCAGCAGGAAAATCAGCCCAATTTGGACTTCATCCGTGACTTCCATCTTCAAGTGCAGGCCCTACCCCAGTATCAGCACTCCTTCATTCCAGTACAATAGTTATAGCTGGAGTATTTACCCTCATCCGCTTTTATCCATTAATAGAAAACAACCTCAC
   ||   |||  ||  ||  |    |           | |    |          |    |    | |||||||||||||||||||          |  |      |        |       |     |         | |      ||         |      |         |  |||
P  ACAGTTACCACTGGTTTGAATGGCGTGCATTTGTACCGTAGCAAGCTCTGCCAAAATATCGGGTACATCTTCAAGTGCAGGCCAGTTGAGCTGAATATTTTGACGCGTAGAAACGTGTGCATAACCACGGTCATATTCTTTTGAAATTTCTGCAATTTTGCGTAATTGTTTTGAATTCAT
Pos                                                                <----------------> 2911491-2911508
R translated                                                   ...S  I  f  k  c  R  P...
```

$\Delta G^0_{min}$ = -11.31 kcal/mol

```
MK136.                |-- net 123 bp loss ----------------------------------------------------------------------------------------------------------------------|
A  TAGAAAGATGAAACACTACACTCCCACTACTAGGTCTTCTCTTAGCCTCAGCAGGAAAATCAGCCCAATTTGGACTTCATCCGTGACTTCCATCAGCTAGTGAAGGCCCTACCCCAGTATCAGCACTCCTTCATTCCAGTACAATAGTTATAGCTGGAGTATTTACCCTCATCCGCTTTTA
   |||||||||||||||||||  |                                                                                                                               | |      |||||||||||||||||||||||
R  TAGAAAGATGAAACACTACCTTTTAC---------------------------------------------------------------------------------------------------------------------------AAAAAAACCGTATTTACCCTCATCCGCTTTTA
       |  ||  |||||||||||||||                                                                                                                           |||||||||||||||||| | ||       |
P  ATAGAGGATTGAACACTACCTTTTAC---------------------------------------------------------------------------------------------------------------------------AAAAAAACCGTATTTACCATAATAGTAGATGT
Pos           <-2948975-------------------------------------------------------------------------------------------------------------------------------------------2948943->
R tl      M  K  H  Y  l  l  q                                                                                                                             k  n  r  I  Y  P...
```

$\Delta G^0_{min}$ = -8.33 kcal/mol (left) / -10.16 kcal/mol (right)

```
MK95.                                                            |- net 3 bp inserted-|
A  ACTACTAGGTCTTCTCTTAGCCTCAGCAGGAAAATCAGCCCAATTTGGACTTCATCCGTGAC-TTCC--ATCAGCTAGT-GAAGGCCCTACCCCAGTATCAGCACTCCTTCATTCCAGTACAATAGTTATAGCTGGAGTATTTACCCTCATCCGCTTTTATCCATTAATAGAAAACAACCTCAC
   |||||||||||||||||||||||||||||||||||||||||||||||||||||||||||||| | ||   ||||| |    || ||||||||||||||||||||||||||||||||||||||||||||||||||||||||||||||||||||||||||||||||||||||||||||||||||||
R  ACTACTAGGTCTTCTCTTAGCCTCAGCAGGAAAATCAGCCCAATTTGGACTTCATCCGTGACGTACCTTTTCAGC-ATCCAAATGCCCTACCCCAGTATCAGCACTCCTTCATTCCAGTACAATAGTTATAGCTGGAGTATTTACCCTCATCCGCTTTTATCCATTAATAGAAAACAACCTCAC
   |    |     | |                 || |  |      ||       | ||||||||||||||||||||| |||||||||| ||  |||    |||     |    |      | ||          |     |       |||      |  |     |  ||             |
P  ATGGGTGCAGTTACGAAACTGACACTGGCCTAATTGGGTGTCTATTTCGGGAAAACCGTGACGTACCTTTTCAGC-ATCCAAATGCCATAATCCAAACTCACGGATACCTGGTGAATCAATAAGTGCACCATTTTCTCCAAAAGTAATCAGTCTTGTGGAGGTGGTGGTATGTTGTCCTAGTGC
                                                          <-3049780--------------3049747->
R translated                                          ...S  V  T  y  l  f  s   I  q  m  P...
```

$\Delta G^0_{min}$ = -16.25 kcal/mol

```
MK142.                                                                      |---| no net gain/loss
A  ACTACTAGGTCTTCTCTTAGCCTCAGCAGGAAAATCAGCCCAATTTGGACTTCATCCGTGACTTCCATCAGCTAGTGAAGGCCCTACCCCAGTATCAGCACTCCTTCATTCCAGTACAATAGTTATAGCTGGAGTATTTACCCTCATCCGCTTTTATCCATTAATAGAAAACAACCTCAC
   ||||||||||||||||||||||||||||||||||||||||||||||||||||||||||||||||||||||||| | | ||||||||||||||||||||||||||||||||||||||||||||||||||||||||||||||||||||||||||||||||||||||||||||||||||||||
R  ACTACTAGGTCTTCTCTTAGCCTCAGCAGGAAAATCAGCCCAATTTGGACTTCATCCGTGACTTCCATCAGCTCGGGCAGGCCCTACCCCAGTATCAGCACTCCTTCATTCCAGTACAATAGTTATAGCTGGAGTATTTACCCTCATCCGCTTTTATCCATTAATAGAAAACAACCTCAC
   | ||  | ||     |  |     |     ||  |          | ||   |             |  ||||||||||||| |    |||  |     |     |   |  |  |            | | |  |    |     |     |    |
P  AATAAAAAGTACAGGCACACTAATATTTCTAAGTTGCTGAATGGGTCGAAAACGAATCAATTGAAAAAAAGCTCGGGCAGGCACCTGACCACTCCAGTAATAATCTGGATTGACGATTGATAAATATCCTAAACGACTATCAGAAGTAGGTAAAAAGCAAAGTTGCTGTTCTGCAACTAC
Pos                                                               3064595<------------->3064608
R translated                                                         ...S  s  g  R  P...
```

$\Delta G^0_{min}$ = -12.72 kcal/mol

```
MK12.                                                                       |-- net 18 bp loss ----------------|
A  ACTACTAGGTCTTCTCTTAGCCTCAGCAGGAAAATCAGCCCAATTTGGACTTCATCCGTGACTTCCATCAGCTAGTGAAGGCCCTACCCCAGTATCAGCACTCCTTCATTCCAGTACAATAGTTATAGCTGGAGTATTTACCCTCATCCGCTTTTATCCATTAATAGAAAACAACCTCAC
   |||||||||||||||||||||||||||||||||||||||||||||||||||||||||||||||||||||||||  |     || | | ||    |||  |   | ||| |||||||||||||||||||||||||||||||||||||||||||||||||||||||||||||||||||||||
R  ACTACTAGGTCTTCTCTTAGCCTCAGCAGGAAAATCAGCCCAATTTGGACTTCATCCGTGACTTCCATCAGCT--T-----CCAT-CTCC----TCA--A---C-TCAGTCCAGTACAATAGTTATAGCTGGAGTATTTACCCTCATCCGCTTTTATCCATTAATAGAAAACAACCTCAC
   |    |   | ||    |     || ||  ||    |      |  |       |     |    ||||||||  |     |||| ||||    |||  |   | |||||||||  ||        |  |     |  |   |               |    |             |
P  ACGTGTTTATTTTGGAGTCATTACAACAAAAATGGAAAAAACCTCAGATAAAAGTACTTAATATTCATCAGCT--T-----CCAT-CTCC----TCA--A---C-TCAGTCCAGCTCAGGTTCAGAAAATTAGTCAAATCAGCCGTAACACAGGGTATCCTGTATCATTTATTCAACCTA
Pos                                                                <-3101742--------------------------------3101772->
R translated                                                   ...S  I  S  f         h   l  l      n        S  v  Q...
```

$\Delta G^0_{min}$ = -9.49 kcal/mol (left) / -8.03 kcal/mol (right)

```
MK36.                           |-- net 27 bp inserted --------------------------------------------------------------------------------|
A  ACTACTAGGTCTTCTCTTAGCCTCAGCAGGAAAATCAGCCCAATTTGGACTTCATCCGTGACTTCC---------------------------ATCAGCTAGTGAAGGCCCTACCCCAGTATCAGCACTCCTTCATTCCAGTACAATAGTTATAGCTGGAGTATTTACCCTCATCCGCTT
   ||||||||||||||||||||||||||||| ||||||  |          |   |       |                                |||        || ||   |   |   |    ||     |||||||||||||||||||||||||||||||||||||||||||||||
R  ACTACTAGGTCTTCTCTTAGCCTCAGCAGAAAAATCGCCAATCCCACAGCCAGACTAAATGCAGTGGTTAAATCTGGAATCCACATGGCACAGCTCACACCAATCAGACCTGCCAATAAATTTCCAACGATCACATTCCAGTACAATAGTTATAGCTGGAGTATTTACCCTCATCCGCTT
         |     ||| |   | ||| |||||||||||||||||||||||||||||||||||||||||||||||||||||||||||||||||||||||||||||||||||||||||||||||||             |      |       |  |
P  GGATGCAATGAATCTGTACTCATCATCAGAAAAATCGCCAATCCCACAGCCAGACTAAATGCAGTGGTTAAATCTGGAATCCACATGGCACAGCTCACACCAATCAGACCTGCCAATAAATTTCCAACGATCACATTCCAGGGTTGAGCCAGGGGACTTGATGGCACAGCAAAAGTAAA
Pos                    <-3179683-------------------------------------------------------------------------------------------------------3179803->
R translated      ...S  L  S  R  K  I  a  n  p  t  a  r  l  n  a  v  v  k  s  g  i  h  m  a  q  l  t  p  i  r  p  a  n  k  f  p  t  i  t  F  Q...
```

$\Delta G^0_{min}$ = -12.37 kcal/mol (left) / -9.05 kcal/mol (right)

```
R19.                                                                 |-- net 45 bp loss ------------------------------------------|
A  TTTGAGAATAGAAAGATGAAACACTACACTCCCACTACTAGGTCTTCTCTTAGCCTCAGCAGGAAAATCAGCCCAATTTGGACTTCATCCGTGACTTCCATCAGCTAGTGAAGGCCCTACCCCAGTATCAGCACTCCTTCATTCCAGTACAATAGTTATAGCTGGAGTATTTACCCTCAT
   ||||||||||||||||||||||||||||||||||||||||||||||||||||||||||||||||||              || | |||   | | ||                            || ||||||||||||||||||||||||||||||||||||||||||||||||||||
R  TTTGAGAATAGAAAGATGAAACACTACACTCCCACTACTAGGTCTTCTCTTAGCCTCAGCAGGAAA--------------GATTGCAT---TAAATT----------------------------TACCAGCACTCCTTCATTCCAGTACAATAGTTATAGCTGGAGTATTTACCCTCAT
    |   | | |  |     | |   | |||    ||  |  |   |  |     |  ||||||||||              ||||||||   ||||||                            ||||||||| | |||| |  |            |     ||     | |  |
P  ATAACGTACACCATTGACAGAAGATTCACCTTGACCGCATGCAGTCGTTGGTACGCCAGCAGGAAA--------------GATTGCAT---TAAATT----------------------------TACCAGCAC-C-TTCAGTGGATAGACTATACGCTGCACCGACAGAGTTCATTGGA
Pos                                                        <-3216659-------------------------------------------------------------------3216622->
R translated                                          ...L  S  R  K                d  c  i     k  f                              t  S  T  P  S...
```

$\Delta G^0_{min}$ = -13.79 kcal/mol (left) / -11.82 kcal/mol (right)

```
MK102.                                                     |-- net 6 bp loss ------|
A  ACTACTAGGTCTTCTCTTAGCCTCAGCAGGAAAATCAGCCCAATTTGGACTTCATCCGTGACTTCCATCAGC-TAGTGAAGGCCCTACCCCAGTATCAGCACTCCTTCATTCCAGTACAATAGTTATAGCTGGAGTATTTACCCTCATCCGCTTTTATCCATTAATAGAAAACAACCTCAC
   ||||||||||||||||||||||||||||||||||||||||||||||||||||||||    | | || |  || | ||| | ||||||||||||||||||||||||||||||||||||||||||||||||||||||||||||||||||||||||||||||||||||||||||||||||||||
R  ACTACTAGGTCTTCTCTTAGCCTCAGCAGGAAAATCAGCCCAATTTGGACTTCATC----AATGCC-T--GCTTCGTGCATGCCCTACCCCAGTATCAGCACTCCTTCATTCCAGTACAATAGTTATAGCTGGAGTATTTACCCTCATCCGCTTTTATCCATTAATAGAAAACAACCTCAC
    | ||   |    |   ||    |      || |    |  |  |   ||||||||    |||||| |  |||||||||||||||| |           || | |  |    |     ||||    |   |    | |                |  ||   | |     |      | |
P  CCAACACAGATGGCCAATACAGGCTTGTTAAATACAGCCTGACGTACCACTTCATC----AATGCC-T--GCTTCGTGCATGCCCTGCATACAGTCGCGCATTGCACCCACACCTGGAAATACGATTTTATCAGCTTTAGCAATCAGCTTTGGATCATTGGTGACATCTACGGTTGCCCCG
Pos                                                <-3310290----------------------3310258->
R translated                                    ...T  S  S      m  p     a  s  c  m  P  Y...
```

$\Delta G^0_{min}$ = -13.82 kcal/mol

```
R5.                                                                     |-------------------| net 6 bp inserted
A  ACTACTAGGTCTTCTCTTAGCCTCAGCAGGAAAATCAGCCCAATTTGGACTTCATCCGTGACTTCCATCAGCTAGTGAAGGCCC------TACCCCAGTATCAGCACTCCTTCATTCCAGTACAATAGTTATAGCTGGAGTATTTACCCTCATCCGCTTTTATCCATTAATAGAAAACAAC
   ||||||||||||||||||||||||||||||||||||||||||||||||||||||||||||||||||||| || |   |||  ||      |||||||||||||||||||||||||||||||||||||||||||||||||||||||||||||||||||||||||||||||||||||||||||
R  ACTACTAGGTCTTCTCTTAGCCTCAGCAGGAAAATCAGCCCAATTTGGACTTCATCCGTGACTTCCATCTGCCACCAAAGAACCCATCATTACCCCAGTATCAGCACTCCTTCATTCCAGTACAATAGTTATAGCTGGAGTATTTACCCTCATCCGCTTTTATCCATTAATAGAAAACAAC
   |    |     |          ||      ||  |  |      | | |    |      | |   ||||||||||||||||||||||||||||||||||||      |    |      |  |      |          ||  ||  |||  |   |      | | | |  || || |
P  ATCTTTCCCAATCACTGCCATATCGTGTTTAAGCTTCGTATTCATGGCATCAAAAGAAATATTCTGATCTGCCACCAAAGAACCCATCATTACCCCAGTATCCTGCTGCGGAAAAAATCCTTTAGGTACCACGATATATAAATAAACATTCAGTCCAATGGTAAAAATGAAAATAATCAGC
Pos                                                                  <-3333089------------------3333124->
R translated                                                      ...I  c  h  q  R  t  h  h  Y  P  S  I...
```

$\Delta G^0_{min}$ = -4.56 kcal/mol (left) / -16.32 kcal/mol (right)

```
K2.                               |-- net 36 bp loss ------------------------------------------|
A  ACTACTAGGTCTTCTCTTAGCCTCAGCAGGAAAATCAGCCCAATTTGGACTTCATCCGTGACTTCCATCAGCTAGTGAAGGCCCTACCCCAGTATCAGCACTCCTTCATTCCAGTACAATAGTTATAGCTGGAGTATTTACCCTCATCCGCTTTTATCCATTAATAGAAAACAACCTCAC
   |||||||||||||||||||||||||||||| |                  ||  ||| ||  | ||| || ||                || ||||||||||||||||||||||||||||||||||||||||||||||||||||||||||||||||||||||||||||||||||||||||
R  ACTACTAGGTCTTCTCTTAGCCTCAGCAGGCA------------------TTACTCCATG--TACCA-CACCTG---------------CAATATCAGCACTCCTTCATTCCAGTACAATAGTTATAGCTGGAGTATTTACCCTCATCCGCTTTTATCCATTAATAGAAAACAACCTCAC
   || |  || |  |       | ||||||||||                  ||||||||||  ||||| ||||||               ||||||||||||| |   |||  | ||   ||    |     |    |  |                   |    ||  |||        |
P  ACGATAAGCTGCTGCTGCGCCTTCAGCAGGCA------------------TTACTCCATG--TACCA-CACCTG---------------CAATATCAGCACTACACAATTGAACTAATCTACGCGTTAAAAGGCACCCATAGAGTGAGTTACGATAATAAATTTATGAAAGCGTGCACC
Pos                    <-3340507------------------------------------------------------------------3340554->
R translated         ...L  S  R  h                    y  s  m    y  h  t  c                  n  I  S  T  P...
```

$\Delta G^0_{min}$ = -11.83 kcal/mol (left) / -14.33 kcal/mol (right)

```
K6.                                 |-- net 129 bp loss ------------------------------------------------------------------------------------------------------------|
A  CAGCCCAATTTGGACTTCATCCGTGACTTCCATCAGCTAGTGAAGGCCCTACCCCAGTATCAGCACTCCTTCATTCCAGTACAATAGTTATAGCTGGAGTATTTACCCTCATCCGCTTTTATCCATTAATAGAAAACAACCTCACTATTCAAACTTCAACACTTCTGGTTCTGGCTCTA
   |||||||||||||||||||||||||||||||||                                                                                                                                 |||||||||||||||||
R  CAGCCCAATTTGGACTTCATCCGTGACTTCCAT---------------------------------------------------------------------------------------------------------------------------------TTCTGGTTCTGGCTCTA
           |    | ||| |   ||||||||||                                                                                                                                 |||||     |  |||
P  TTCATTCTTCAAAAATTCCTGTTTGACTTCCAT---------------------------------------------------------------------------------------------------------------------------------TTCTGCAATGGTATCTC
Pos                       <-3346647------------------------------------------------------------------------------------------------------------------------------3346661->
R translated          ...V  T  S  I                                                                                                                                   S  G...
```

$\Delta G^0_{min}$ = -12.64 kcal/mol (left) / -3.61 kcal/mol (right)

**MK43.**

```
                                                                               |-- net 6 bp loss ------|
A  ACTACTAGGTCTTCTCTTAGCCTCAGCAGGAAAATCAGCCCAATTTGGACTTCATCCGTGACTTCCATCAGCTAGTGAAGGCCCTACCCCAGTATCAGCACTCCTTCATTCCAGTACAATAGTTATAGCTGGAGTATTTACCCTCATCCGCTTTTATCCATTAATAGAAAACAACCTCAC
R  ACTACTAGGTCTTCTCTTAGCCTCAGCAGGAAAATCAGCCCAATTTGGACTTCATCCGTGACTTCCATCAAC-AGCG-CGG--C-A-CCCAGTACCAGCACTCCTTCATTCCAGTACAATAGTTATAGCTGGAGTATTTACCCTCATCCGCTTTTATCCATTAATAGAAAACAACCTCAC
P  GTCGATTCAGACAACAACACACCCGCCAGAAATGCGCCCATCGCCATCGACAAACCACTGACCTGCATCAAC-AGCG-CGG--C-A-CCCAGTACCACCAGCAAGGCTGCTGCGGTCATCACTTCACGCGCTTTGGATGCGGCCAGTAATCTAAACAGTGGATTAAGCAACCAGAAACCT
Pos                                                                    <-3360211------------------------3360246->
R translated                                                       ...V  T  S  I  n   s  a   a     P  S  t  S  T...
```

$\Delta G^0_{min}$ = -25.91 kcal/mol

**MK25.**

```
                                  |-- net 3 bp loss ---------------------------------------------------|
A  ACTACTAGGTCTTCTCTTAGCCTCAGCAG-GAAAATCAGCCCAATTTGGACTTCATCCGTGACTTCCATCAGCTAGTGAAGGCCCTACCC-CAGT--ATCAGCACTCCTTCATTCCAGTACAATAGTTATAGCTGGAGTATTTACCCTCATCCGCTTTTATCCATTAATAGAAAACAACC
R  ACTACTAGGTCTTCTCTTAGCCTCAGCCGTCAAAATTTGCGCAA---GGTTTTC-TTTGTG--TTCATTGATCAGGTGAA-ACCATGCCCAGAGTACATCAGCACTCCTTCATTCCAGTACAATAGTTATAGCTGGAGTATTTACCCTCATCCGCTTTTATCCATTAATAAAAAACAACC
P  TTCTGCCAATGGCTTGCCTTGCTCAGCCGTCAAAATTTGCGCAA---GGTTTTC-TTTGTG--TTCATTGATCAGGTGAA-ACCATGCCCAGAGTACATCAGCACGTTGTAATGCGGTCTGGGTTTTCCATTTGGCCTGTGCTTGAGCCGAACGCTGAATTGCAGCTTCAACACCTGAAC
Pos                          <-3366020-----------------------------------------------------------------3366096->
R translated             ...L  S  r  q  N  l  r  k     v  f   f  v   f  i  d  q  v  k   p  c  p  e  y  I  S  T...
```

$\Delta G^0_{min}$ = -11.57 kcal/mol (left) / -21.37 kcal/mol (right)

**U96.**

```
                                                                                         |----| no net gain/loss
A  ACTACTAGGTCTTCTCTTAGCCTCAGCAGGAAAATCAGCCCAATTTGGACTTCATCCGTGACTTCCATCAGCTAGTGAAGGCCCTACCCCAGTATCAGCACTCCTTCATTCCAGTACAATAGTTATAGCTGGAGTATTTACCCTCATCCGCTTTTATCCATTAATAGAAAACAACCTCAC
R  ACTACTAGGTCTTCTCTTAGCCTCAGCAGGAAAATCAGCCCAATTTGGACTTCATCCGTGACTTCCATCAGCAATTGCAGGCCCTACCCCAGTATCAGCACTCCTTCATTCCAGTACAATAGTTATAGCTGGAGTATTTACCCTCATCCGCTTTTATCCATTAATAGAAAACAACCTCAC
P  AGCATGACTTGAAATACCATTATCGACAGTGATCAACAGCTCTGGTTGACATTGTTGATGCGCCAGATCAGCAATTGCAGGGGTTAAGCCATAACCATACTTAAAGCGATCAGGCACAATGTAATCAACTCTTGCGCCCATTTCTTTTAAAGCTAGACACATCAAAGCCGTACTGGTTGC
Pos                                                                    <-------------> 3425419-3425433
R translated                                                       ...I  S  n  c  R...
```

$\Delta G^0_{min}$ = -11.38 kcal/mol

**MK29.**

```
                                        |-- net 18 bp loss ------------------------------|
A  ACTACTAGGTCTTCTCTTAGCCTCAGCAGGAAAATCAGCCCAATTTGGACTTCATCCGTGACTTCCATCAGCTAGTGAAGGCCCTACCCCAGTATCAGCACTCCTTCATTCCAGTACAATAGTTATAGCTGGAGTATTTACCCTCATCCGCTTTTATCCATTAATAGAAAACAACCTCAC
R  ACTACTAGGTCTTCTCTTAGCCTCAGCAGGAAAATCATCCAAATTTACCGATGCTGCA------------------TGGACCGATTACCCAGTATCAGCACTCCTTCATTCCAGTACAATAGTTATAGCTGGAGTATTTACCCTCATCCGCTTTTATCCATTAATAGAAAACAACCTCAC
P  GCGGCGATGTATATGCAGAAGCTGAATTAAAAAATCATCCAAATTTACCGATGCTGCA------------------TGGACCGATTACCCAGTCTAAAGCGATGATGCGTCGTTACAATGAAATTAATCAATTATTCCAGCCTGTCAATTTACGTTTAACTGAGTTATATCTAACAGAAC
Pos                               <-3441979------------------------------------------------3442022->
R translated                  ...K  I  i  q  i  y  r  c  c  m                  d  r  l  P  S...
```

$\Delta G^0_{min}$ = -14.71 kcal/mol (left) / -7.01 kcal/mol (right)

**MK14.**

```
                                                                               |-- no net gain/loss ----|
A  ACTACTAGGTCTTCTCTTAGCCTCAGCAGGAAAATCAGCCCAATTTGGACTTCATCCGTGACTTCCATCAGCTAGTGAA-G-GCC-CTACCCCAGTATCAGCACTCCTTCATTCCAGTACAATAGTTATAGCTGGAGTATTTACCCTCATCCGCTTTTATCCATTAATAGAAAACAACCTCAC
R  ACTACTAGGTCTTCTCTTAGCCTCAGCAGGAAAATCAGCCCAATTTGGACTTCATCCGTGACTTCCATCA-CCACAAACTGCGCCACCACCCCA--ATCAGCACTCCTTCATTCCAGTACAATAGTTATAGCTGGAGTATTTACCCTCATCCGCTTTTATCCATTAATAGAAAACAACCTCAC
P  ATCACACCAATCGCGATCTCTGAAGGCAACTGAAATAACTGACACAGTAAAAAGGCCAACCCTGGCATCA-CCACAAACTGCGCCACCACCCCA--ATCAGCACAGCTTTGGGATAGCGCAATATTCCCTGAAAATCTTGTAAGGACATGGTCATGCCCATCCCAAGCATAATGATCCCCAGC
Pos                                                                   <-3502921-----------------------3502959->
R translated                                                      ...S  I  t   t  n  c  a  t  t  p    S  I  T...
```

$\Delta G^0_{min}$ = -3.44 kcal/mol (left) / -21.47 kcal/mol (right)

**MK96.**

```
                                         |-- net 24 bp inserted ----------------------------------------------------------|
A  ACTACTAGGTCTTCTCTTAGCCTCAGCAGGAAAATCAG-CCCAATTTGGACTTCATCCGTGACTT------------------------CCATCAGCTAGTGAAGGCCCTACCCCAGTATCAGCACTCCTTCATTCCAGTACAATAGTTATAGCTGGAGTATTTACCCTCATCCGCTTTTA
R  ACTACTAGGTCTTCTCTTAGCCTCAGCAGGAAAATCAGTCCCA-TTGCCACTGTAGGTAACCCTAGACCTAATGTACCTTTAACCAGCCCTGCAAAAATAAAAACCCCCATGACCAGTAGCAGCACTCCTTCATTCCAGTACAATAGTTATAGCTGGAGTATTTACCCTCATCCGCTTTTA
P  GCTGCTGCCTGAAAAGGTGAAATCACCATACTCATCAGTCCCA-TTGCCACTGTAGGTAACCCTAGACCTAATGTACCTTTAACCAGCCCTGCAAAAATAAAAACCCCCATGACCAGTAGCAGCATGTCCATTTGTTTTCTCTCCAGATTCAGTCTTTATGCTAACCAAATTAAGATCAGG
Pos                                 <-3544557---------------------------------------------------------------3544647->
R translated                     ...I  S  p  i   a  t  v  g  n  p  r  p  n  v  p  l  t  s  p  a  k  i  k  t  p  m  t  S  s  S  T...
```

$\Delta G^0_{min}$ = -9.84 kcal/mol (left) / -14.21 kcal/mol (right)

-----

SPDIR mutations formed with donor DNA from rRNA operons:

**MK145.**

```
                                                                                                                                                                |- net 5 bp loss -|
A   TAGCCTCAGCAGGAAAATCAGCCCAATTTGGACTTCATCCGTGACTTCCATCAGCTAGTGAAGGCCCTACCCCAGTATCAGCACTCCTTCATTCCAGTACAATAGTTATAGCTGGAGTATTTACCCTCATCCGCTTTTATCCATTAATAGAAAACAACCTCACTATTCAAACTTCAACAC
    |||||||||||||||||||||||||||||||||||||||||||||||||||||||||||||||||||||||||||||||||||||||||||||||||||||||||||||||||||||||||||||||||||||||||||||| |||  |||  |  |||  ||||||||||||||||
R   TAGCCTCAGCAGGAAAATCAGCCCAATTTGGACTTCATCCGTGACTTCCATCAGCTAGTGAAGGCCCTACCCCAGTATCAGCACTCCTTCATTCCAGTACAATAGTTATAGCTGGAGTATTTACCCTCATCCGCTTTTATCCATTCATA-CAAATGA--TCA--ATTCAAACTTCAACAC
     |       |   |||         |         | |   |   ||  ||     ||   ||          ||       | |       |   |    |       |   |          |  |         || ||||||| |||||||  |||  |||||||||| |
P   AATTTATTCCTCTAAAGAGCCGTTCAAGACTAGGACGTTGATAGGTTGGATGTGGAAGCATAGTGATATGTGAAGCTGACCAATACTAATTGCTCGTGAGGCTTGACTATACAACACCCAAACAGTTGTTGTATCAAGATAAATTCATA-CAAATGA--TCA--ATTCAAACTTGATTTA
Pos                         23461..23487; 218246..218272; 651174..651200; 1665745..1665771; 2942228..2942202; 3071325..3071299; 3559737..3559711 <------------------------------>
R translated                                                                                                                                                        m  i    n    S  N  F...
```

$\Delta G^0_{min}$ = -9.40 kcal/mol

Note: Ectopic DNA patch originating from one of the seven rRNA operon in *A. baylyi*.

-----

**K143.**

```
                                                                                 |------| no net gain/loss
A   ACTACTAGGTCTTCTCTTAGCCTCAGCAGGAAAATCAGCCCAATTTGGACTTCATCCGTGACTTCCATCAGCTAGTGAAGGCCCTACCCCAGTATCAGCACTCCTTCATTCCAGTACAATAGTTATAGCTGGAGTATTTACCCTCATCCGCTTTTATCCATTAATAGAAAACAACCTCAC
    ||||||||||||||||||||||||||||||||||||||||||||||||||||||||||||||||||||||||||    ||| ||||||||||||||||||||||||||||||||||||||||||||||||||||||||||||||||||||||||||||||||||||||||||||||||||||||
R   ACTACTAGGTCTTCTCTTAGCCTCAGCAGGAAAATCAGCCCAATTTGGACTTCATCCGTGACTTCCATCAGCTACCTTAGGACCTACCCCAGTATCAGCACTCCTTCATTCCAGTACAATAGTTATAGCTGGAGTATTTACCCTCATCCGCTTTTATCCATTAATAGAAAACAACCTCAC
           ||       |    |||            ||  |   ||  |  |||  | |  |||  |       |  ||||||||||||||        ||    | |   ||  |     | | ||    |      |      || || ||||   |     | |    |       |
P   CTCTGCTGGAGACAGCGCCGCCATCATTATGCCATTCGTGCAGGTCGGAACTTACCCGACAAGGAATTTCGCTACCTTAGGACCGTTATAGTTACGGCCGCCGTTTACTGGGGCTTCGATCAAGAGCTTCGCTTACGCTAACCCCATCAATTAACCTTCCAGCACCGGGCAGGCATCACA
POS                                                                       <------------->  22459..22446; 217244..217231; 650171..650158; 1664743..1664730; 2943230.. 2943243;
                                                                                           3072327..3072340; 3560739..3560752 (A. baylyi)
        |   |              |  |     |  |  |    | |||  | | |||  |      |  ||||||||||||||        ||    | |   ||  |     | ||||    |      |      ||  | |  |          | |    |       |       |
BS  CTCTCGTTGAGACAGTGCCCAGATCGTTGCGCCTTTCGTGCGGGTCGGAACTTACCCGACAAGGAATTTCGCTACCTTAGGACCGTTATAGTTACGGCCGCCGTTTACTGGGGCTTCAATTCGCACCTTCGCTTACGCTAAGCGCTCCTCTTAACCTTCCAGCACCGGGCAGGCGTCAGC
POS                                                                       <------------->  136571..13658; 34139..34126; 94216..94203; 100072..100059; 164571..164558; 170177..170164;
                                                                                           175176..175163; 639117..639104; 950380..950367; 3174956..3174969 (B. subtilis)
R translated                                                           ...S  y  l  R  t...
```

$\Delta G^0_{min}$ = -4.46 kcal/mol

Note: Found in a transformation experiment with *B. subtilis* DNA. Ectopic DNA patch originating from an rRNA operon either from *A. baylyi* or *B. subtilis* (17 possible sources).

-------

SPDIR mutations formed with donor DNA from *Bacillus subtilis* 168 [NC_000964]:

**B127.**

```
                                                                              |---| no net gain/loss
A   ACTACTAGGTCTTCTCTTAGCCTCAGCAGGAAAATCAGCCCAATTTGGACTTCATCCGTGACTTCCATCAGCTAGTGAAGGCCCTACCCCAGTATCAGCACTCCTTCATTCCAGTACAATAGTTATAGCTGGAGTATTTACCCTCATCCGCTTTTATCCATTAATAGAAAACAACCTCAC
    ||||||||||||||||||||||||||||||||||||||||||||||||||||||||||||||||||||||||| ||  |||||||||||||||||||||||||||||||||||||||||||||||||||||||||||||||||||||||||||||||||||||||||||||||||||||||
R   ACTACTAGGTCTTCTCTTAGCCTCAGCAGGAAAATCAGCCCAATTTGGACTTCATCCGTGACTTCCATCAGGTATAGAAGGCCCTACCCCAGTATCAGCACTCCTTCATTCCAGTACAATAGTTATAGCTGGAGTATTTACCCTCATCCGCTTTTATCCATTAATAGAAAACAACCTCAC
    |    ||| |  |    | |       |         |||   | ||      | |     || | |  ||||||||||||||||| ||       |      |    |             | | |    |     |       |     |   |||      |   |   | |
BS  ATATGTAGTTTGTGGATCAAAAGAGGGTATGTTATCTATCAAACAAACTCGTTCAGAGTTAGTCGAATCAGGTATAGAAGGCCTTAAGGATATTATCTTAAAAATAACCGGAGAAAAAGTGAAAAGTTTTCATACTGATTTAAGCTCCCGGACAGGTGAACGAGTGATGGTATTTAAATT
Pos                                                                    <-210793----210812->
R translated                                                        ...I  r  y  r  R  P  Y...
```

$\Delta G^0_{min}$ = -11.84 kcal/mol

**B45.**

```
                                          |-- net 111 bp loss -------------------------------------------------------------------------------------------|
A   AAAATCAGCCCAATTTGGACTTCATCCGTGACTTCCATCAGCTAGTGAAGGCCCTACCCCAGTATCAGCACTCCTTCATTCCAGTACAATAGTTATAGCTGGAGTATTTACCCTCATCCGCTTTTATCCATTAATAGAAAACAACCTCACTATTCAAACTTCAACACTTCTGGTTCTGGC
    |||||||||||||||||||||||||||||| |||||                                                                                                               ||||||||||||||||||||
R   AAAATCAGCCCAATTTGGACTTCATCCGTGACCTCCATGTTCATCCAT---------------------------------------------------------------------------------------------------TTCAACACTTCTGGTTCTGGC
     |   || |    ||| |   || |  ||||||||||||||||||||                                                                                                               ||||||||||||
BS  GATTCCATCATTTTTTTGTTATCTTTTGTGACCTCCATGTTCATCCAT---------------------------------------------------------------------------------------------------TTCAACACTTCTTCACGCTCG
Pos                         <-541057------------------------------------------------------------------------------------------------------------------------541025->
R translated              ...V  T  S  m  f  i  h                                                                                                                F  N  T  S...
```

$\Delta G^0_{min}$ = -11.62 kcal/mol (left) / -15.59 kcal/mol (right)

**AB8.**

```
                                                                                 |-| no net gain/loss
A   ACTACTAGGTCTTCTCTTAGCCTCAGCAGGAAAATCAGCCCAATTTGGACTTCATCCGTGACTTCCATCAGCTAGTGAAGGCCCTACCCCAGTATCAGCACTCCTTCATTCCAGTACAATAGTTATAGCTGGAGTATTTACCCTCATCCGCTTTTATCCATTAATAGAAAACAACCTCAC
    |||||||||||||||||||||||||||||||||||||||||||||||||||||||||||||||||||||||||||| | ||||||||||||||||||||||||||||||||||||||||||||||||||||||||||||||||||||||||||||||||||||||||||||||||||||||||
R   ACTACTAGGTCTTCTCTTAGCCTCAGCAGGAAAATCAGCCCAATTTGGACTTCATCCGTGACTTCCATCAGCTATTCAAGGCCCTACCCCAGTATCAGCACTCCTTCATTCCAGTACAATAGTTATAGCTGGAGTATTTACCCTCATCCGCTTTTATCCATTAATAGAAAACAACCTCAC
          ||   || | | |          |            | ||   ||||     |    | ||||||||||| ||     ||     |     ||  | |  |    | | |   |    |  |     | ||               | |   | |   |
BS  GAGCTGAGTCTTTATTTAAATACACATCCTCATGATGAAGATGCGTTGAAGCAATTCAATCAATATTCCGGCTATTCAAGGCACTTAAAAAGACAGTTCGAATCCTCTTACGGACCGCTTCTGCAGTTCGGCAACAGCCCCGCGGGCAAGGATTGGGATTGGGGAAAGGGCCATGGCCG
Pos                                                                      <--------------> 756279-756295
R translated                                                        ...I  S  y  s  R  P...
```

$\Delta G^0_{min}$ = -14.70 kcal/mol (BS)

**AB15.**

```
                                                                          |-----| no net gain/loss
A  TCTACTAGGTCTTCTCTTAGCCTCAGCAGGAAAATCAGCCCAATTTGGACTTCATCCGTGACTTCCATCAGCTAGTGAAGGCCCTACCCCAGTATCAGCACTCCTTCATTCCAGTACAATAGTTATAGCTGGAGTATTTACCCTCATCCGCTTTTATCCATTAATAGAAAACAACCTCAC
   ||||||||||||||||||||||||||||||||||||||||||||||||||||||||||||||||||||||| |  | |||||||||||||||||||||||||||||||||||||||||||||||||||||||||||||||||||||||||||||||||||||||||||||||||||||||||||
R  TCTACTAGGTCTTCTCTTAGCCTCAGCAGGAAAATCAGCCCAATTTGGACTTCATCCGTGACTTCCATCAGATGTTCAAGGCCCTACCCCAGTATCAGCACTCCTTCATTCCAGTACAATAGTTATAGCTGGAGTATTTACCCTCATCCGCTTTTATCCATTAATAGAAAACAACCTCAC
     ||    | |       |        | |       | |     |     |   |     |        | |||||||||||||||||||  | ||          |
BS AATAAAGAGACACGCGCTCCATATAGGATCCGGCTGAAAATATCACGTCCAAATTTATCCGTTCCCATCAGATGTTCAAGGCCCGGCGCCTTATATTTATCAAGCAGGCTTTGCTGCTCTTGGGGATAAGGTGCAATTAAAGGGGCGAAAACAGCAGACATAATAATGATAAAAAGAATA
Pos                                                             <-1216456----1216437->
R translated                                               ...T  S  I  r  c  s  R  P...
```

$\Delta G^0_{min}$ = -23.99 kcal/mol

**AD60.**

```
                                                                  |-- net 45 bp loss ---------------------------------------------------|
A  TTTGAGAATAGAAAGATGAAACACTACACTCCCACTACTAGGTCTTCTCTTAGCCTCAGCAGGAAAATCAGCCCAATTTGGACTTCATCCGTGACTTCCATCAGCTAGTGAAGGCCCTACCCCAGTATCAGCACTCCTTCATTCCAGTACAATAGTTATAGCTGGAGTATTTACCCTCATC
   ||||||||||||||||||||||||||||||||||||||||||||||||||||||||||||||| ||| ||||   | |                                                     ||||||||||||||||||||||||||||||||||||||||||||||||
R  TTTGAGAATAGAAAGATGAAACACTACACTCCCACTACTAGGTCTTCTCTTAGCCTCAGCAGGCAAA-CAGCATCAATGACCACGTTTG--------------------------------------------CTCCTTCATTCCAGTACAATAGTTATAGCTGGAGTATTTACCCTCATC
            | |               ||||  || | | |      | |       |  | ||||||||| ||||||||||||||||||||                                            ||||||||||||     | ||       | | |              |  |
BS        CCACATGTCCCGCCGGTGACACAGCCTCAATTGAAAGTGCAAAAGTCTCCGGCAGGCAAA-CAGCATCAATGACCACGTTTG--------------------------------------------CTCCTTCATTCCAGTCCATTCAAGGACCCGCTCGGCAACATGTTCTG
Pos                                                        <-1305161-----------------------------------------------------------------------------1305118->
R translated                                      ...S  L  S  R  q  t  a  s  m  t  t  f  a                                              P  S  F  Q  Y...
```

$\Delta G^0_{min}$ = -15.51 kcal/mol (left) / -16.30 kcal/mol (right)

**B68.**

```
                                                                            |-| no net gain/loss
A  ACTACTAGGTCTTCTCTTAGCCTCAGCAGGAAAATCAGCCCAATTTGGACTTCATCCGTGACTTCCATCAGCTAGTGAAGGCCCTACCCCAGTATCAGCACTCCTTCATTCCAGTACAATAGTTATAGCTGGAGTATTTACCCTCATCCGCTTTTATCCATTAATAGAAAACAACCTCAC
   |||||||||||||||||||||||||||||||||||||||||||||||||||||||||||||||||||||||||   ||||||||||||||||||||||||||||||||||||||||||||||||||||||||||||||||||||||||||||||||||||||||||||||||||||||||||||||||
R  ACTACTAGGTCTTCTCTTAGCCTCAGCAGGAAAATCAGCCCAATTTGGACTTCATCCGTGACTTCCATCAGCTGTAGAAGGCCCTACCCCAGTATCAGCACTCCTTCATTCCAGTACAATAGTTATAGCTGGAGTATTTACCCTCATCCGCTTTTATCCATTAATAGAAAACAACCTCAC
   |               ||      |  ||  ||| |  |  | |||    |      |||    ||||| |||||||||  ||||||| || |  || |     |         ||  | |    ||||  | |   | |  |         |   |       |  || ||  ||    |      |
BS ATCTTCGTTAAACGCTTTTAAAACGTCAATAAAGTTTGTTCCATTTACAGGGTCGCCGCCGATTCCAACAGCTGTAGATTGCCCTACACCCGCTTCTGAGAGCTGGTGCACCGCTTCGTATGTTAATGTTCCTGAACGTGATACAACGCCTACATGACCCTTTTTATGGATGTATCCGGG
Pos                                                             <-1682148----------1682121->
R translated                                               ...T  S  I  S  c  r  R  P  Y...
```

$\Delta G^0_{min}$ = -22.42 kcal/mol

**B25.**

```
                                                                          |-----| no net gain/loss
A  TCTACTAGGTCTTCTCTTAGCCTCAGCAGGAAAATCAGCCCAATTTGGACTTCATCCGTGACTTCCATCAGCTAGTGAAGGCCCTACCCCAGTATCAGCACTCCTTCATTCCAGTACAATAGTTATAGCTGGAGTATTTACCCTCATCCGCTTTTATCCATTAATAGAAAACAACCTCAC
   ||||||||||||||||||||||||||||||||||||||||||||||||||||||||||||||||||||||| || || ||||||||||||||||||||||||||||||||||||||||||||||||||||||||||||||||||||||||||||||||||||||||||||||||||||||||||||||
R  TCTACTAGGTCTTCTCTTAGCCTCAGCAGGAAAATCAGCCCAATTTGGACTTCATCCGTGACTTCCATCAGATACTGCAGGCCCTACCCCAGTATCAGCACTCCTTCATTCCAGTACAATAGTTATAGCTGGAGTATTTACCCTCATCCGCTTTTATCCATTAATAGAAAACAACCTCAC
        |      |   |   || ||       | |  |||      ||          |     ||||||||||||||||||||||      ||       |       |   |                ||      |    |           | |   | |      |    |   || || |   |
BS AAATTTTTTCAATACTTCTACCGCAAATTCCAGAGGAGCAATTCCTGATGCAGTAACCAAATTTCATCAGATACTGCAGGCCCCTTTTCATAGAATGTTTCTCCTTTGTAATTAGGGCAGACCATATTCATATAGTCGAGGTCATTGCTTGTGTGCTTTCTCGAATTTAAGTATCCTTC
Pos                                                             <-2047836----2047857->
R translated                                               ...T  S  I  S  y  c  R  P...
```

$\Delta G^0_{min}$ = -23.61 kcal/mol

**B63.**

```
                                                                           |-----| no net gain/loss
A  ACTACTAGGTCTTCTCTTAGCCTCAGCAGGAAAATCAGCCCAATTTGGACTTCATCCGTGACTTCCATCAGCTAGTGAAGGCCCTACCCCAGTATCAGCACTCCTTCATTCCAGTACAATAGTTATAGCTGGAGTATTTACCCTCATCCGCTTTTATCCATTAATAGAAAACAACCTCAC
   |||||||||||||||||||||||||||||||||||||||||||||||||||||||||||||||||||||||| ||  | |||||||||||||||||||||||||||||||||||||||||||||||||||||||||||||||||||||||||||||||||||||||||||||||||||||||||
R  ACTACTAGGTCTTCTCTTAGCCTCAGCAGGAAAATCAGCCCAATTTGGACTTCATCCGTGACTTCCATCAGCCAGCTACGGCCCTACCCCAGTATCAGCACTCCTTCATTCCAGTACAATAGTTATAGCTGGAGTATTTACCCTCATCCGCTTTTATCCATTAATAGAAAACAACCTCAC
       ||    |                 ||||||| |       |    ||| || |         |||||||||||||||||||  |  |     |  |  |  |  |  |   |  |           |        | |     | | |      || | |  |
BS CAACCTTCATAGATGAAGTAGAAATAAAGGAAAAACCAAAGCAGAAGGAATTAAGTATACGAGTCCATCAGCCAGCTACGGCATTTTCTATAAAAGATGATACACTGTTAAAAAAATTGCTTAAAGAATCCTTAAAATGGAGTCCTTATGAAAAAGTCAAATTTTTATCGGATCAAAATG
Pos                                                              <-2414144-2414126->
R translated                                                  ...S  I  S  q  l  R  P...
```

$\Delta G^0_{min}$ = -18.72 kcal/mol

**AD47.**

```
                                                                                                                                     |----| net 1 bp inserted
A  TCAGCCCAATTTGGACTTCATCCGTGACTTCCATCAGCTAGTGAAGGCCCTACCCCAGTATCAGCACTCCTTCATTCCAGTACAATAGTTATAGCTGGAGTATTTACCCTCATCCGCTTTTATCCATTAATAG-AAAACAACCTCACTATTCAAACTTCAACACTTCTGGTTCTGG
   |||||||||||||||||||||||||||||||||||||||||||||||||||||||||||||||||||||||||||||||||||||||||||||||||||||||||||||||||||||||||||||||| ||   ||||||||||||||||||||||||||||||||||||||||||
R  TCAGCCCAATTTGGACTTCATCCGTGACTTCCATCAGCTAGTGAAGGCCCTACCCCAGTATCAGCACTCCTTCATTCCAGTACAATAGTTATAGCTGGAGTATTTACCCTCATCCGCTTTTATCCATTTATGCAAAAACAACCTCACTATTCAAACTTCAACACTTCTGGTTCTGG
    | |  | ||     |    | || | |      | | | |     |     |    |     |         | |        ||   | | |       |   |                |||||||||||||||||||  |    |     |    |         |   |  |
BS GCGGTACGATAGAAGCGGAAACCATTAACGGATTGATCGATCTCCGAGGGAAAAGCGAATCCATTGACGTTCAAAGCTTTAACGGAAATATTGCGATTAACGTAACAGAATCAGACTGCCGCTCCATTTATGCAAAAACAACGACTGGAAATGTAGAGCTTGCAATTCCGGACGAT
Pos                                                                                                                          <-3607576---3607557->
R translated                                                                                                                        m  q  K  Q  P...
```

$\Delta G^0_{min}$ = -12.63 kcal/mol

```
B1.                                                                                  |------| no net gain/loss
A  ACTACTAGGTCTTCTCTTAGCCTCAGCAGGAAAATCAGCCCAATTTGGACTTCATCCGTGACTTCCATCAGCTAGTGAAGGCCCTACCCCAGTATCAGCACTCCTTCATTCCAGTACAATAGTTATAGCTGGAGTATTTACCCTCATCCGCTTTTATCCATTAATAGAAAACAACCTCAC
   |||||||||||||||||||||||||||||||||||||||||||||||||||||||||||||||||| |||| | ||||||||||||||||||||||||||||||||||||||||||||||||||||||||||||||||||||||||||||||||||||||||||||||||||||||||||||
R  ACTACTAGGTCTTCTCTTAGCCTCAGCAGGAAAATCAGCCCAATTTGGACTTCATCCGTGACTTCCATAAGCTTGAGAAGGCCCTACCCCAGTATCAGCACTCCTTCATTCCAGTACAATAGTTATAGCTGGAGTATTTACCCTCATCCGCTTTTATCCATTAATAGAAAACAACCTCAC
   |     |         | |  | |   |  |  ||        |     |||          | ||||||||||||||||||||| |      || ||||   |     | |      |      |       |   |               |   |   ||  |  || |    || ||     |
BS ATCTTGATCCGAAAATTGACGCACGAGAAAACGATGTTGAAGAAGGAGACAAATGGACAGGCTTCCATAAGCTTGAGAAGGCCATTTGGAAGGATCAAGATATTTCCGGGGAAAAGGCAACAGCTGACCAGCTTCTCAAGGATGTAAAAGAGCTTGACGGTTCAATTCAATCACTGAAAT
Pos                                                             <-3928430------3928407->
R translated                                                  ...T  S  I  S  l  r  R  P  Y...
```

$\Delta G^0_{min}$ = -25.24 kcal/mol

```
B60.                       |-- net 168 bp loss -----------------------------------------------------------------------------------------------------|
A  GGAAAATCAGCAGGAAAATCAGCCCAATTTGGACTTCATCCGTGACTTCCATCAGCTAGTGAAGGCCCTACCCCAGTATCAGCACTCCTTCATTCCAG-(63×N)-CACTATTCAAACTTCAACACTTCTGGTTCTGGCTCTAATGTCACTAATGAAAAAATGCGTTTCTGGAGTCCAGA
   ||||||||||||||||||||||    |       |                                                                                                                     ||||||||||||||||||||||||||||
R  GGAAAATCAGCAGGAAAATCAGCTTCAGACCACTTATCC--------------------------------------------------------------------------------------------------------------------ATGAAAAAATGCGTTTCTGGAGTCCAGA
          | |  | ||||||||||||||||||||||||                                                                                                                     |||||||||| |   || |  |  |
BS ATGTCTGCGGCGGCAAAATCAGCTTCAGACCACTTATCC--------------------------------------------------------------------------------------------------------------------ATGAAAAAATTCTCTTTTCCAAGCATAG
Pos                <-4105958----------------------------------------------------------------------------------------------------------------------------4105992->
R translated  ...K  I  S  f  r  p  l  i  h                                                                                                                     E  K  M  R...
```

$\Delta G^0_{min}$ = -10.25 kcal/mol (left) / -9.92 kcal/mol (right)

```
AA5.                                                                                       |-- net 3 bp loss ----|
A  ACTACTAGGTCTTCTCTTAGCCTCAGCAGGAAAATCAGCCCAATTTGGACTTCATCCGTGACTTCCATCAGCT--AGTGAAGGCCCTACCCCAGTATCAGCACTCCTTCATTCCAGTACAATAGTTATAGCTGGAGTATTTACCCTCATCCGCTTTTATCCATTAATAGAAAACAACCTCAC
   ||||||||||||||||||||||||||||||||||||||||||||||||||||||||||||||||||||||||   ||  || |    |   |||  |||||||||||||||||||||||||||||||||||||||||||||||||||||||||||||||||||||||||||||||||||||||||
R  ACTACTAGGTCTTCTCTTAGCCTCAGCAGGAAAATCAGCCCAATTTGGACTTCATCCGTGACTTCCATCAGCTTGAG-AAAAG----AAGGCAGATTCAGCACTCCTTCATTCCAGTACAATAGTTATAGCTGGAGTATTTACCCTCATCCGCTTTTATCCATTAATAGAAAACAACCTCAC
        |     |     |    ||        ||   | ||    ||    |  | |         |    |||||||||||| |||||    ||||||||||||||||   |    ||    | |            | |       |     | ||   |    |          |     |      | |
BS GGCTATGACATTCAATGTGAAATCGTCGGAGAAGACGGCATCATCAAGCTTCCCGAGCCATCAAGCATCAGCTTGAG-AAAAG----AAGGCAGATTCAGCACTGATATTTTGATGGATTGGCAGAGACGCTTTGTCGCTGCGTATGATGTGGAAATCCAAGACTTTATTGATTCGATTCAA
Pos                                                                   <-4076055---------------------4076022->
R translated                                                       ...S  I  S  l  r   k  e     g  r  f  S  T...
```

$\Delta G^0_{min}$ = -9.49 kcal/mol (left) / -10.03 kcal/mol (right)

```
AD64.                                                                 |-- net 15 bp loss -------------------------|
A  ACTACTAGGTCTTCTCTTAGCCTCAGCAGGAAAATCAGCCCAATTTGGACTTCATCCGTGACTTCCATCAGCTAGTGAAGGCCCTACC-CCAGTATCAGCACTCCTTCATTCCAGTACAATAGTTATAGCTGGAGTATTTACCCTCATCCGCTTTTATCCATTAATAGAAAACAACCTCAC
   |||||||||||||||||||||||||||||||||||||||||||||||||| ||| |       || ||  ||| ||      || |||  |||| |||||||||||||||||||||||||||||||||||||||||||||||||||||||||||||||||||||||||||||||||||||||||||||||
R  ACTACTAGGTCTTCTCTTAGCCTCAGCAGGAAAATCAGCCCAATTTG-ACCTCAGC-------TCGAT--GCTCGT------CC-ACCTGCAGTGTCAGCACTCCTTCATTCCAGTACAATAGTTATAGCTGGAGTATTTACCCTCATCCGCTTTTATCCATTAATAGAAAACAACCTCAC
        | ||    | |     | |       ||         ||| ||||||||       |||||  ||||||      || |||||||||||||||  | ||| || | |                |  |  | |||                |    | | |          |
BS GGGCTTGGGCACGCACCAGTTCACTTGTTTTTAACAGCCATTCATTG-ACCTCAGC-------TCGAT--GCTCGT------CC-ACCTGCAGTGTCAGCAAGCGTTCTTTGCTGAGTCTGGTGAGCCTTTCTATTTTTTGAAAGTGGTTCTCAGTTCAGAATGAATCGGCTTGGCTATAT
Pos                                                 <-4178790-------------------------------------------------4178741->
R translated                                    ...I  W  T  S  a        r  c    s  s         t  c  s  v  S  T  P  S  F...
```

$\Delta G^0_{min}$ = -5.29 kcal/mol (left) / -20.87 kcal/mol (right)

-------
